# Cancer-associated snaR-A noncoding RNA interacts with core splicing machinery and disrupts processing of mRNA subpopulations

**DOI:** 10.1101/2024.07.02.601767

**Authors:** Sihang Zhou, Simon Lizarazo, Leela Mouli, Sandip Chorghade, Ruiying Cheng, K C Rajendra, Auinash Kalsotra, Kevin Van Bortle

**Author notes:** Correspondence should be addressed to K.V.B.

## Abstract

RNA polymerase III (Pol III) activity in cancer is linked to the production of small noncoding (nc)RNAs that are otherwise silent in most tissues. snaR-A (small NF90-associated RNA isoform A) - a hominid-specific ncRNA shown to enhance cell proliferation, migration, and invasion - is a cancer-emergent Pol III product that remains largely uncharacterized despite promoting growth phenotypes. Here, we applied a combination of genomic and biochemical approaches to study the biogenesis and subsequent protein interactions of snaR-A and to better understand its role as a putative driver of cancer progression. By profiling the chromatin landscapes across a multitude of primary tumor types, we show that predicted snaR-A upregulation is broadly linked with unfavorable outcomes among cancer patients. At the molecular level, we unexpectedly discover widespread interactions between snaR-A and mRNA splicing factors, including SF3B2 - a core component of the U2 small nuclear ribonucleoprotein (snRNP). We find that SF3B2 levels are sensitive to high snaR-A abundance and that depletion of snaR-A alone is sufficient to decrease intron retention levels across subpopulations of mRNA enriched for U2 snRNP occupancy. snaR-A sensitive genes are characterized by high GC content, close spatial proximity to nuclear bodies concentrated in pre-mRNA splicing factors, and functional enrichment for proteins involved in deacetylation and autophagy. We highlight examples of splicing misregulation and increased protein levels following snaR-A depletion for a wide-ranging set of factors, suggesting snaR-A-driven splicing defects may have far-reaching effects that re-shape the cellular proteome. These findings clarify the molecular activities and consequences of snaR-A in cancer, and altogether establish a novel mechanism through which Pol III overactivity may promote tumorigenesis.

## Introduction

The Pol III transcriptome includes multiple classes of small, highly structured noncoding RNA species involved in diverse cellular processes. For example, tRNA and 5S, the smallest ribosomal RNA (rRNA), play integral roles in translation and protein accumulation. Other Pol III-transcribed genes encode ncRNAs involved in splicing (U6, U6atac), transcription regulation (7SK), RNA processing and stability (H1, RMRP, Y RNA), as well as autophagy (Vault). The signal recognition particle RNA (7SL), on the other hand, targets secretory proteins for translation and appropriate folding at the endoplasmic reticulum. In this way, Pol III transcription produces a multitude of RNAs with core housekeeping functions^1–3^.

In humans, retrotransposition of the highly expressed 7SL RNA has established hundreds of 7SL pseudogenes, which are thought to be transcriptionally and functionally incompetent^4^. Through subsequent duplications and sequence divergence, Alus - the most abundant class of Short Interspersed Nuclear Elements (SINEs) - evolved and further proliferated^5^. Two hominid-specific Pol III-transcribed genes, BC200 and snaR, are evolutionarily derived from Alu sequences. The expression of BC200 and snaR are restricted to specific contexts, with BC200 expressed in neurons and snaR most highly expressed in testis^6,7^. Despite evidence that both ncRNAs also re-emerge in human tumors, functional characterization of BC200 and snaR in tissue- and cancer-specific contexts remains lacking.

snaR - an acronym for <u>s</u>mall <u>N</u>F90-<u>a</u>ssociated <u>R</u>NA - was first reported in 2007 as an RNA species upregulated in EBV-infected B cells and enriched in biochemical pulldown experiments targeting an RNA-binding protein, ILF3^7,8^. Multiple isoforms for snaR, ranging from snaR-A to snaR-I, are encoded in multiple clusters of repetitive sequence elements, with snaR-A the dominant isoform expressed in both testis and cancers^7,9^. Recent reports indicate that snaR-A promotes cancer cell migration and invasion, as well as proliferation, in multiple contexts^10–17^. Thus, deconstructing the molecular function and/or consequence of snaR-A emergence remains critical for understanding how snaR-A contributes to cancer initiation and progression.

Here, we applied genomic and biochemical approaches to advance our molecular understanding of snaR-A and its significance in cancer. We discover that snaR-A interacts with core splicing proteins and promotes processing defects in mRNA subpopulations that are characterized by multiple features, including U2 residency, nuclear speckle proximity, and hallmarks of low splicing efficiency. snaR-A-driven perturbations reduce protein abundance for a wide-ranging set of factors, suggesting the emergence of snaR-A in cancer leads to disruption of cellular homeostasis via its role as a molecular disruptor of splicing. Together, these findings expand our understanding of the nature of snaR-A activity and provide new insights into the link between Pol III overactivity, cancer initiation, and disease progression.

## Results

### A chromatin-based survey links snaR-A to unfavorable outcomes in cancer

snaR-A is overlooked in most genomic studies, in part due to its small (∼120 nt), highly structured nature and absence in Z-traditional RNA-seq experiments, its omission from certain gene annotation sets, and its repetitive multi-copy gene origins^9,18^. As a consequence, a full survey of snaR-A expression across cancer subtypes remains largely lacking despite evidence of upregulation in multiple contexts^7,10–14,17^, and the molecular target(s) of the simple, thermodynamically stable stem-loop structure of snaR-A remain similarly uncharacterized (**Figure 1a-b**). We therefore devised a chromatin-based framework for uniformly defining the expression “state” of *SNAR-A* genes using currently available resources. Gene accessibility, which correlates with the occupancy of RNA polymerase III and transcription, presents an effective method for predicting the activity of Pol III-transcribed genes, circumventing the otherwise unreliable interpretation of transcription from steady-state RNA levels^15^. Using this approach, we annotated the “ON” and “OFF” states of *SNAR-A* and other Pol III-transcribed genes using chromatin accessibility defined by ATAC-seq across > 400 human tumors^19^.

**Figure 1.**
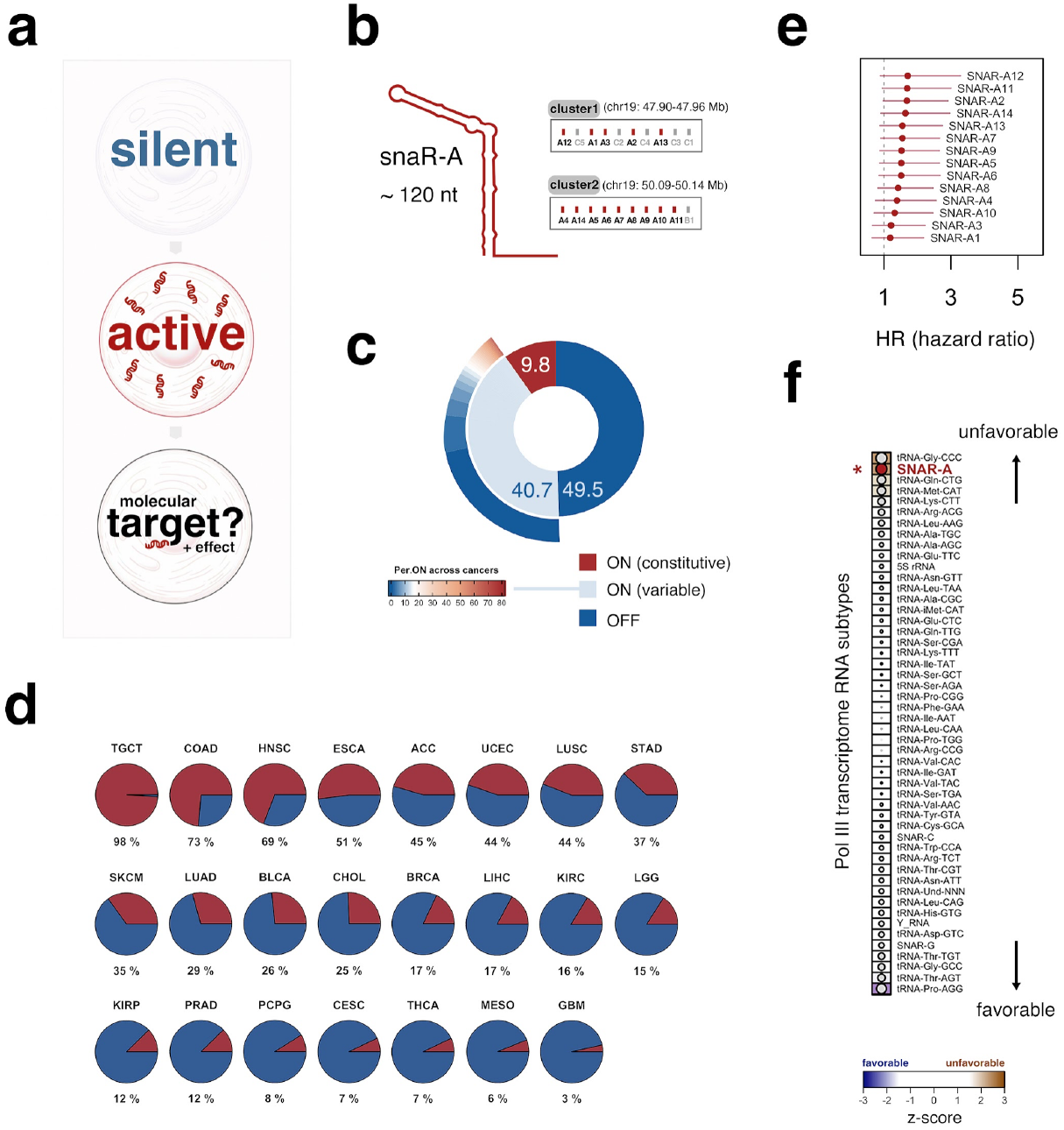
Active Pol III signatures at snaR-A genes are associated with unfavorable outcomes in cancer. **(a)** Illustrative overview of the dynamic, context-restricted activity of snaR-A and the present study’s aim to uncover its molecular significance. **(b)** RNAfold secondary structure prediction for snaR-A ncRNA (left), which is derived from two multi-copy gene clusters on chromosome 19 (right). *SNAR-A* genes (1-14) highlighted in red. **(c)** Percentage of annotated ncRNA genes (< 1 Kb, n = 20,133) predicted to be constitutively ON (active), constitutively OFF (inactive), or to have variable activity (e.g. *SNAR-A*) across 23 cancer subtypes. **(d)** High-to-low cancer-specific frequencies of predicted *SNAR-A* gene ON (red) and OFF (blue) state. **(e)** Hazard ratios and corresponding 95% confidence interval for all *SNAR-A* genes (Cox proportional hazard regression model). **(f)** High-to-low ranking of Pol III-transcribed gene “types” based on predicted positive / negative roles in cancer. score reflects the observed median Wald statistic for a group “type” compared against a randomization-based null distribution.

Nearly half of the annotated ncRNA genes < 1 Kb are silent across all 23 cancer subtypes, whereas a small subset (9.8%) is constitutively active across diverse cancers. *SNAR-A*, on the other hand, is included among the remaining genes encoding small ncRNAs characterized by a variable ON/OFF state across tumor subtypes (**Figure 1c**). We note that genes encoding additional isoforms of snaR, including snaR-B and snaR-C, are dispersed within the same repetitive clusters of *SNAR-A* genes on chromosome 19 but are characterized by divergent sequence features and are less often expressed in tissues or cancer^7,9^ (**Figure 1b**, Supplemental Figure 1).

*SNAR-A* expression, defined using our binary ON/OFF approach, is most frequently observed in Testicular Germ Cell Tumors (TGCT), an observation that is consistent with high snaR-A abundance reported in human testis^7,9^. *SNAR-A* is otherwise commonly active across > 50 % of Colon adenocarcinoma (COAD, 73%), Head and Neck squamous cell carcinoma (HNSC, 69%), and Esophageal carcinoma (ESCA, 51%), and most frequently silent in Glioblastoma multiforme (GBM, 3%), Mesothelioma (MESO, 6%), and Thyroid carcinoma (THCA, 7%).

To explore the potential impact of emergent *SNAR-A* expression on patient outcomes, we applied a Cox proportional hazard regression for *SNAR-A* gene state across cancer, restricting for cancer types with variable *SNAR-A* expression^20^. Briefly, cancers with predicted uniform ON and uniform OFF states, defined using the median statistic across all 14 *SNAR-A* genes, were excluded from further analysis. Cancer subtypes with variable expression were subsequently restricted to tumor samples with confident ON and confident OFF states, which were thereafter used as input for survival analysis. Using this approach, we find that snaR-A activity is a predicted negative factor in cancer, with a median hazard ratio of 1.52 (**Figure 1e**, n = 187, HR range 1.19 (snaR-A1) to 1.71 (snaR-A12)). *SNAR-A* genes are also characterized by a comparatively high median Wald statistic in comparison to Pol III-transcribed gene types and other small ncRNA genes < 400 bp (**Figure 1e-f**, z = 2.46). These findings indicate a link between context-specific *SNAR-A* expression and unfavorable outcomes in patients, motivating further exploration of the molecular targets and consequences of snaR-A expression.

### Integrated proteomic and genomic discovery of snaR-A RNA-protein interactions

Most Pol III-derived small ncRNAs were discovered as core RNA components of specific ribonucleoprotein complexes (RNPs), typically central to the established roles of each respective ncRNA^21^. Similarly, the origin story of snaR-A includes its discovery as an RNA molecule enriched in biochemical pulldown experiments for NF90, a specific isoform of the RNA-binding protein, ILF3^7^. However, the functional significance of snaR-NF90 interactions and whether additional, presently uncharacterized snaR-A-binding factors are more centrally involved in its potential role remains unknown. We, therefore, sought to discover snaR-A RNA-protein interactions *de novo*, using 5’ biotinylated snaR-A to pull down interacting proteins from THP-1 cell lysates followed by mass spectrometry (MS) (**Figure 2a**). In total, our *in vitro* pull-down assay yielded 375 protein hits with 61 exhibiting significant enrichment in snaR-A pull downs compared to both empty bead and scramble RNA control experiments (**Figure 2a**, Supplemental Table 1). Most of the 61 snaR-A-protein interactions identified correspond to a multitude of RNA-binding proteins (RBPs), with La (SSB), an RNA-chaperone protein important for processing and stability of Pol III-transcribed genes, the most confident hit (**Figure 2a-b**)^22^. ILF3 (NF90) is also notable among the list of significantly enriched proteins, indicative of snaR-A-NF90 interactions in THP-1 monocytes (**Figure 2b**). We confirm snaR-A-La and snaR-A-ILF3 interactions through reciprocal pulldown experiments, in which flag-tagged La and ILF3 isoforms and associated RNA are captured by RNA immunoprecipitation (RIP) (**Figure 2c**). snaR-A RNA is enriched in pull-down experiments for either NF90 or NF110 isoforms, suggesting snaR-A-ILF3 interactions are most likely isoform-independent (**Figure 2c**).

**Figure 2.**
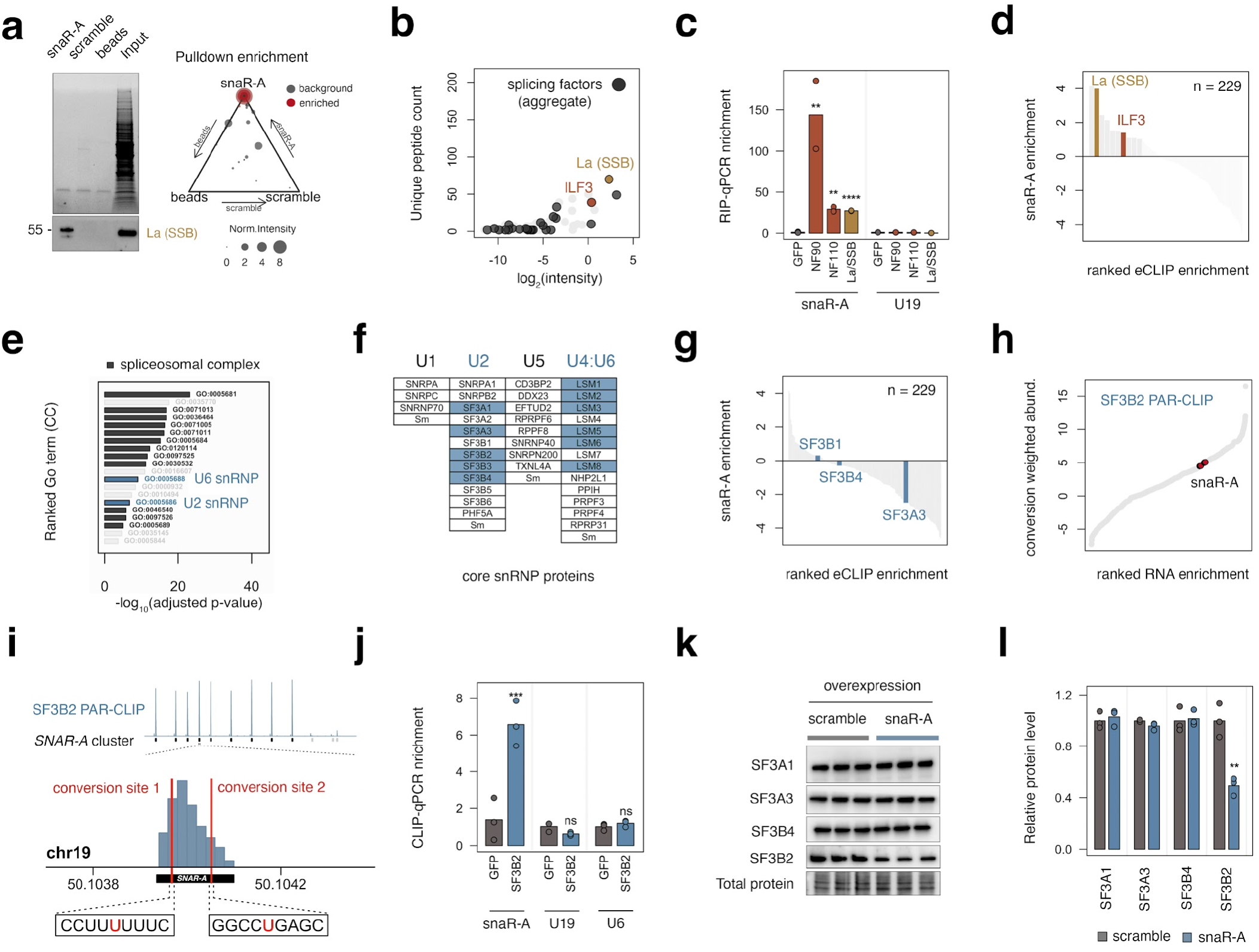
snaR-A ncRNA interacts with mRNA splicing machinery, including U2 snRNP protein SF3B2. **(a)** Immunoblot showing total protein (top) and RNA chaperone protein La (bottom) in snaR-A, scramble, or empty beads pull-down experiments. Triangle plot depicts normalized intensity of proteins enriched for snaR-A binding (red) and non-enriched proteins (gray). **(b)** Normalized intensity and sum of unique peptides across pull-down replicates. ILF3, La, and splicing-associated proteins are highlighted. **(c)** RNA Immuno-precipitation (RIP)-qPCR quantification of target RNA enrichment (snaR-A, U19) in Flag-NF90, Flag-NF110, and Flag-La (SSB) RIP experiments compared to Flag-GFP, relative to the enrichment of control small ncRNA, Z30. **(d)** Bar plot visualization of the maximum snaR-A enrichment score across > 200 eCLIP datasets; ILF3 and La are highlighted. **(e)** Gene Ontology Cellular Component (GO:CC) enrichment analysis for significantly enriched proteins in snaR-A RNA pull-down experiments. **(f)** Visualization of the core snRNP proteins of U1, U2, U5, and U4/U6, with all identified snaR-A interacting proteins highlighted in blue. **(g)** Analogous eCLIP survey barplot, highlighting snaR-A enrichment in experiments for U2 snRNP proteins SF3A3, SF3B1, and SF3B4. **(h)** snaR-A enrichment, weighed by conversion-based evidence of UV cross-linking, in SF3B2 PAR-CLIP experiments. **(i)** Visualization of SF3B2 PAR-CLIP read pileup across snaR-A gene clusters, and inspection of sequence conversion sites indicative of direct RNA-protein cross-linking sites. **(j)** Crosslinking immunoprecipitation (CLIP)-qPCR quantification of target RNA enrichment (snaR-A, U19, U6 snRNA) in Flag-SF3B2 CLIP experiments compared to Flag-GFP, relative to the enrichment of control small ncRNA, Z30. **(k-l)** Immunoblot (k) and corresponding quantification (l) of U2 snRNP proteins SF3A1, SF3B1, SF3B4, and SF3B2 in HEK293T cells overexpressing snaR-A.

To complement our proteomics result, we further conducted a large-scale survey for snaR-A enrichment in genomic RNA cross-linking and immunoprecipitation (eCLIP-seq) experiments available through ENCODE^23^. Using this approach, which integrates the RNA-binding patterns for > 150 unique RBPs (Supplemental Table 2), we find that both La and ILF3 eCLIP experiments are among the highest with respect to snaR-A enrichment (**Figure 2d**, Supplemental Figure 2), further supporting these interactions and the validity of our *in vitro* binding assay.

### snaR-A interacts with mRNA splicing machinery, including U2 snRNP protein SF3B2

Beyond uncovering interactions with chaperone protein La and reaffirming ILF3-snaR-A binding, our proteomic analysis unexpectedly pointed to enrichment for a multitude of splicing factors (**Figure 2b**). Gene ontology (GO) enrichment analysis confirms overrepresentation of numerous spliceosomal complex-related terms and, more specifically, U2 and U6 snRNP proteins (**Figure 2e**). Among the core U1, U2, U5, and U4/U6 snRNP proteins, snaR-A interactions were indeed specific to U2 and U4/U6 components (**Figure 2f**). Revisiting the large-scale eCLIP survey, we find that experiments for SF3B1, SF3B4, and SF3A3 - components of the U2 snRNP complex - are not enriched for snaR-A ncRNA, limiting the likelihood of direct interactions between snaR-A and these particular proteins (**Figure 2g**).

In contrast, analysis of PAR-CLIP (<u>P</u>hoto <u>A</u>ctivatable <u>R</u>ibonucleoside-enhanced) experimental data for SF3B2, another U2 snRNP protein, captures nucleotide-level evidence of direct snaR-A-SF3B2 interactions (**Figure 2h-i**)^24^. We confirm snaR-A-SF3B2 interactions by CLIP-qPCR and further show that overexpression of snaR-A results in subunit-specific depletion of SF3B2, in contrast to U2 snRNP proteins SF3A1, SF3A3, and SF3B4 (**Figure 2j-l**). These data, which altogether suggest that snaR-A interactions may selectively disrupt specific snRNP proteins, prompted us to investigate the downstream effects of snaR-A abundance on splicing outcomes.

### snaR-A depletion reduces intron retention in specific mRNA subpopulations enriched for U2-binding

To determine the consequences of snaR-A expression on splicing, we sought to deplete snaR-A and explore the effect of reduced snaR-A abundance on mRNA processing through deep RNA sequencing. We reduced snaR-A using a series of siRNAs designed against 3’-proximal sequences, which achieved maximal reduction compared to siRNAs targeting other regions within the small stem-loop structure of snaR-A (Supplemental Figure 3). Splicing analysis uncovers reduced intron retention (IR, an indicator of unprocessed mRNA) following snaR-A depletion, suggesting the presence of snaR-A may functionally perturb efficient splicing. We note that this pattern is independently observed using either IRFinder^25^, a tool designed to quantify intron retention levels (**Figure 3a**), or rMATS^26^, a tool that more broadly queries multiple forms of differential splicing events (**Figure 3b**). PCR analysis confirms dynamic intron retention levels for several rMATS-defined differential splicing events following snaR-A depletion (Supplemental Figure 4).

**Figure 3.**
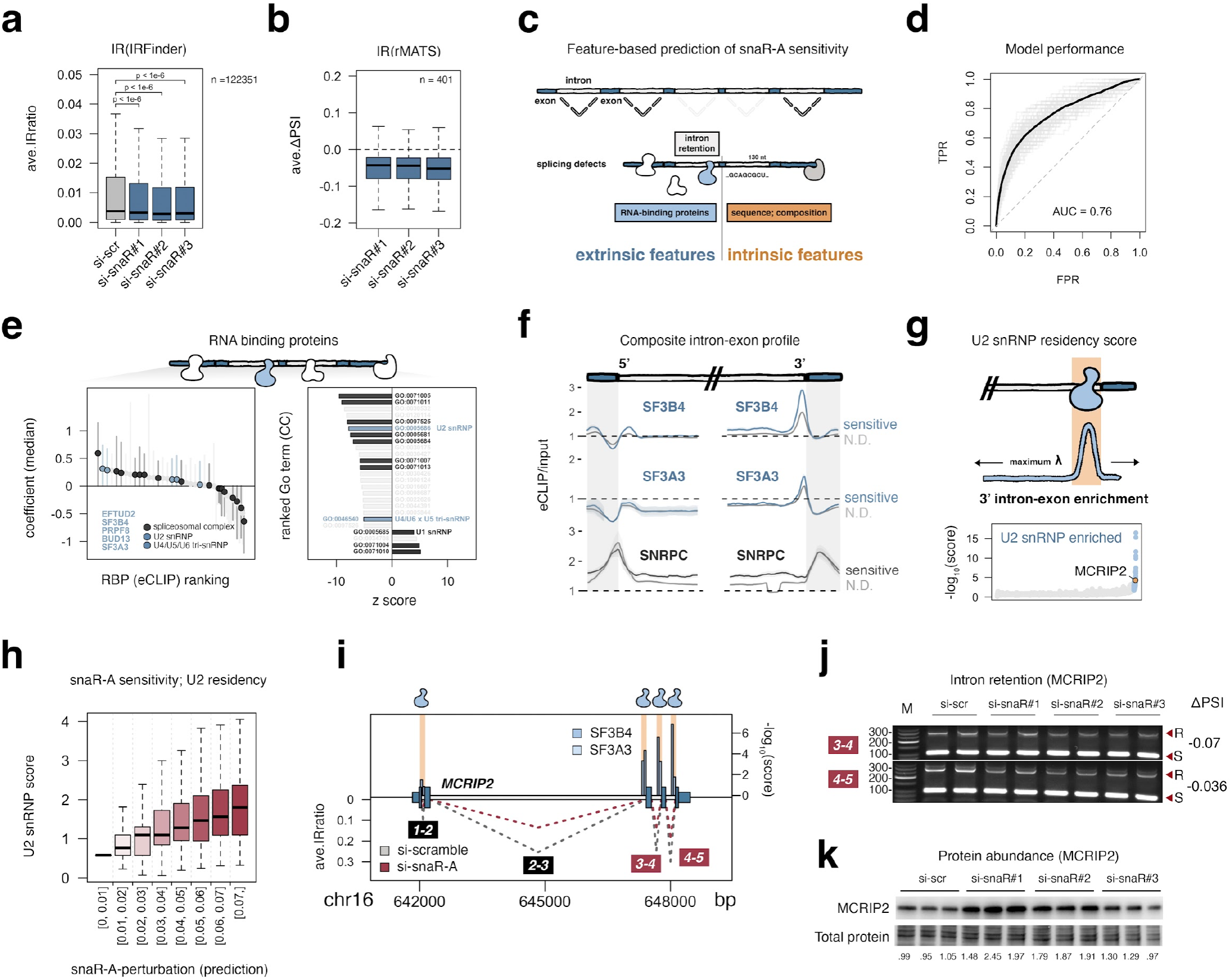
Depletion of snaR-A reduces intron retention in mRNA subpopulations enriched for U2 snRNP residency, linking snaR-A expression with splicing defects. **(a)** Distribution of intron retention levels in control (si-scramble) and snaR-A siRNA-1/2/3 experiments, determined via IRFinder across 122,351 introns. ave.IRratio = average Intron Retention ratio. **(b)** Distribution of delta PSI (percent spliced-in) levels following snaR-A depletion, determined via rMATS across 401 introns. **(c)** Illustration of feature prediction framework for profiling snaR-A-related mRNA splicing sensitivity. Logistic regression model combines extrinsic (e.g. RBP-binding) and intrinsic features (e.g. GC content). **(d)** ROC curve depicts model performance for predicting snaR-A sensitivity. **(e)** Analysis of regression coefficients across all RBPs (left), and mRNA population snaR-A sensitivity scores for RBPs grouped by GO:CC term (right). **(f)** Composite eCLIP signal pileup at 5’- and 3’-intron-exon boundaries for U2 snRNP proteins SF3B4 and SF3A3, and U1 snRNP protein SNRPC, across snaR-A sensitive and snaR-A insensitive mRNA populations. **(g)** Illustration of U2 snRNP residency scoring framework. Signal within 50 nt upstream of each 3’-intron-exon boundary is compared to a maximum expectation - lambda - of the signal upstream, downstream, or across all analogous junctions. **(h)** Distributions of U2 snRNP residency scores over mRNA populations as a function of snaR-A sensitivity prediction. **(i)** Transcript structure, U2-binding, and intron retention of MCRIP2, a snaR-A-sensitive transcript. **(j)** PCR analysis of MCRIP2 intron retention levels corresponding to exon-exon junctions 3-4 and 4-5. **(k)** Immunoblot analysis of MCRIP2 protein levels following snaR-A depletion.

**Figure 4.**
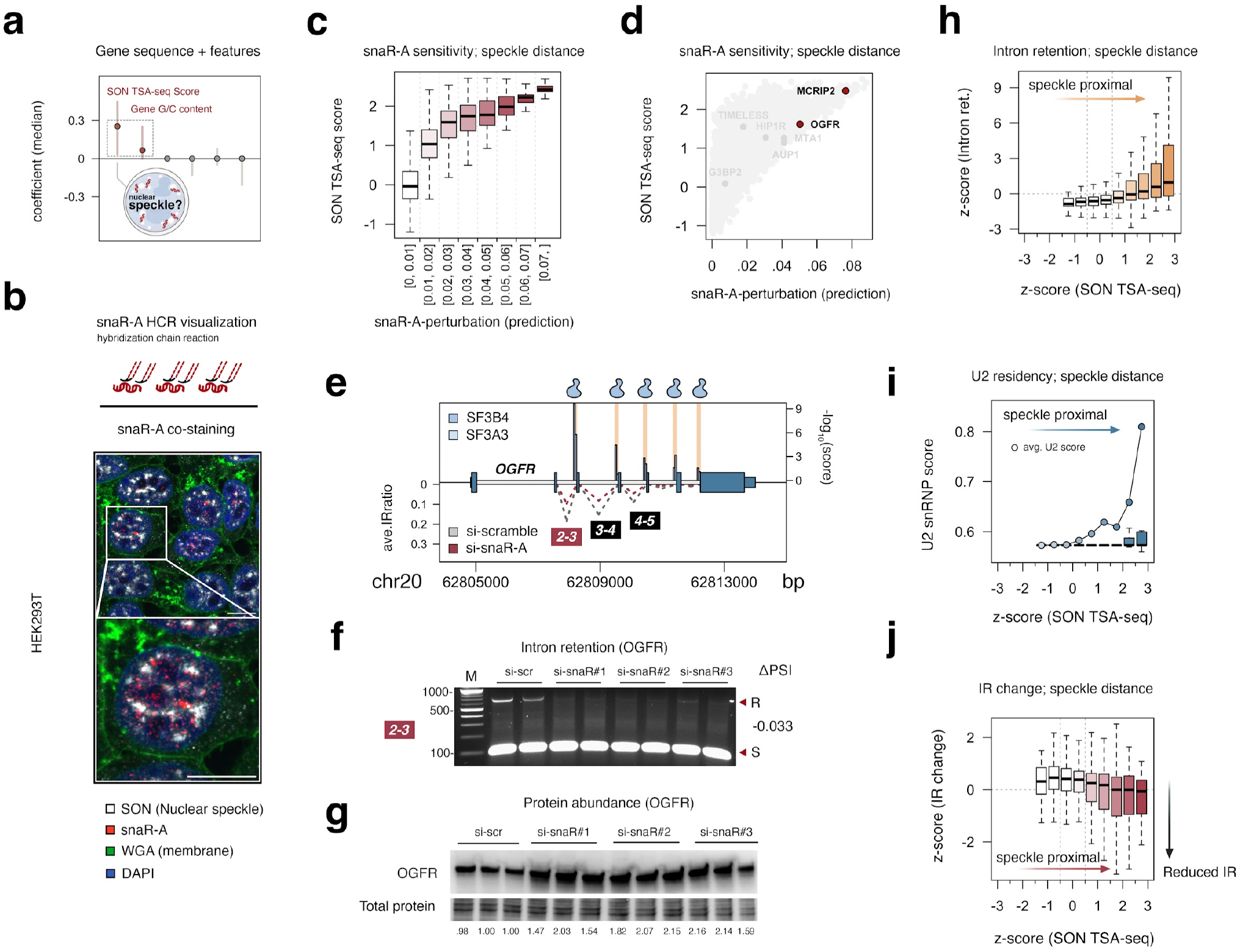
Nuclear speckle proximity is linked to higher intron retention, U2 residency, and snaR-A-related mRNA splicing sensitivity. **(a)** Analysis of regression coefficients (related to Figure 3E), restricted to nuclear speckle proximity (SON-TSA-seq) and intrinsic features (i.e. sequence composition) **(b)** Hybridization chain reaction-fluorescence in situ hybridization (HCR-FISH) targeting snaR-A, co-stained for SON (nuclear speckles), wheat germ agglutinin (WGA; membrane), and DAPI. **(c)** Distribution of SON-TSA-seq scores for genes binned by predicted snaR-A sensitivity. **(d)** Individual gene SON-TSA-seq scores as a function of predicted snaR-A sensitivity - with gene examples highlighted. **(e)** Transcript structure, U2-binding, and intron retention of OGFR, a nuclear speckle-proximal snaR-A-sensitive transcript. **(f)** PCR analysis of OGFR intron retention levels corresponding to exon-exon junction 2-3. Arrows indicate retained (R) and spliced (S) introns. **(g)** Immunoblot analysis of OGFR protein levels following snaR-A depletion.**(d)** Gene intron retention (IR) level scores binned by nuclear speckle proximity (SON-TSA-seq). **(h-j)** Distributions of intron retention levels (h), U2 snRNP residency scores (i), and dynamic transcript-wide IR scores (j) following snaR-A depletion as a function of nuclear speckle distance (SON TSA-seq score). Higher scores reflect closer spatial proximity to nuclear speckles.

Given that snaR-A depletion appears to broadly reduce IR levels, we next considered whether specific mRNA sub-populations were characterized by particularly significant changes in intron splicing following snaR-A disruption. To this end, we computed a transcript-centric dynamic IR score that compares the distribution of differential intron ratios following si-snaR-A across all annotated introns for each gene (Supplemental Figure 5). Using this approach, we identify 136 genes with significant differences in IR levels when accounting for all introns (adjusted p-value < 0.05). By further integrating a series of extrinsic (e.g. RBP-binding) and intrinsic attributes (e.g. GC content) inherent to each transcript, we applied machine learning to assess the predictive power of specific features for identifying snaR-A sensitivity (**Figure 3c**). Using a penalized logistic regression model to discriminate snaR-A sensitive transcripts (AUC = 0.76, **Figure 3d**), we find that several U2 and U4/U6 snRNP proteins are positive predictors of snaR sensitivity, includeing the core U2 snRNP protein SF3B4 (**Figure 3e**). This result is consistent with a complementary survey of snaR-A sensitivity scores across RNA populations bound by proteins ascribed to specific GO terms, such that U2 snRNP and related splicing-associated terms are linked to reduced IR levels in response to snaR-A depletion (**Figure 3e**). In contrast to U2, specific U1 snRNP features are inversely associated with snaR-A-related splicing defects, indicating some level of snRNP specificity (**Figure 3e**, Supplemental Figure 6). These data altogether point to a functional link between snaR-A, U2 snRNP interactions, and disruption of efficient splicing. We, therefore, sought to further investigate the relationship between U2 snRNP residency and snaR-A-related perturbations.

**Figure 5.**
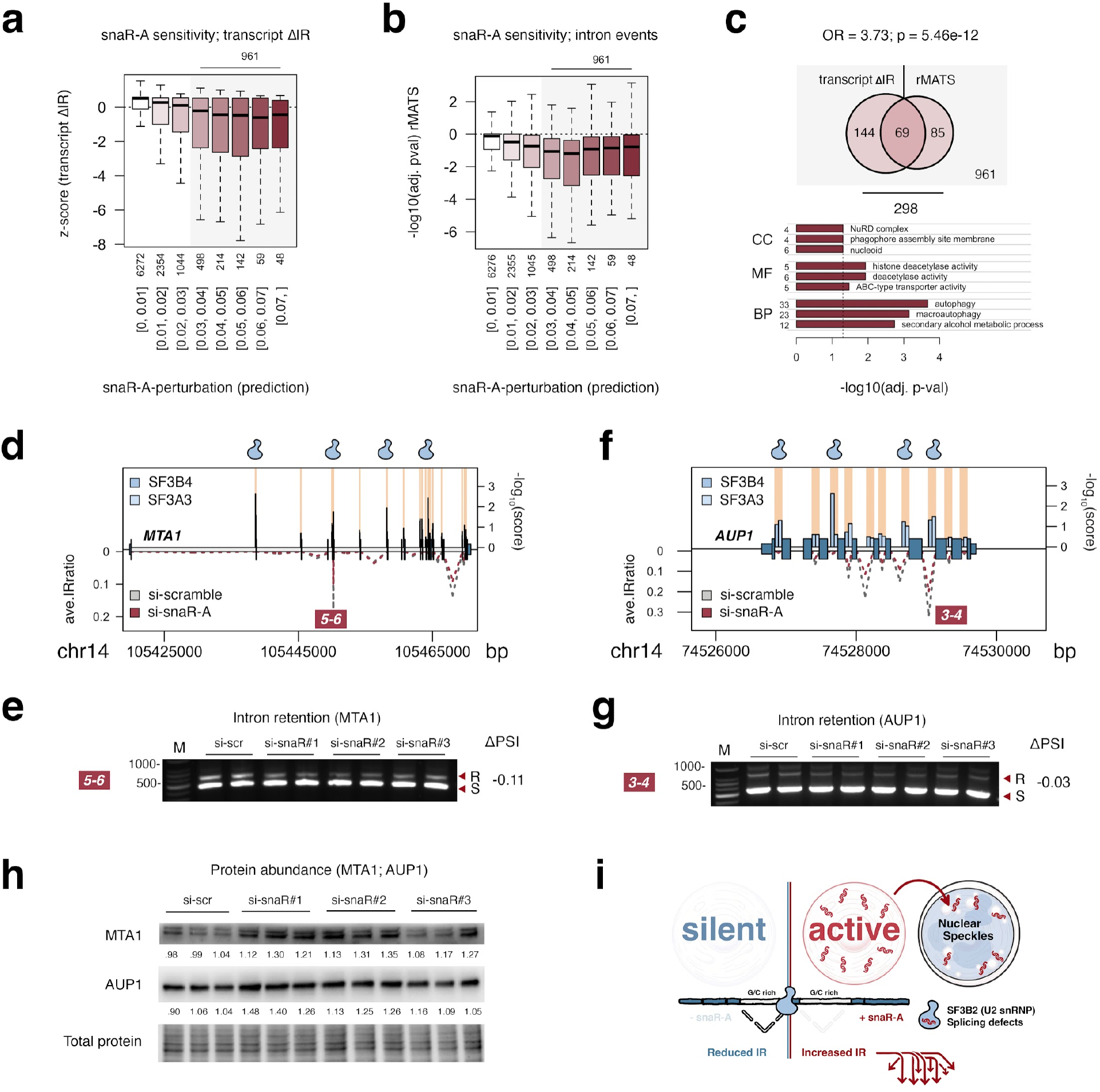
snaR-A-related splicing defects are overrepresented for genes encoding histone deacetylases and factors involved in cell autophagy. **(a-b)** Distributions of transcript-wide (a) and intron-centric (b) differential IR scores following snaR-A depletion, binned by model prediction of snaR-A sensitivity. **(c)** Gene ontology (GO) enrichment analysis for genes with significant transcript-wide (z < -3) and/or intron-centric (padj < 0.01) scores and predicted snaR-A sensitivity. Enriched GO terms for Cellular component (CC), Molecular function (MF), and Biological process (BP) are shown (padj < 0.05). **(d)** Transcript structure, U2-binding, and intron retention levels related to MTA1, a NuRD complex subunit. **(e)** PCR analysis of MTA1 intron retention levels corresponding to exon-exon junctions 5-6. **(f-g)** Analogous transcript profile (f) and PCR analysis of differential intron retention levels corresponding to exons 3-4 (g) for AUP1, an autophagy related protein. **(h)** Immunoblot analysis of MTA1 and AUP1 protein levels following snaR-A depletion. **(g)** Model: through interactions with U2 snRNP proteins, particularly SF3B2, snaR-A emergence drives intron retention in mRNA subpopulations. snaR-A localizes to nuclear foci, and its disruption of splicing is particularly notable for sub-populations of nuclear speckle-associated transcripts. The disruption of splicing reduces protein levels for a wide-ranging set of factors, like to have far-reaching consequences (arrows).

### U2 snRNP 3’-intron-exon residency levels predict snaR-A-related splicing defects

eCLIP experiments are capable of capturing signal enrichment for U1 and U2 snRNP proteins at the 5’ exon-intron and 3’ intron-exon junctions of unspliced introns, respectively, in accordance with their roles in splicing^27^. We note that eCLIP signal pileup for U2 snRNP proteins SF3B4 and SF3A3, enriched at the 3’ intron-exon junction, is visually higher across snaR-A sensitive mRNAs, consistent with our logistic regression model (**Figure 3f**). This pattern contrasts that of U1 snRNP protein SNRPC, which is not linked with snaR-A sensitive mRNAs (**Figure 3e-f**).

To quantitatively score U2 snRNP residency at 3’ intron junctions, we compared the signal spanning a 50 nt interval upstream of each intron-exon junction to an expectation (lambda), computed as the maximum signal upstream, downstream, or globally across all intervals (**Figure 3g**). A Poisson probability was then computed for each intron-exon eCLIP pileup (observation) and transformed into a U2 “residency” score. We show that U2 snRNP residency scores determined using this approach are linked to the predicted snaR-A sensitivity metrics and help identify numerous transcripts with significant U2 residency (**Figure 3h**). For example, *MCRIP2*, which is characterized by significant differential IR events by rMATS (**Figure 3b**), is also includeed among a list of targets with significant U2 residency (**Figure 3g**, adjusted pval < 0.05). Visualization of *MCRIP2* splicing and U2 residency features illustrates the link between U2 signal and intron retention, which we validated by PCR (**Figure 3i-j**). Depletion of snaR-A reduces *MCRIP2* IR levels while conversely increasing MCRIP2 protein levels, suggesting efficient processing leads to increased translation of fully mature mRNA (**Figure 3k**).

### snaR-A and downstream splicing defects are associated with nuclear splicing bodies

In addition to RBPs, our logistic regression model included specific gene, transcript, and intron-related features, such as nuclear position, G/C content, and intron length. G/C content is a consistent positive predictor of snaR-A splicing sensitivity at the gene-, intron-, and splice junction levels, whereas intron length is a negative predictor, suggesting transcripts with high GC content and short introns may be more susceptible to snaR-A (**Figure 4a**, Supplemental Figure 6). However, we find that spatial proximity to nuclear speckles, measured by SON TSA-seq, is an even stronger positive predictor of snaR-A perturbation (**Figure 4a**). Speckles are subnuclear bodies enriched for splicing factors and higher levels of co-transcriptional splicing^28–31^. Using an amplification-based *in situ* FISH approach, we discover that snaR-A is primarily localized within nuclear foci that are adjacent to bodies stained for SON, a nuclear speckle protein^32,33^, suggesting snaR-A likely perturbs splicing at or in close spatial proximity to nuclear speckles (**Figure 4b**).

SON TSA-seq captures nuclear speckle-proximity by distant-dependent tyramide signal amplification (TSA) reactions directed towards sites recognized by SON primary antibodies^34^. We specifically surveyed all available SON TSA-seq experiments (https://www.4dnucleome.org/) and integrated these data into a summary score, such that higher scores reflect closer spatial proximity to nuclear speckles according to previous studies. As shown for U2 residency, SON TSA-seq scores are strongly linked to predicted snaR-A splicing sensitivity (**Figure 4c-d**). Revisiting *MCRIP2*, we find this gene is characterized by particularly high SON TSA-seq scores, suggesting *MCRIP2*-related IR is associated with both high U2 3’-intron-exon residency and speckle proximity (**Figure 4d**). Investigation of other genes with strong SON TSA-seq scores, such as *OGFR*, confirms similar examples of reduced intron retention and increased protein abundance following snaR-A depletion (**Figure 4e-g**). The improved splicing efficiency of *OGFR* and upregulation of Opioid growth factor receptor protein levels may be particularly noteworthy in the context of snaR-A and cancer, given that OGFR functions as an inhibitor of cell proliferation and tumorigenesis^35^.

### Speckle-associated genes are broadly enriched for intron retention and higher U2 residency levels

In parallel to our snaR-A-centered analyses, which identified U2 residency and speckle proximity as key predictors of enhanced splicing following snaR-A depletion, we also asked whether IR signatures, U2 patterns, and splicing sensitivity scores were broadly associated, either positively or negatively, with nuclear speckle proximity. We integrated our transcript-centric IR analysis with SON TSA-seq, for example, to understand how IR levels correlate with spatial proximity to speckles. Surprisingly, we show that transcript-centric IR levels are positively associated with nuclear speckle proximity, such that speckle-associated features exhibit significantly higher intron retention signatures, in contrast to those that are speckle distant (**Figure 4h**). Although conceptually these results run counter to patterns of higher splicing efficiency reported at nuclear speckles, they are otherwise consistent with evidence that specific, inefficiently spliced mRNA subpopulations are stably enriched in nuclear speckles^31,36,37^.

We similarly find that features with high SON TSA-seq scores are characterized by higher U2 residency, suggesting mRNAs produced by speckle-proximal genes are more likely to be unspliced and bound by U2 snRNP (**Figure 4i**). Although TSA-seq scores are also generally linked with reduced intron retention follow snaR-A depletion, most speckle-proximal genes - taken as a distribution - are not characterized by significant differential IR levels (**Figure 4j**). These results suggest snaR-A depletion effects may be pronounced for a smaller subset of speckle-related genes.

### snaR-A-driven splicing defects are enriched for factors involved in deacetylation and autophagy

While our global model predictions of snaR-A sensitivity generally track with the transcript-wide and intron-centric differential IR scores observed following snaR-A depletion, even the strongest prediction scores include genes that lack evidence of differential IR events (**Figure 5a-b**). These trends are consistent with the mild differences in IR observed for most speckle-proximal genes (**Figure 4j**), again suggesting some level of selectivity in snaR-A-driven splicing perturbation. These findings prompted us to reexamine the mRNA subpopulations with particularly strong transcript-wide and/or intron-centric IR dynamics. We restricted our analysis to transcripts with significant IR reduction signatures, as well as relatively high prediction scores, indicating consistency with the overall patterns of snaR-A-related splicing defects. We note that transcripts with significant intron events, determined by rMATS, significantly overlap the cohort of genes with significant transcript-wide IR dynamics (p = 5.46e-12, Fisher’s exact test, **Figure 5c**).

To understand the functional significance of snaR-A splicing perturbation, we performed gene ontology (GO) enrichment analysis on the subpopulation of genes with notable splicing defects and prediction scores (n = 298). Among the significantly overrepresented GO terms, we find that the NuRD complex (Nucleosome Remodeling and Deacetylase) and histone deacetylase activity are the top cellular component (CC) and molecular function (MF) annotations, respectively (**Figure 5c**). Splicing defects are otherwise significantly enriched for factors involved in autophagy - the biological process (BP) of self-degradation (**Figure 5c**). These findings are supported by examination of MTA1 - a central regulatory subunit of NuRD - and AUP1 - a key regulator of lipid metabolism and autophagy. Both MTA1 and AUP1 transcripts are characterized by significant U2 residency at 3’ intron-exon boundaries corresponding to introns with downregulated IR following snaR-A depletion (**Figure 5d-g**). We show that, like other transcripts with reduced IR, both MTA1 and AUP1 protein levels increase following snaR-A depletion, suggesting snaR-A-related splicing defects influence the cellular abundance of proteins related to chromatin remodeling and autophagy (**Figure 5h**).

## Discussion

Nearly 20 years following the discovery of snaR-A by two contemporaneous studies in 2007, its molecular activities and the consequences arising from its expression in cancer have remained poorly defined. Here, our survey of snaR-A protein interactions and ensuing functional genomic study provides evidence that snaR-A disrupts mRNA processing through interactions with splicing machinery, most likely through U2 snRNP protein SF3B2. This model is supported by *in vitro* pull-down experiments, signatures of direct snaR-A RNA-protein cross-linking in SF3B2 PAR-CLIP experiments, and reduced intron retention following snaR-A depletion for transcripts with notable U2 snRNP occupancy. In addition, SF3B2 protein levels are uniquely sensitive to snaR-A overexpression, in contrast to other snaR-A-binding proteins, including ILF3 isoforms NF90 and NF110 (Supplemental Figure 7) or other U2 snRNP proteins tested (**Figure 2k**). We also note that direct visualization of snaR-A revealed nuclear foci in proximity to nuclear speckles (**Figure 4b**), indicating that snaR-A-driven splicing perturbation likely takes place at or near sites of mRNA processing, rather than through indirect mechanisms such as cytoplasmic sequestration of splicing machinery. Together, these findings give rise to a model in which snaR-A expression drives processing defects in specific mRNA subpopulations through direct interactions with splicing machinery in the nucleus (**Figure 5g**).

Structurally, SF3B2 is one of several proteins that comprise the SF3b subcomplex, a core component of the U2 snRNP with important roles in spliceosome assembly and stability, branch point recognition, and splicing catalysis^38^. Within SF3b, SF3B2 interactions are largely peripheral, such that SF3B2 may primarily contribute to regulatory and stabilizing interactions rather than the actual catalytic process of splicing ^39^. The peripheric nature of SF3B2 raises intriguing possibilities related to the precise mechanism of snaR-A-driven splicing perturbation. snaR-A-triggered U2 instability, for example, would conceivably be maximally disruptive for mRNAs with weak splicing characteristics^40^. The confluence of additional regulatory layers, such as recruitment of specific RBPs and ancillary splicing factors, may ultimately produce the added specificity of snaR-A-related splicing defects.

In addition to U2 residency, nuclear-speckle proximity was a surprisingly important predictor of splicing defects that were improved following snaR-A depletion. This result led us to a counterintuitive finding of higher intron retention levels observed across transcripts produced by speckle-proximal genes. Although ectopic targeting of genes to nuclear speckles is shown to increase splicing efficiency (consistent with the expectation of enhanced splicing activity at sites enriched for splicing factors), recent work has linked genes with high G/C content and short introns to inefficient splicing and nuclear speckle retention^31,36^. Though its subnuclear localization provides preliminary evidence that snaR-A may perturb splicing either at or near speckles, the convergence of U2-binding, intron length, sequence content, and speckle proximity altogether obscure whether one feature is particularly significant. Even so, whether the subnuclear coalescence of splicing factors itself is simply a consequence of snRNP retention at unspliced events, thereby driving nucleation of otherwise efficient splicing bodies, remains a possibility.

Deeper analysis of the mRNA subpopulation affected by snaR-A levels uncovered an overrepresentation of histone deacetylases (HDACs) and factors involved in autophagy. The implications of snaR-A interference of factors involved in chromatin remodeling and autophagy are likely complex and context-dependent, given roles for these processes in both tumor initiation and suppression^41,42^. Nevertheless, we show that resolution of processing defects following snaR-A depletion leads to increased protein abundance for these and several other factors, including OGFR - a putative tumor suppressor - as well as G3BP2 - a stress granule protein^43^. With respect to G3BP2 disruption, we find that stress granules are notably larger in snaR-A depleted cells, suggesting snaR-A expression may interfere with cellular stress responses (Supplemental Figure 8). Thus, the emergence of snaR-A and its downstream perturbation of splicing is very likely to have wide-ranging implications. We speculate that Pol III production of snaR-A may essentially introduce “sand into the gears” of splicing and that various protein and macromolecular vulnerabilities to these defects may ultimately allow for opportunistic selection of alterations that promote growth- and survival-related pathways.

## Limitations of this study

Our study made use of available ATAC-seq (chromatin accessibility) datasets to predict snaR-A gene activity in human tumors, given multiple practical benefits of ATAC (e.g. uniformly applied experimental and sequencing approaches, direct indication of gene-level activity rather than steady-state RNA abundance, etc.). The interpretation of this approach is nevertheless limited by the indirect nature of assessing Pol III transcription through gene accessibility, whereas a direct measure of nascent snaR-A might naturally have been preferred. Even so, RNA-sequencing methods are highly variable in nature and the current absence of nascent-seq experiments applied across 100s of human tumors precludes any possibility of broadly assessing snaR-A expression and clinical outcome signatures. This holds similarly true for methods that directly map RNA polymerase III occupancy.

The *in vitro* nature of our biotin-snaR RNA pull-down experiments may not accurately reflect the subcellular localization, protein stoichiometry, or cellular modification state of snaR-A ncRNA, and thus alternative methods of RNA-protein interaction mapping may have captured protein interactions that our approach did not identify. We note that our identification of SF3B2 as a primary interaction candidate was further supported by CLIP and PAR-CLIP experiments. Nevertheless, snaR-A may interact with many other proteins, including additional splicing factors unaccounted for in our study.

## Materials and Methods

### Cell lines and culture conditions

THP-1 monocytes (ATCC) were cultured in RPMI 1640 Medium (Corning), and HEK293T (ATCC) were cultured in complete Dulbecco’s Modified Eagle’s Medium (DMEM) (Corning), both supplemented with 10% fetal bovine serum (Corning) and 1% penicillin-streptomycin (Gibco). THP-1 cells were maintained at 2-8 × 10^5^ cells/mL. HEK293T cells were maintained in 10-cm dishes by seeding 1 × 10^6^ cells with complete medium every 2–3 days. All cells were maintained in a humidified atmosphere at 37 °C with 5% CO_2_.

### Plasmids and cell transfection

pcDNA3.1+/C-(K)DYK plasmids expressing eGFP, SF3B2 (Catalog#OHu26570D), La/SSB (Catalog#OHu28669D), NF90 (Catalog# OHu26506D), NF110 (Catalog# OHu26447D), U6-promoter-snaR-A plasmids, and U6-promoter-scramble-snaR-A plasmids were obtained from GenScript. HEK293T cells (1 × 10^5^ cells/cm^2^) were seeded into 12-well Cell-Culture Treated Multidishes (Thermo Scientific) and incubated overnight. Plasmids were transfected using Lipofectamine 3000 (Invitrogen) according to the manufacturer’s protocol. Cells were collected 48 hours post transfection.

### Small interfering RNA (siRNA) transfection

siRNAs were mixed with OPTI-MEM media and transfected with Lipofectamine RNAiMAX Transfection Reagent (Invitrogen) per manufacturer’s instructions. Three siRNAs against snaR-A and scramble siRNA control were added at a final concentration of 12.5nM. RNA was extracted 48 hours post-transfection.

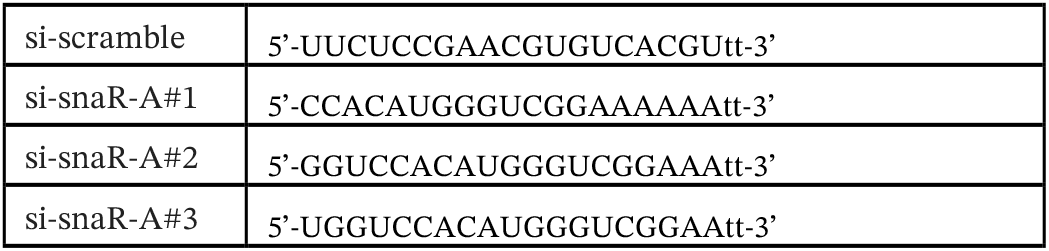

### UV cross-linking and immunoprecipitation (CLIP)

HEK293T cells (∼ 5 ×10^6^ cells per 6 mm dish, 1 dish per sample) were transfected with FLAG-GFP and FLAG-SF3B2 (GenScript) (14 µg DNA per dish) using Lipofectamine 3000 Transfection Kit (Invitrogen) per manufacturer’s instruction. Cells were washed once with 1× PBS (Corning) 48 hours post-transfection. Cells were then UV cross-linked at 254-nm with an energy of 400 mJoules/cm^2^. Cells were lysed with 500ul Pierce IP lysis buffer (Thermo Scientific), supplemented with Halt™ Protease Inhibitor Cocktail (Thermo Scientific) and RNase inhibitor (New England Biolabs) for 5 min on ice. The lysates were cleared by centrifugation at 140,000 g for 10 min at 4 °C. To co-immunoprecipitate RNA-bound FLAG-GFP and FLAG-SF3B2, cleared lysates were mixed with anti-FLAG-antibody-conjugated magnetic beads (Thermo Scientific), and were incubated at room temperature with rotation for 1 hr. Beads were washed two times with high salt wash buffer (50mM Tris-HCl pH 7.4, 1M NaCl, 1mM EDTA, 1% NP-40, 0.1% SDS), one time with IP lysis buffer (50 mM NaCl, 25 mM Tris (pH=7.5), 1mM EDTA, 1%NP-40, 5% glycerol) and one time with molecular grade water followed by incubation with proteinase K (New England Biolabs) at 55 °C for 30 minutes with constant shaking. The bound RNA was extracted using mirVana PARIS RNA and Native Protein Purification kit (Invitrogen) following the manufacturer’s instructions for total RNA extraction.

### RNA immunoprecipitation (RIP)

HEK 293T cells (∼ 5 ×10^6^ cells per 6 mm dish, 1 dish per sample) were transfected with FLAG-GFP, FLAG-NF90, FLAG-NF110, or FLAG-SSB (14 µg DNA per dish) using Lipofectamine 3000 Transfection Kit (Invitrogen) per manufacturer’s instruction. Cells were harvested 48 hours post-transfection and lysed in Pierce IP lysis buffer (Thermo Scientific), supplemented with Halt™ Protease Inhibitor Cocktail (Thermo Scientific) and RNase inhibitor (New England Biolabs) for 5 min on ice. The lysates were cleared by centrifugation at 140,000 g for 10 min at 4 °C. To co-immunoprecipitate RNA-bound FLAG-GFP, FLAG-NF90, FLAG-NF110, and FLAG-SSB, cleared lysates were mixed with anti-FLAG-conjugated magnetic beads (Thermo Scientific), and the binding reaction was incubated for 1 h at room temperature with rotation. The beads were washed three times with IP lysis buffer (50 mM NaCl, 25 mM Tris (pH=7.5), 1mM EDTA, 1%NP-40, 5% glycerol) and one time with molecular grade water followed by incubation with proteinase K (New England Biolabs) at 55 °C for 30 minutes. The bound RNA was extracted using mirVana PARIS RNA and Native Protein Purification kit (Invitrogen) following the manufacturer’s instructions for total RNA extraction.

### RNA isolation and RT-qPCR

RNA were isolated using E.Z.N.A.® Total RNA Kit I (Omega Bio-tek), with 100% ethanol instead of 70% ethanol for inclusion of small RNA. For small RNA transcripts, reverse transcription of RNA (10-200ng) was performed using the TaqMan MicroRNA Reverse Transcription Kit (Applied Biosystems) with the target-specific Taqman™ RT primers (Thermo Fisher) according to manufacturer’s instructions. For mRNA, reverse transcription of RNA (1µg) was performed using High-Capacity cDNA Reverse Transcription Kit (Applied Biosystems). Quantitative PCR (qPCR) was performed using TaqMan™ Fast Advanced Master Mix (Thermo Fisher) and predesigned TaqMan™ Gene Expression Assays (Thermo Fisher) for selected genes. small RNA abundances were normalized against the geometric mean of Z30 and U19. mRNA abundance are normalized against the geometric mean of GAPDH and ACTB. The fold change of each target gene relative to controls was calculated using the Comparative CT Method (ΔΔCT Method).

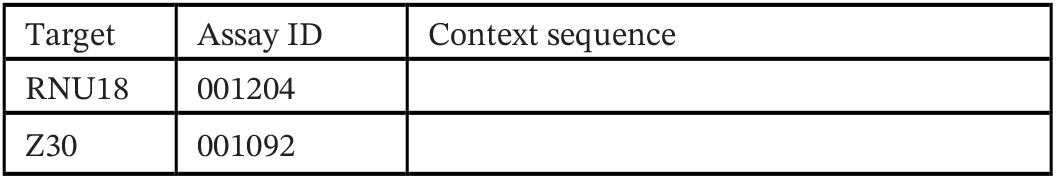
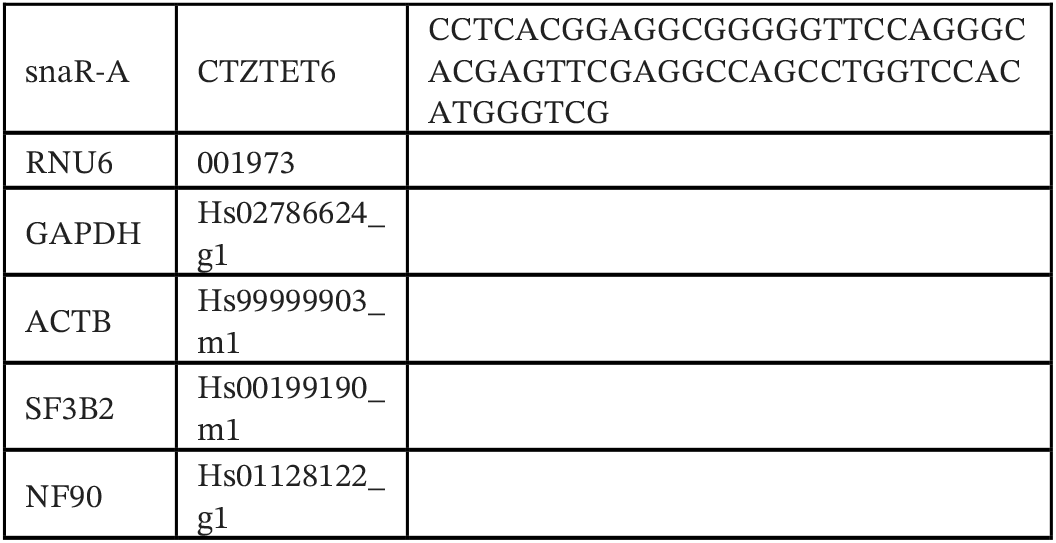

### Immunoblotting

Immunoblotting was performed as previously described^44^. Briefly, transfected cells were lysed with cell disruption buffer (Thermo Scientific) for 5 minutes on ice, followed by centrifugation at 140,000 g for 5 minutes at 4 °C. The lysate were separated on a 4–20% Mini-PROTEAN® TGX Stain-Free™ Protein Gels (BIO-RAD), transferred to polyvinylidene difluoride membranes (0.2 um) (Invitrogen). Transfer membranes were blocked with 5% blotting-grade blocker (BIO-RAD) in 1×PBST (Cytiva), followed by incubation with the primary antibodies at 4°C overnight. Membranes were washed 3× 5min with 1×PBST and incubated with mouse or rabbit secondary antibodies conjugated with horseradish peroxidase (Invitrogen) for 1 h at room temperature followed 3 × 5 min wash by 1× PBST. Proteins were visualized using SuperSignal WestFemto (ThermoScientific) with ChemiDoc™ Touch Imaging System (BIO-RAD). Protein abundances were calculated by normalizing to total protein. Fold changes were calculated agaisnt the mean of the controls.

| Target | Cat number | dilution |
| --- | --- | --- |
| La (SSB) | sc-80656 (Santa Cruz Biotechnology) | 1:1000 |
| SF3A1 | 15858-1-AP (proteintech) | 1:1000 |
| SF3A3 | 12070-1-AP (proteintech) | 1:1000 |
| SF3B4 | 10482-1-AP (proteintech) | 1:1000 |
| SF3B2 | A5875 (ABclonal) | 1:1000 |
| MCRIP2 | 20808-1-AP (proteintech) | 1:1000 |
| OGFR | 11177-1-AP (proteintech) | 1:1000 |
| TIMELESS | 14421-1-AP (proteintech) | 1:1000 |
| HIP1R | 16814-1-AP (proteintech) | 1:1000 |
| ILF3 | 19887-1-AP (proteintech) | 1:1000 |
| MTA1 | 30545-1-AP (proteintech) | 1:1000 |
| AUP1 | 13726-1-AP (proteintech) | 1:500 |

### RNA pulldown and mass spectrometry

THP-1 cells (8 million cell/sample) were lysed in 300 ul CE buffer (10 mM HEPES, 150 mM KCl, 1mM EDTA, 0.075%(v/v) NP40, 1mM DTT, 1mM PMSF), followed by centrifugation at 140,000 g for 10 min at 4 °C to clear the lysate. 400 pmol Biotinylated snaR-A or biotinylated scramble sequence of snaR-A (IDT) were pre-heated to 65°C for 10 minutes and cooled down to 4°C to restore proper secondary structures. Pierce Streptavidin Magnetic Beads (80ul/sample) (Thermo Scientific) were washed twice with an equal volume of 20mM Tris (pH 7.5) and resuspended in an equal volume of RNA Capture Buffer (20mM Tris (pH 7.5), 1M NaCl, 1mM EDTA). Biotinylated snaR-A or the scramble sequence was mixed with pre-washed Pierce Streptavidin Magnetic Beads and incubated at room temperature with rotation for 30 minutes. The beads were then washed twice with an equal volume of 20 mM Tris (pH 7.5) and resuspended in THP-1 cell lysate. The beads were incubated at 4°C for 2 hours. After incubation, beads were washed three times with TEKN buffer (10 mM Tris (pH 8), 1 mM EDTA, 250 mM KCl, 0.1% (v/v) NP40) followed by one last wash with molecular grade water (Lonza). The beads were then subjected to RNase digestion in a 25 ul reaction containing 10mM Tris-HCl (pH 7.5), 1 mM MgCl2, 40 mM NaCl, and 10ul of A/T1 RNase (Thermo Scientific), for 1 hr at 37°C^45^. The RNA-bound proteins were then released in the reaction and subjected to mass-spectrometry by nano-LC-MS/MS (Ultimate 3000 coupled to a Q-Exactive HF, Thermo Scientific) at the University of Illinois Carver proteomics core.

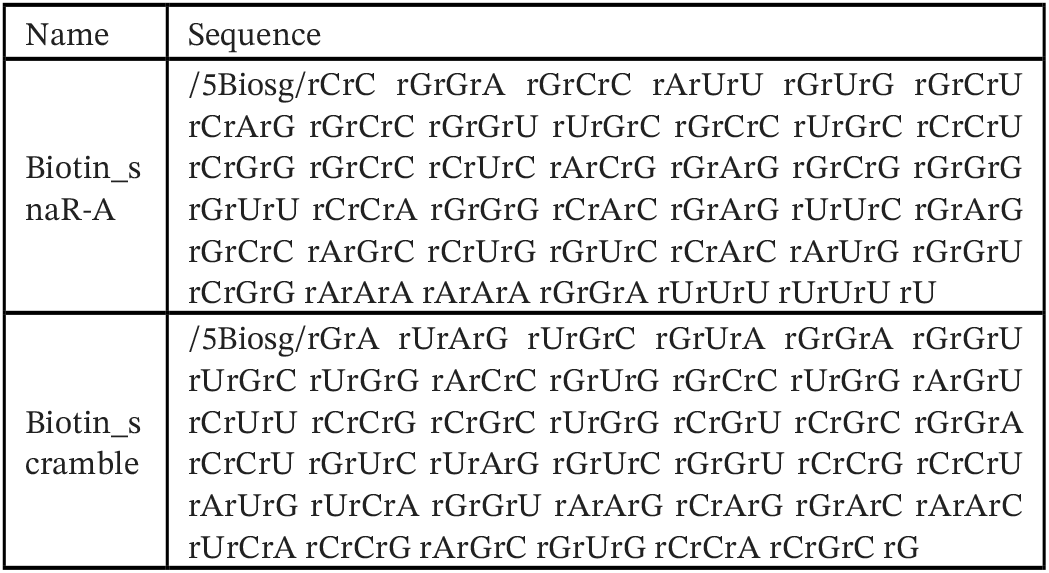

### Proteomics analysis

Peptide intensities were normalized to the total intensity of each individual sample. Proteins enriched for snaR-A binding were identified using the following criteria: the presence of at least one unique peptide in both snaR-A pulldown samples and a log_2_ ratio of normalized intensity (snaR-A pulldown) to normalized intensity in both scramble pulldown and beads pulldown greater than 2.

### RNA FISH-IF

To visualize snaR-A localization relative to nuclear speckles in HEK293T, we performed HCR™ RNA-FISH together with immunofluorescence (IF) to detect SON as a marker for nuclear speckles. In brief, 24 h after plating of the cells, the samples were rinsed once with 1× PBS (Corning) and then fixed in 4% paraformaldehyde. After fixation, we rinsed the samples twice with 1× PBS then permeabilized in 70% ethanol overnight at -20°C. For hybridization, we rinsed the samples twice with 1× PBS and then 1× SSC. Samples were then pre-hybridized in pre-heated hybridization buffer (Molecular Instruments) for 30 min at 37°C. The pre-hybridization buffer was then removed and hybridization buffer containing RNA FISH probe targeting snaR-A (4nM) was then added to the samples and the samples were incubated overnight at 37°C in a humidified incubator. The probes were designed by Molecular Instruments, Inc. Excess probes were removed by rinsing the samples 3× 5min with 75%, 50%, and 25% wash buffer (Molecular Instruments) diluted with 5× SSCT (5× sodium chloride sodium citrate, 0.1% Tween 20), followed by 3× 5min wash with 5×, 2×, and 1× SSCT at room temperature. Samples were then pre-amplified in amplification buffer (Molecular Instruments) for 30min at room temperature. Hairpin h1 and h2 (molecular instruments) were snap-cooled after incubation at 95°C for 90 seconds to room temperature for 30 min. Hairpin solutions were prepared by mixing hairpin h1 and h2 in amplification buffer and added to the samples. Samples were incubated in the dark at room temperature overnight. Excess hairpins were removed by 5× 5min wash with 5×, 2×, 1× SSCT, and PBST at room temperature. Samples were then blocked with blocking buffer (5% BSA, 5% goat serum in 1× PBST) for 1hr at room temperature. Samples were washed once with 1× PBST for 5min followed by incubation of primary antibody against SON provided by Dr. Andrew Belmont (University of Illinois) at 1:2000 dilution. A secondary antibody was applied after 3× 5min PBST wash for 1hr at room temperature. Samples were again washed with 3× 5min PBST. We added 50 µg ml–1 DAPI and Wheat Germ Agglutinin (WGA) to stain for nuclei and cell membrane respectively. We rinsed the samples twice with 1× PBS, added Vectashield Antifade solution and proceeded with imaging on a ZEISS LSM 900 confocal. For each sample, we acquired z stacks at 0.5 µm intervals using a 60× oil objective.

### Intron-Exon Junction PCR

Total RNA after scramble and snaR-A knockdown was extracted using E.Z.N.A.® Total RNA Kit I (Omega Bio-tek) according to the manufacturer’s instructions. Genomic DNA was removed using DNase (Thermo Scientific). RNA (1 µg) was reverse transcribed using High-Capacity cDNA Reverse Transcription Kit (Thermo Fisher), and PCR was performed using GoTaq® Green Master Mix (Promega). The splicing efficiency is monitored using RT-PCR with primers designed to target the upstream and downstream exons. Amplified products were separated on a 1% agarose/TAE gel with SYBR™ Safe DNA Gel staining (Invitrogen) and visualized on a Bio-Rad ChemiDoc Imager. The percent spliced-in (PSI) is quantified intensity_retained_/(intensity_retained_+intensity_spliced_) ImageLab.

### RNA sequencing and analysis

RNA purified from si-scramble, si-snaR-A#1,#2, #3 treated HEK 293T cells using E.Z.N.A.® Total RNA Kit I (Omega Bio-tek) were subjected to library preparation after rRNA depletion and poly(A)^+^ RNA enrichment using the Kapa Hyper Stranded mRNA library kit (Roche) according to manufacturer’s instructions. All high-throughput sequencing libraries were sequenced for 121 cycles using NovaSeq 6000 at the Univerisity of Illinois, Roy J.Carver Biotechnology Center. poly(A)^+^ RNA reads were first trimmed using TrimGalore^46^ and then mapped to the transcriptome using quasi-mapper Salmon to generate read counts^47^. Differentially expressed RNA was determined using edgeR^48^.

### eCLIP/iCLIP-seq analysis

eCLIP-seq FASTQ files were retrieved from the ENCODE web portal (https://www.encodeproject.org/)^23^. Reads were first trimmed using TrimGalore^46^ and then mapped to the genome (GRCh38) using Bowtie2^49^. PCR duplicates from ENCODE eCLIP-seq were removed using UMI-Tools^50^. Read counts were extracted over a comprehensive RNAcentral database annotation of noncoding RNA using BEDTools^15,51^. The extracted read counts were then quantile normalized. The fold-enrichment of each RNAcentral entry was calculated against the maximum between corresponding control eCLIP-seq or the average of all control eCLIP-seq datasets.

### Gene “ON”/ “OFF” analysis

Primary solid tumor ATAC-seq alignment bam files were retrieved from the Genomic Data Commons Data Portal (https://portal.gdc.cancer.gov). All ATAC-seq samples (328) were equivalently down-scaled to 250 million total reads to ensure the baseline strength for determining statistical significance. Signals were extracted over a comprehensive RNAcentral database annotation of noncoding RNAs and the expected signals were computed as the local background regions including 1kb (λ1k), 10kb (λ10k), 100kb (λ100k), and the whole genome (λgenome) for each RNAcentral gene coordinate entry. The λ value is thereafter determined as the maximum λ value of the distance set. The probability density function (PDF) utilized is given by P(X = x) = (e^(-λ) * λ^x) / x!. If k represents the observed signal for the specific gene, the p-value tied to a specific gene enrichment is calculated using P(X > k). P-values were adjusted globally using the Benjamini-Hochberg method. Genes shorter than 1000 bp with a median p-value less than 0.05 across different cancer contexts were considered always “ON.” For the remaining genes, those with a median p-value less than 0.05 in at least one cancer subtype were considered “ON/OFF”; otherwise, they were considered “OFF”.

### Survival analysis

Gene-specific hazard ratios (HR) and Wald statistics were determined by applying Cox proportional hazard models to our map of gene “ON/OFF” status, using the ‘survival’ R package^20^. For each gene, analyses were restricted to cancer subtypes in which both ON and OFF events were present, thereby preventing [1] analysis of genes with constitutively ON or OFF states, and [2] avoiding the influence of tissue-specific characteristics. For example, snaR-A is “ON” in 100% of clinically matched TGCT tumors (related to expression in testes); and thus TGCT is not included in the survival analysis for snaR-A. Down-stream analyses were restricted to genes < 400 bp. For comparison across Pol III-transcribed genes, a median Wald statistic was assigned to cohorts of genes encoding a particular RNA type (e.g. 5S rRNA represents all genes encoding 5S rRNA, as snaR-A represents all snaR-A genes). For each RNA type, a z-score was calculated by comparing the median Wald statistic for a given gene set against an empirically determined null distribution that accounts for the same number of genes.

### RNA secondary structure prediction

snaR-A secondary structure was predicted using the Vienna RNAfold web server with standard settings^52^.

### Differential splicing (Intron-centric) analysis

Alternatively, spliced events were identified using rMATs-turbo^55^ with a default splicing difference cutoff of 0.0001. These events were determined based on reads mapping to splice junctions and are categorized into five types: retained intron (RI), skipped exons (SE), alternative 5’ splice site (A5SS), alternative 3’ splice site (A3SS), and mutually exclusive exons (MXE). To determine the maximum number of overlapping events among three different siRNAs targeting snaR-A, events were ranked by a significance-weighted changes in percent spliced-in (ΔPSI) (-log_10_(FDR)*ΔPSI). We performed 10,000 permutations to calculate the expected number of overlapping events for the three different siRNAs. Permutations were performed separately for inclusion and exclusion events for each type of splicing event. A p-value cutoff of 0.05 was used to determine the number of events to be considered as overlapping. The permutations were done separately for inclusion and exclusion events for each type of splicing event. After identifying overlapping events for the three siRNAs targeting snaR-A, events were further filtered using a minimum false discovery rate (FDR) of < 0.05 and a maximum absolute PSI value of >0.1 across all samples (in total, 287 introns).

### Transcript-centric analysis

Global intron retention (IR) ratios were determined using IRFinder by mapping to the human genome GRCh38 and gencode v44 primary assembly annotation^53^. IR events were thereafter filtered on read quality metrics, including a minimal gene expression score (CPM > 1), plus minimal read coverage spanning the flanking 3’ and 5’ exons (> 10 across all samples, taken individually). Post-filtered IR ratios were then compared between control (scramble) and the distribution of IR ratios for the three independent siRNAs designed against snaR-A. Individual intron IR scores were thereafter defined as the -log_10_(pvalue), determined by Q-test on the distribution of IR ratios across control and si-snaR-A experiments. In scenarios of overlapping intron annotation (i.e. a larger parent or smaller “nested” intron), a median IR score was assigned to a singular intron entity. The significance of transcript-wide IR scores was then assessed by comparing the full distribution of intron IR scores to an empirically derived null distribution for a given number of events (introns). This approach was applied using two frameworks, the first [1] accounting only for introns with positive IR scores (-log_10_(pvalue)>0), and the second [2] accounting for all introns, including those with 0 scores (Supplemental Figure 5). Finally, a list of features with significant transcript-wide reduction in IR was determined as any gene with an adjusted permutation p-value < 0.05 in both frameworks [1] and [2] (in total, 136 genes).

### SON TSA-seq (Nuclear Speckle distance) analysis

All available SON TSA-seq experiments with normalized signal counts were retrieved from the 4D Nucleome portal (https://www.4dnucleome.org/)^54,55^. In total, 25 experimental datasets derived from H1 hESC, K562, HFFc6, and HCT116 cells were compiled into an average SON TSA-seq score, thereby representing the average positional relationship between a given gene and nuclear speckles in currently tested cell lines.

### Logistic regression model of IR

Transcripts that showed significant reductions in IR levels (see above) were labeled as ‘snaR-A-disruption-sensitive’ (136 transcripts), while the remaining transcripts were labeled as ‘snaR-A-disruption-insensitive’ (10,488 transcripts). A penalized logistic regression model, incorporating both L1 and L2 regularization techniques^56^, was trained using 277 features, which included transcript length, median intron length, gene G/C content, distance to nuclear speckles (SON TSA-seq score), and RNA-binding protein (RBP) occupancy data from the ENCODE database (Supplemental Table 2). To address the class imbalance between ‘sensitive’ and ‘insensitive’ transcripts, an oversampling technique was employed. Model training was performed using leave-one-out cross-validation, with bootstrapping to conduct 100 simulations for model generation. The average area under the receiver operating characteristic (ROC) curve (AUC) was calculated to assess the model’s predictive performance on unseen data. A reduced model was constructed using selected variables after previous variable selection step, including gene length, gene G/C content, median intron length, distance to nuclear speckles, SF3B4 occupancy, and SF3A3 occupancy. A mean predicted probability for each transcript for snaR-A sensitivity was calculated from the reduced model by running 100 simulations and taking the mean between across 100 simulations.

Introns that showed significant reductions in IR levels (see above) were labeled as ‘snaR-A-disruption-sensitive’ (218 introns), while the remaining introns were labeled as ‘snaR-A-disruption-insensitive’ (8,558 introns). A separate penalized logistic regression model, also incorporating L1 and L2 regularization, was trained using 277 features, which included intron G/C content, G-quadruplex density predicted using G4Hunter^57^, intron length, and RBP occupancy. Model training and evaluation were conducted similarly, using leave-one-out cross-validation with 100 bootstrapped simulations. The average AUC was calculated to measure the model’s predictive performance.

### U2 snRNP-residency score calculation

500 bp upstream and downstream of every 3’ intron-exon junctions of expressed genes are binned into 100 10bp binds. Reads mapping to each bin for all of the junctions are extracted from SF3B4 (HepG2) and SF3A3 (HepG2) eCLIP-seq (ENCODE) using bedtools. Enrichment scores for 5 bins into the introns of the 3’ junction were determined using a Poisson framework. Briefly, for each bin, the expected signal within 500bp upstream (λup), 500bp downstream (λdown), and anaverage across the entire 1000bp window for all transcript (λwhole) is computed. The λ value is thereafter determined as the maximum lambda value of the distance set. The probability density function (PDF) utilized is given by P(X = x) = (e^(-λ) * λ^x) / x!. If k represents the observed signal for the bin, the p-value tied to a specific bin enrichment is calculated using P(X > k). Finally, a U2 snRNP residency score is computed as the negative logarithm (base 10) of the minimum p-value among the 5 bins.

### Overrepresentation analysis of RBP patterns (eCLIP)

Functional enrichment analysis was performed for genes encoding 154 RNA-binding proteins (RBPs) from the ENCODE eCLIP-seq dataset. Information on the transcripts bound by these 272 RBPs was retrieved from the ENCODE database^23^. Proteins associated with each Gene Ontology (GO) term (cellular component, CC) were considered as functional groups for collective analysis.The significance of changes in IR levels of bound transcripts for each functional protein group was determined by comparing the observed distribution of z-scores from the transcript-centric analysis against an empirical null distribution derived from 1,000 permutations. The calculated p-values were then transformed into z-scores.

### Functional enrichment analysis

Gene Ontology enrichment analysis was performed using the R package clusterProfiler^58^ with genome-wide annotation for Human org.Hs.eg.db ^59^. The gene set was defined as transcripts with significant transcript-wide (z < -3) and/or intron-centric (padj. < 0.01) scores and predicted snaR-A sensitivity (n=298) (i.e. both observed and predicted snaR-A sensitivity), and compared against a universe of genes with prediction scores (n=10,637). Enrichments were calculated for GO cellular component (CC), Molecular Function (MF), and Biological Process (BP), and filtered for ontologies with duplicate gene groups and/or <= 3 factors. GO enrichment p-values were adjusted using the Benjamini–Hochberg method.

## Resource Availability

The RNA-seq data generated for this study are available through the NCBI Gene Expression Omnibus with accession number **GSE271057**.

## Acknowledgements

We thank Alvaro Hernandez, Chris Wright, Danman Zhang, and staff at the Carver Biotechnology Center for sequencing services, and administrators of the Carl R. Woese Institute for Genomic Biology (UIUC) Biocluster for computational support. We thank Peter Yau, Justine Arrington, Brian Imai, and staff at the Carver Biotechnology Center for proteomics services. We thank Prof. Andy Belmont, Prof Kannanganattu V. Prasanth, Prof Stephanie Ceman, and members of the Van Bortle lab and the Prasanth lab for helpful suggestions. We thank the Belmont lab for generously sharing anti-SON primary antibodies. This work was supported by the National Institutes of Health, National Human Genome Research Institute (NHGRI) grant R00HG010362 to K.V.B., and National Heart, Lung, and Blood Institute (NHLBI) grant R01HL126845 and Chan-Zuckerberg Biohub Chicago’s Investigator Award to A.K.

## Author contributions

Study design: SZ, KVB. Data collection: SZ, SL, LM, SC, RC, KVB. Data analysis: SZ, SL, LM, KVB, RKC. Data interpretation: SZ, AK, KVB. Writing: SZ and KVB with comments from all authors

## Supplemental Figures

**Supplemental Figure 1.**
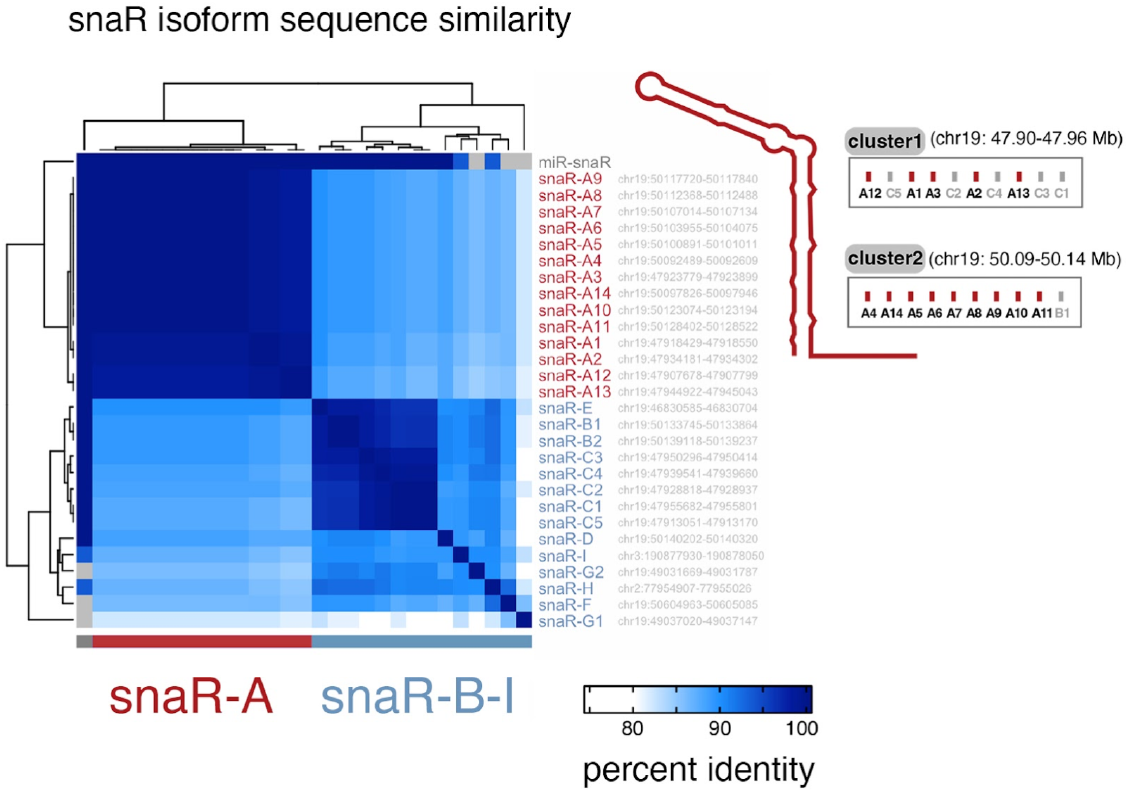
Sequence similarity of snaR-A, -B, -C, -D, -E, -F, -G, -H, and -I isoforms. Hierarchical clustering of individual snaR isoform sequences on the basis of sequence similarity (percent identity), with sequences corresponding to snaR isoform A (snaR-A; red) notably distinct from those corresponding to snaR isoforms B-I (blue). Chromosome and genomic intervals correspond to GRCh38. Inset highlights the individual positions of snaR-A genes in clusters 1 and 2 on chr19, mapped to isoform number.

**Supplemental Figure 2.**
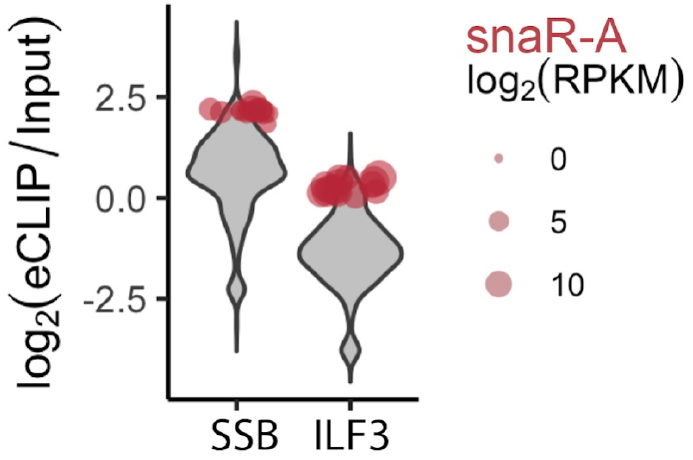
snaR-A enrichment across La (SSB) and ILF3 eCLIP experiments, compared against Pol III-transcribed genes. Violin plots represent the eCLIP enrichment distributions of all Pol III-transcribed genes for La (left) and ILF3 (right). snaR-A genes, highlighted in red, are among the top enriched RNA species in both experiments

**Supplemental Figure 3.**
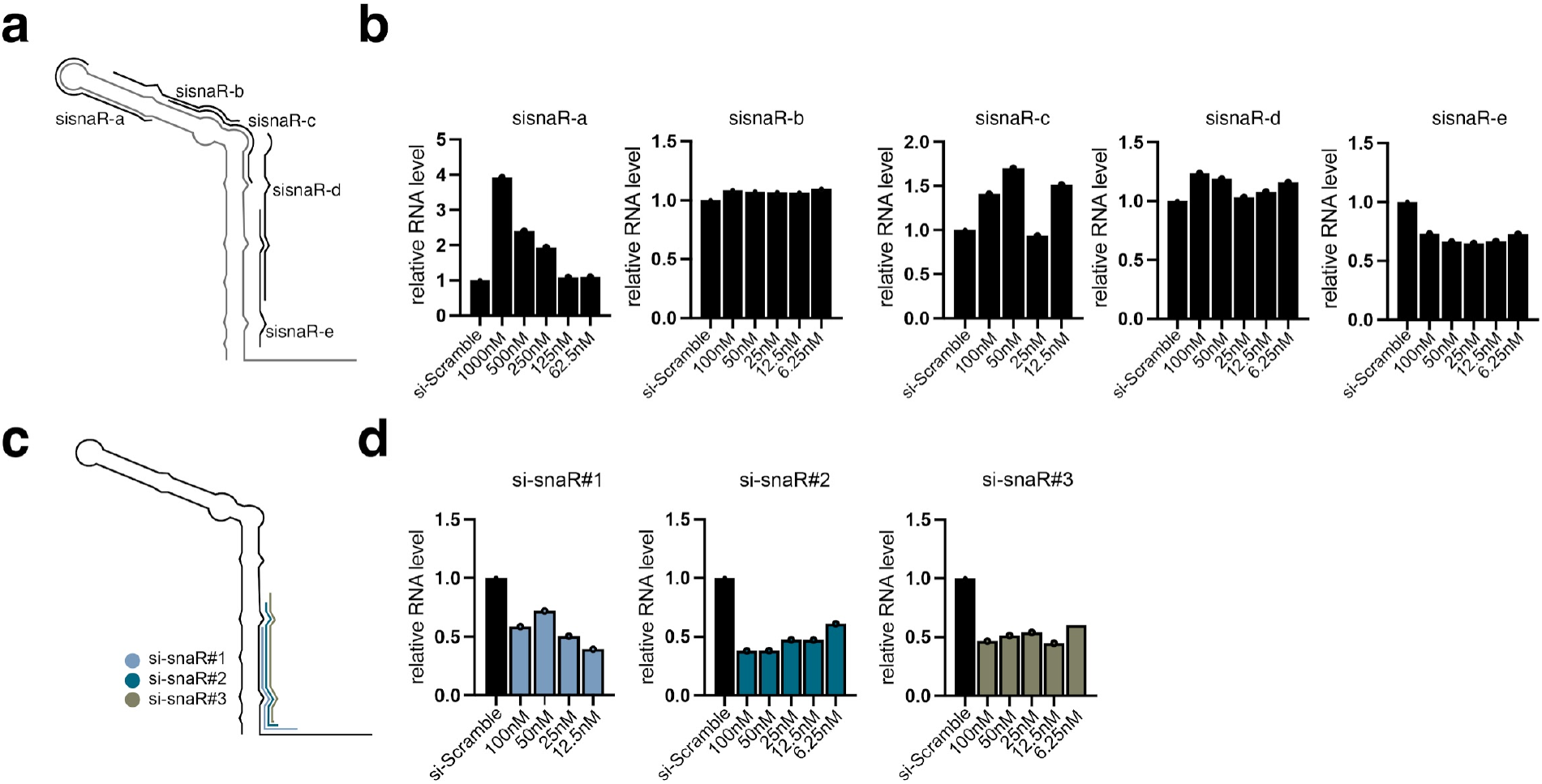
Identification of effective siRNA target sequences within snaR-A. **(a)** illustration of several siRNA target regions tested against snaR-A that were unsuccessful in reducing snaR-A levels. **(b)** RT-qPCR analysis of snaR-A levels at various concentrations of siRNAs illustrated in panel A. **(c)** illustration of siRNA target regions for siRNA #1, #2, and #3, which were successful in reducing snaR-A levels by >= 50%. **(d)** RT-qPCR analysis of snaR-A levels at various concentrations of siRNAs illustrated in panel C.

**Supplemental Figure 4.**
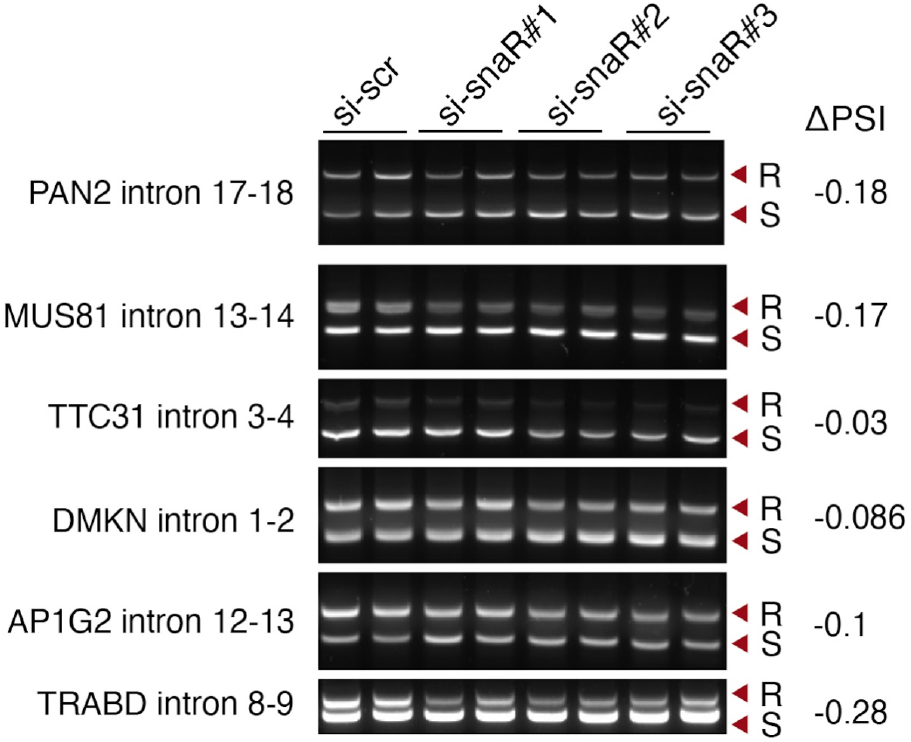
PCR analysis and validation of differential intron retention splicing events identified by rMATS following snaR-A depletion. PCR analysis of intron retention (IR) levels corresponding to specific introns determined to be significantly downregulated following snaR-A depletion. Introns were tested for several independent transcripts (PAN2, MUS81, TTC31, DMKN, AP1G2, and TRABD). Arrows indicate retained (R) and spliced (S) introns. Delta PSI (Percentage Spliced In) values indicate the change in IR levels following snaR-A depletion, negative values indicate a lower proportion of transcripts with the intron retained (i.e. decreased intron retention).

**Supplemental Figure 5.**
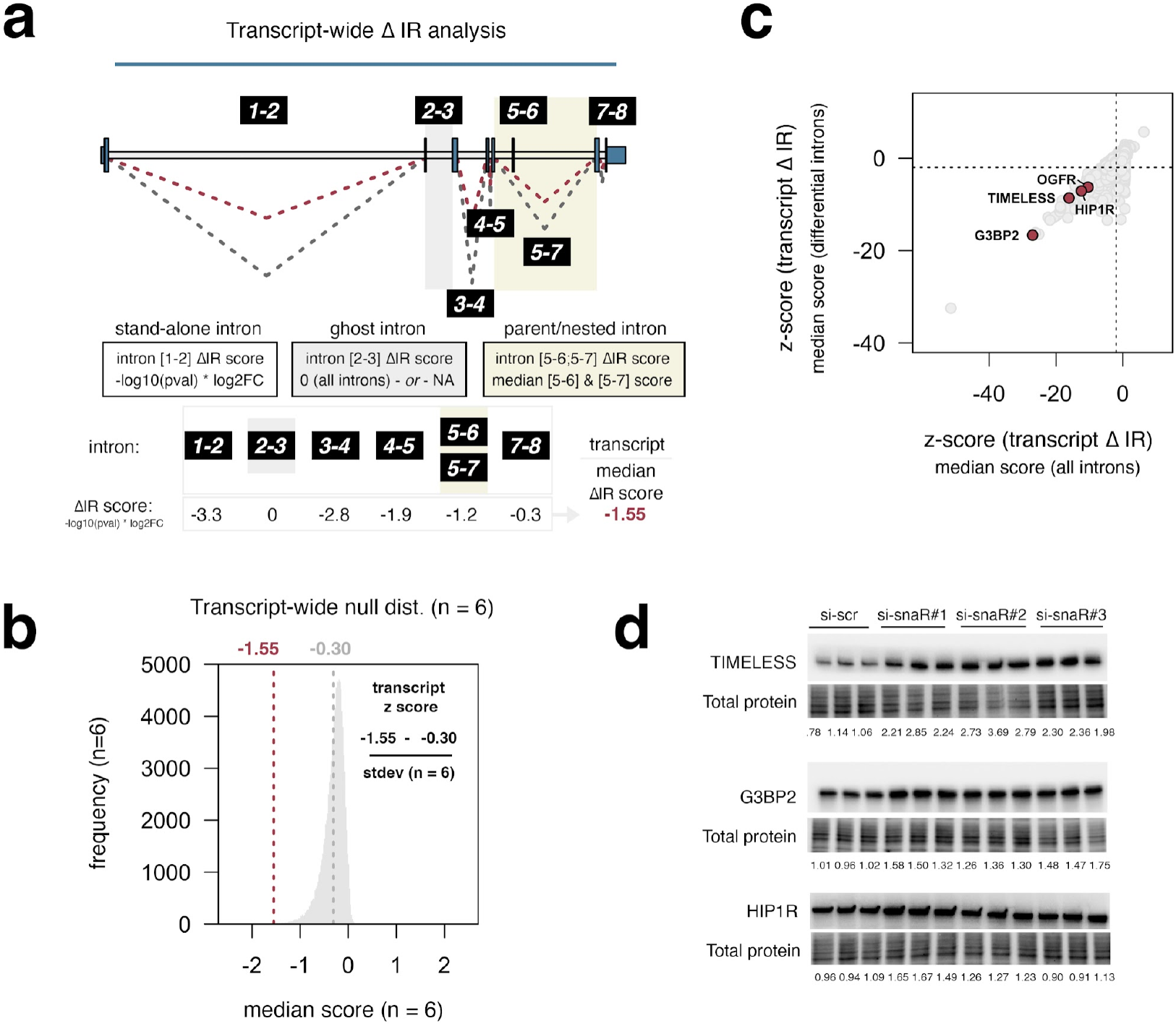
Transcript-wide analysis of differential intron retention levels and survey of protein disruption effects following snaR-A depletion. **(a)** Overview of transcript-wide quantification of differential splicing events. Introns are named by upstream and downstream exon numbers; intron retention (IR) levels are compared between control (scramble, gray) and three si-snaR-A (3 unique siRNAs, 3 biological replicates each. red). The median differential intron score for a given transcript, initially measured as the -log10(p-val) * log2FC for each intron, is compared against a randomization-based null distribution of the same number of intron events. In cases of overlapping (parent and nested) intron annotations, a median intron score is assigned to the cluster of overlapping features. Ghost (undetected) introns are either considered as 0 values (“all introns”) or omitted from further analysis. **(b)** Standardized z-scores are assigned to each transcript on the basis of its observation, distribution mean, and standard deviation of its corresponding null distribution (example illustration for transcript in panel a with 6 intron events). **(c)** Analysis of the transcript-wide survey results including ghost (“all”) intron scores (x-axis) versus the same survey with ghost introns omitted (y-axis) identified 136 unique genes with significant differential IR levels following snaR-A depletion (adjusted p-val < 0.05 in both approaches). **(d)** Immunoblot analysis and quantification of protein levels corresponding to genes with differential IR levels (TIMELESS, G3BP2, and HIP1R; highlighted in panel c)

**Supplemental Figure 6.**
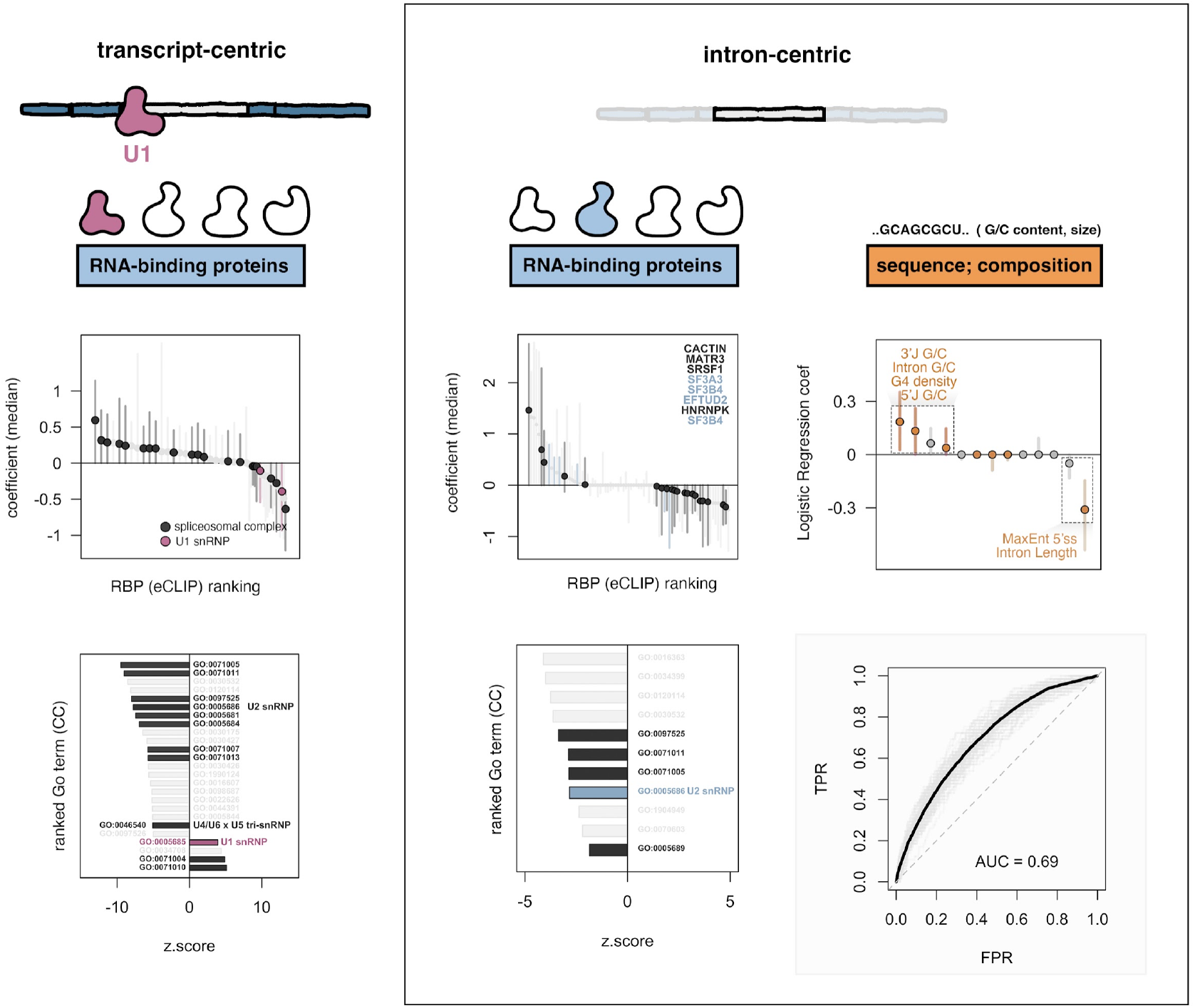
Transcript-centric and intron-centric logistic regression coefficients for specific RNA-binding proteins and intrinsic sequence features. Transcript-centric analysis of regression coefficients across all RBPs (top-left), and mRNA population snaR-A sensitivity as a function of RBP occupancy grouped by GO:CC term (bottom-left). Transcript-centric plots (left) are related to Figure 3e, re-colored to emphasize U1 snRNP. Intron-centric analysis of regression coefficients similarly identifies enrichment for U2 snRNP proteins (top-middle), and increased snaR-A sensitivity for mRNAs bound by U2 snRNP, as defined by GO:CC term (bottom-middle). Intron-centric analysis of regression coefficients, restricted to intrinsic features (i.e. sequence composition, top-right), identifies G/C content, particularly at the 3’ intron/exon junction, as a positive predictor of snaR-A sensitivity. Intron length (i.e. longer introns) is a negative predictor of differential IR levels. ROC curve (bottom-right) depicts the intron-centric logistic regression model performance (AUC = 0.69).

**Supplemental Figure 7.**
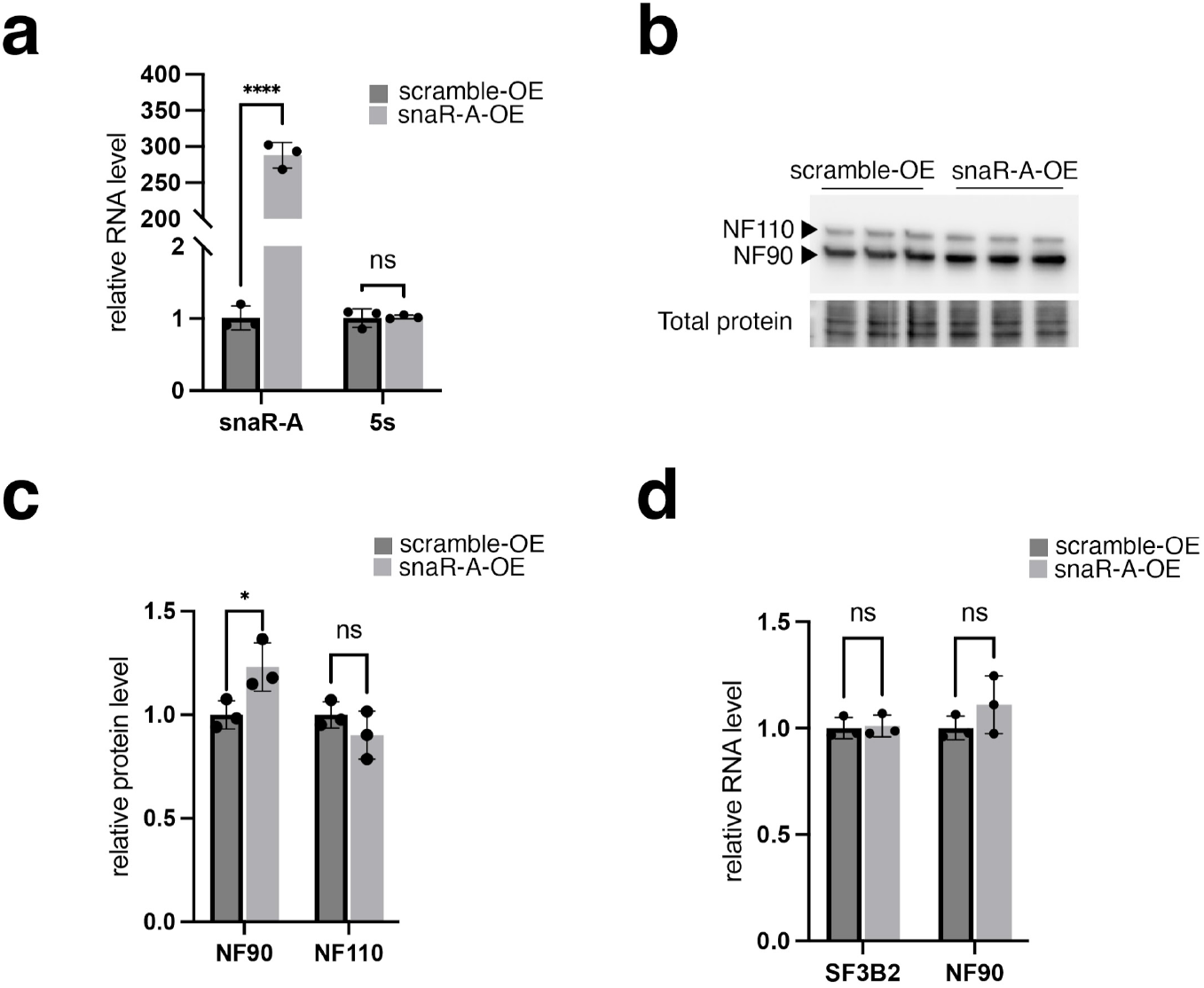
Overexpression of snaR-A does not disrupt ILF3 isoforms NF90 and NF110, nor SF3B2 RNA levels. (**a**) RT-qPCR analysis of snaR-A and 5s rRNA levels following snaR-A overexpression indicating successful overexpression of snaR-A RNA. (**b**,**c**) Immunoblots analysis (b) and quantification (c) of protein levels of NF110 and N90 following snaR-A overexpression (**d**) RT-qPCR analysis of SF3B2 and NF90 mRNA levels following snaR-A overexpression

**Supplemental Figure 8.**
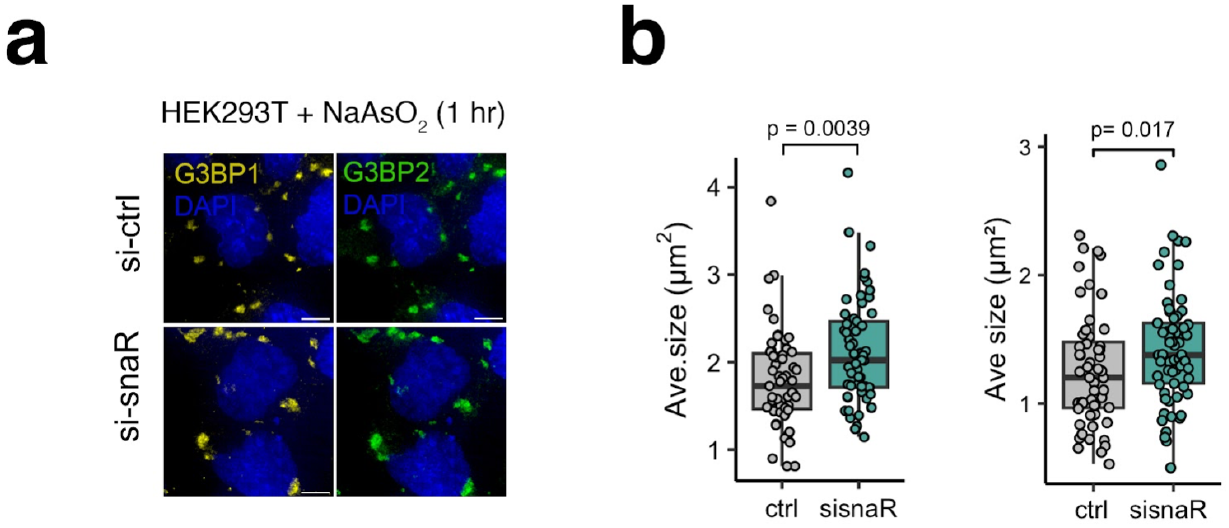
Depletion of snaR-A increases stress granule size, concomitant with increased G3BP2 protein abundance. (**a**) Immunofluorescence microscopy of stress granule marker G3BP1 (yellow) and G3BP2 (green) following snaR-A knockdown and NaAsO2 treatment (1hr) to induce stress granule. Images are maximum-intensity z projected for a 1 μm section. Scale bars, 5 *µ*m (**b**) Quantifications of average stress granule size using G3BP1 as an indicator for stress granules across two biological replicates.

## Supplemental Tables

**Supplemental Table 1.**
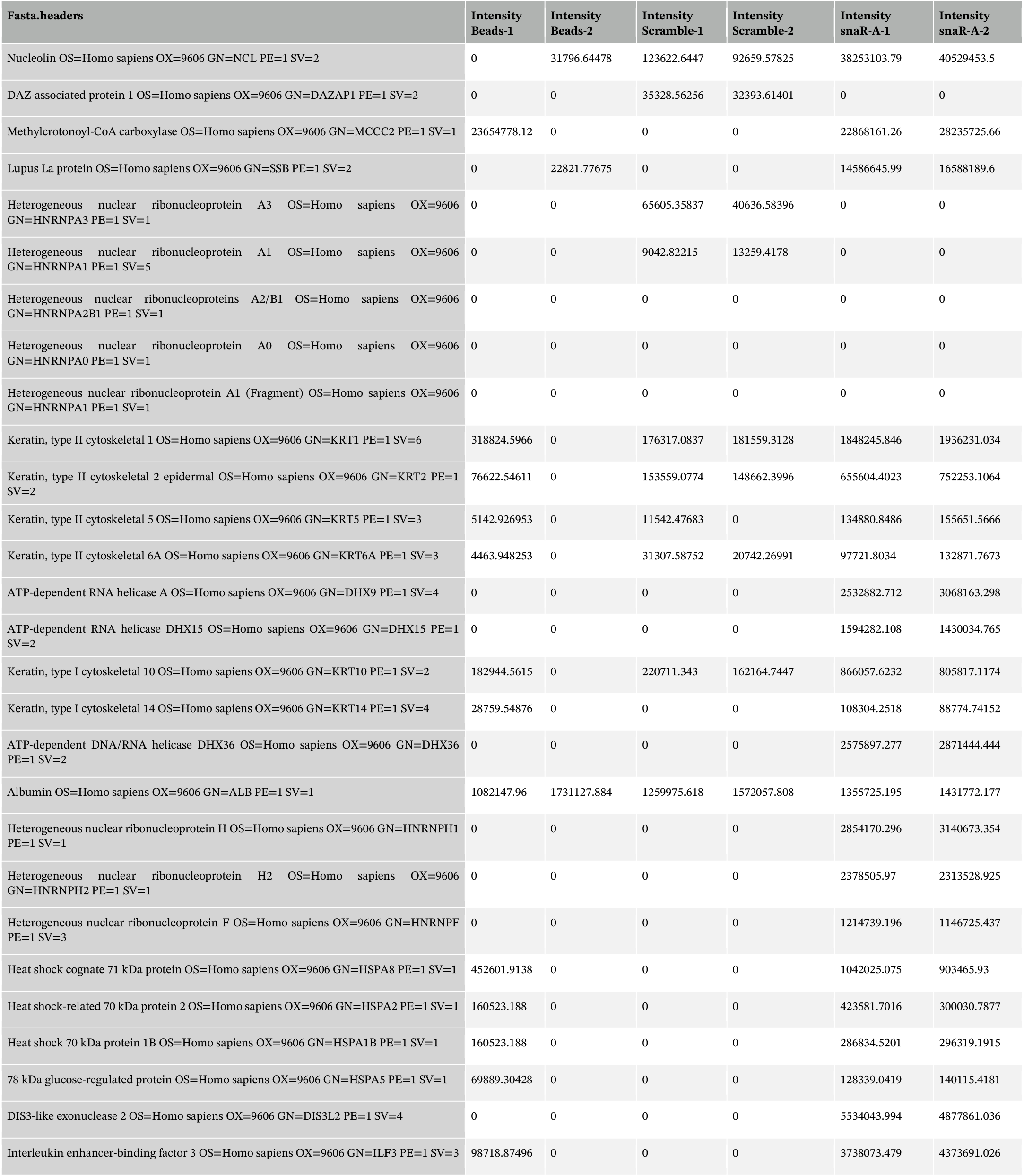
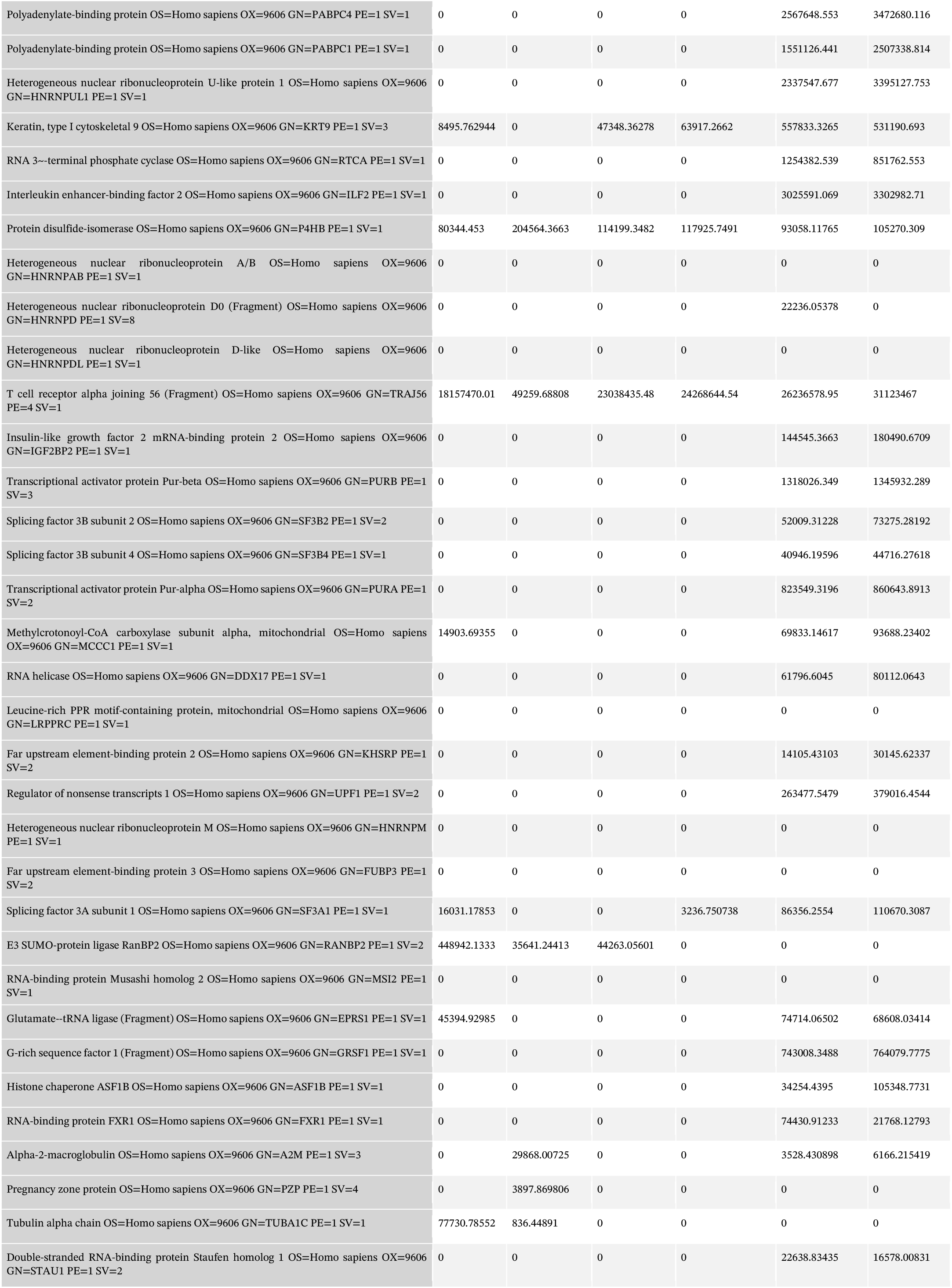
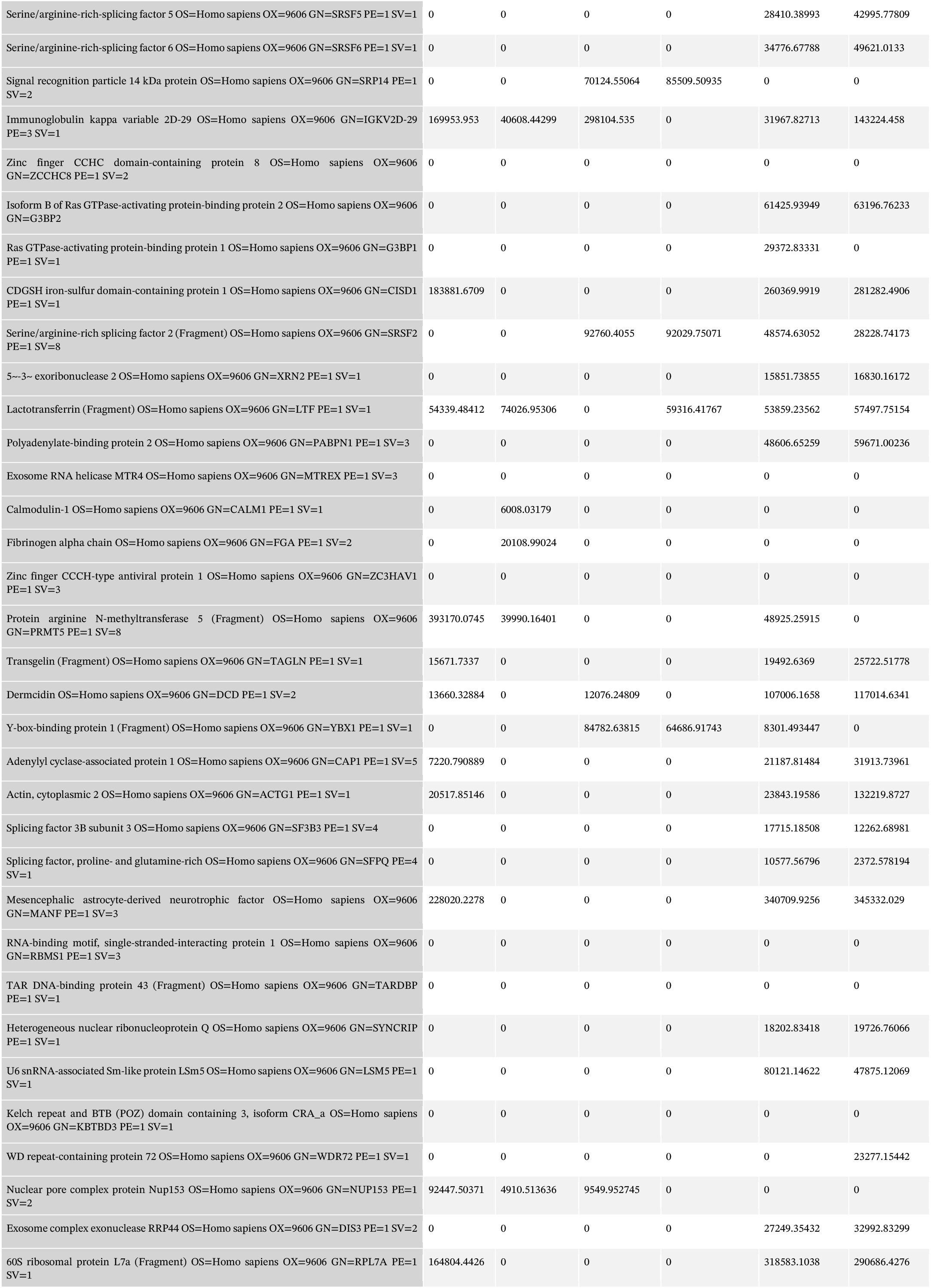
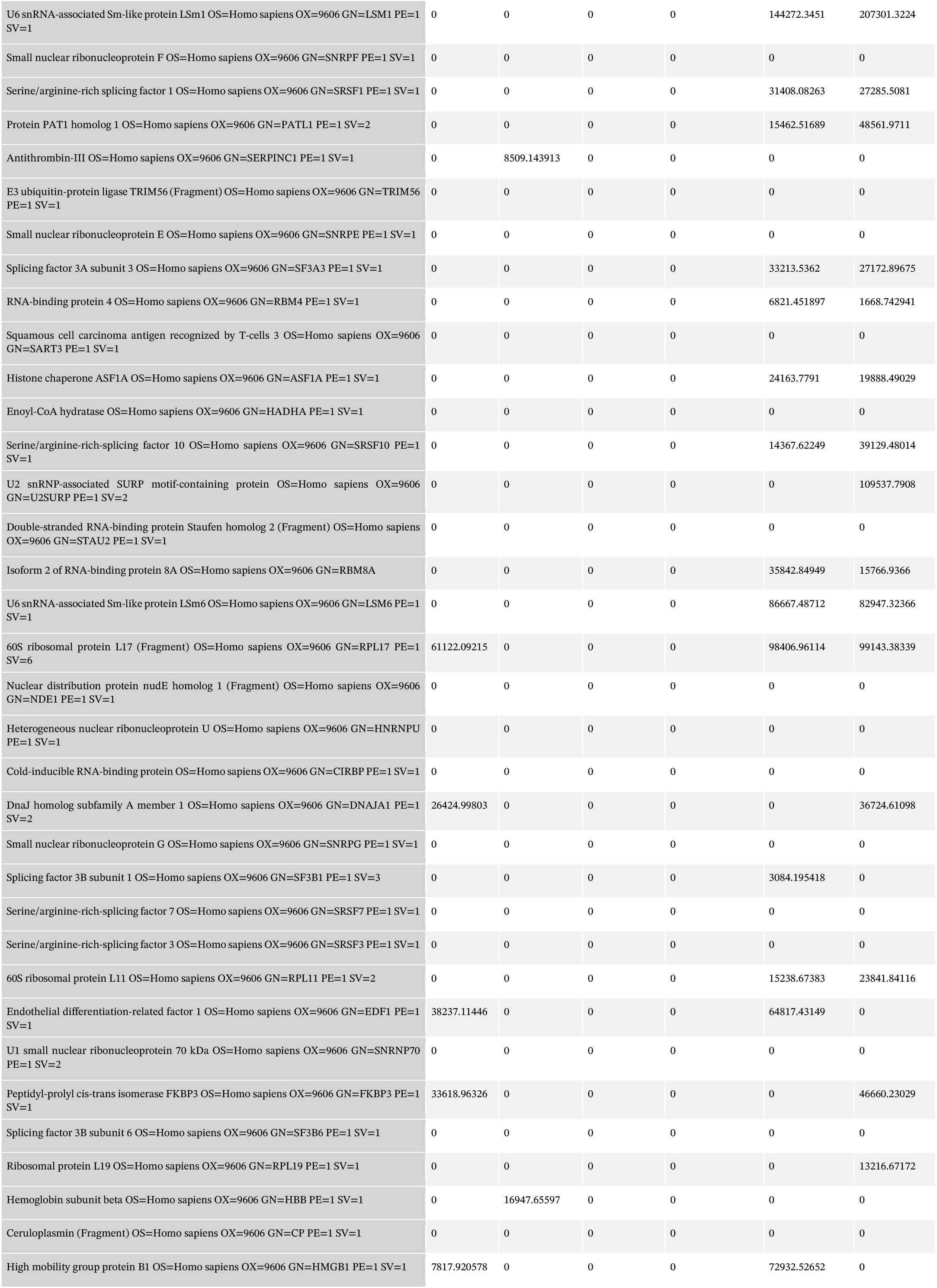
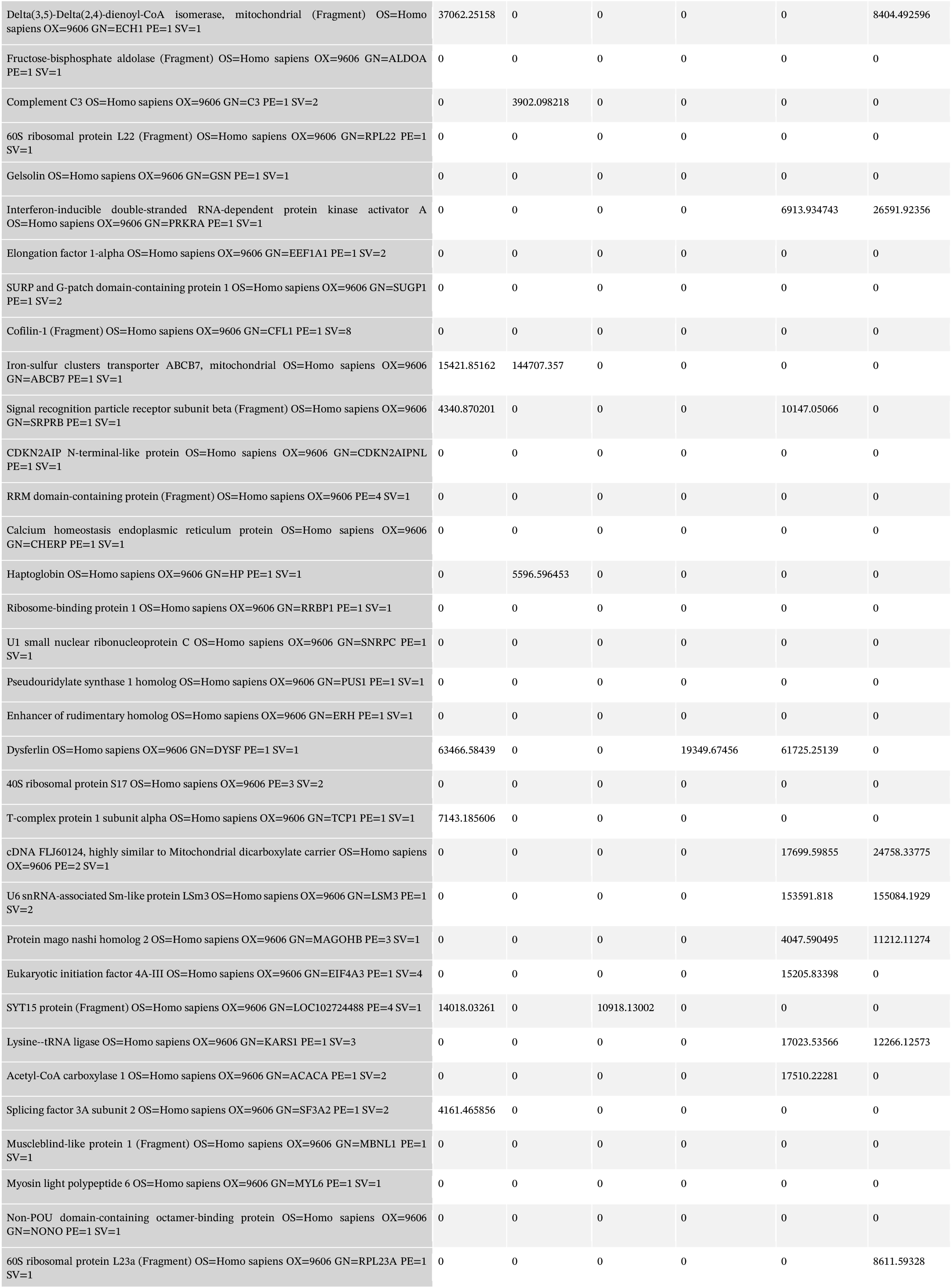
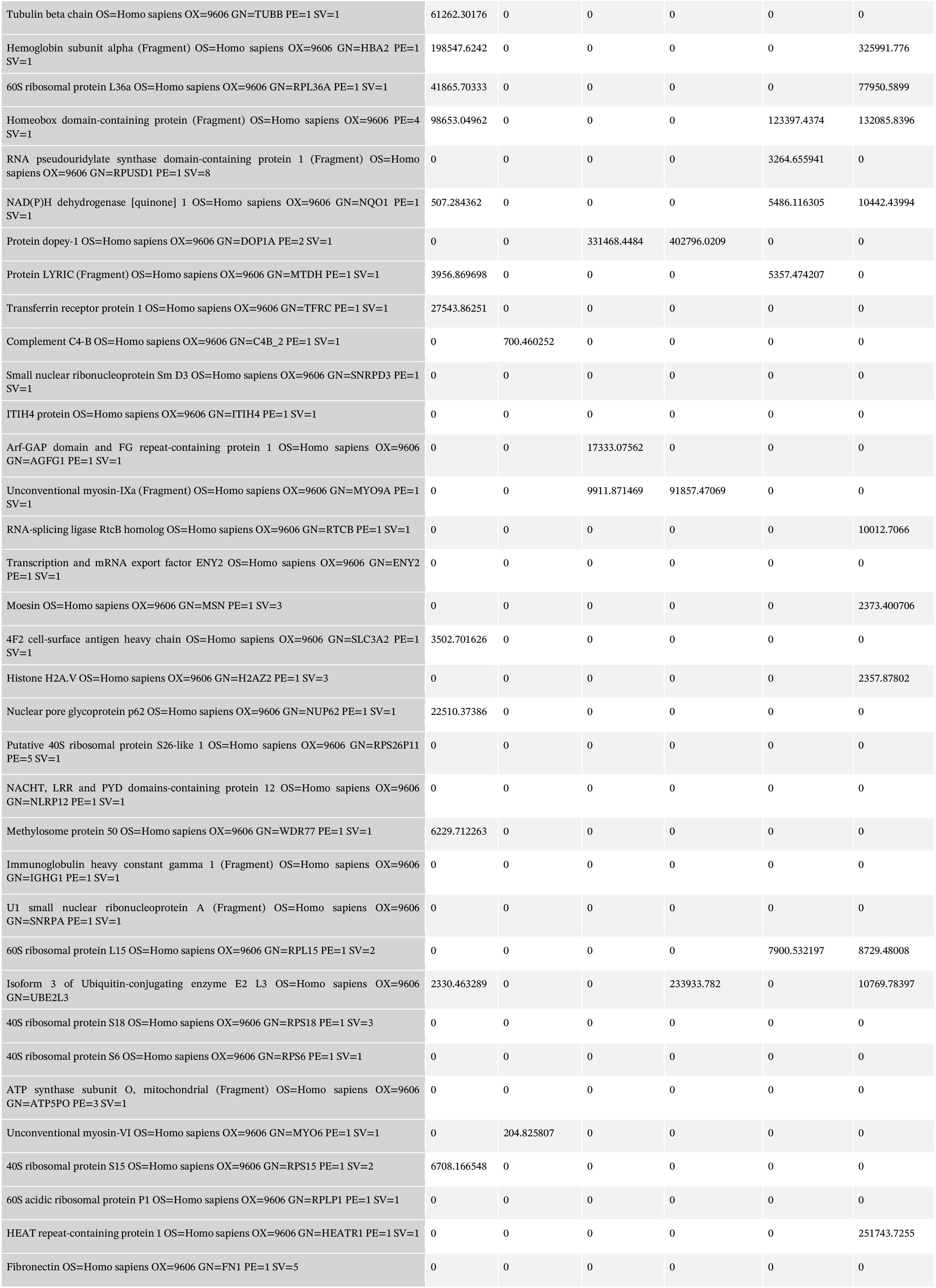
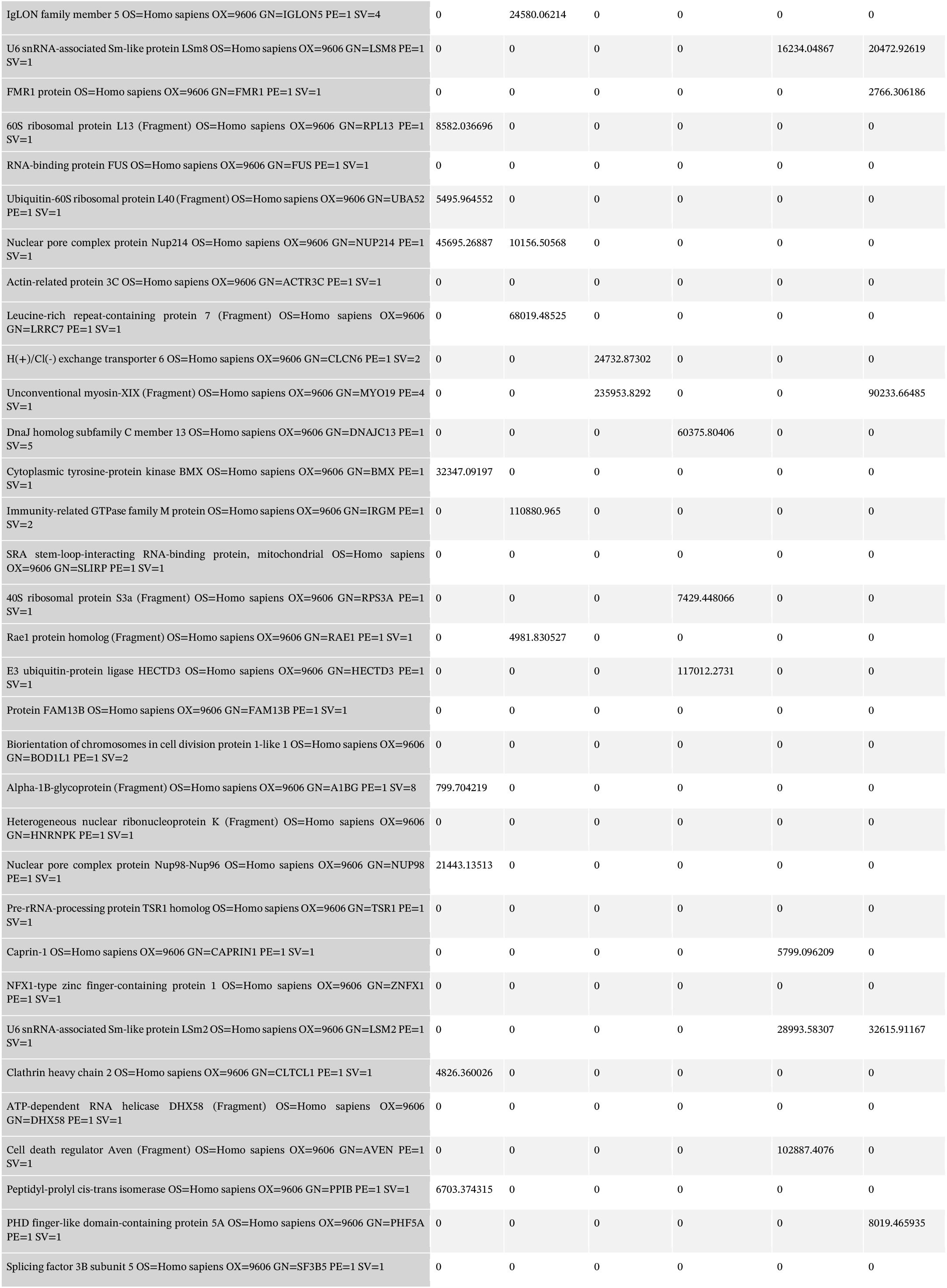
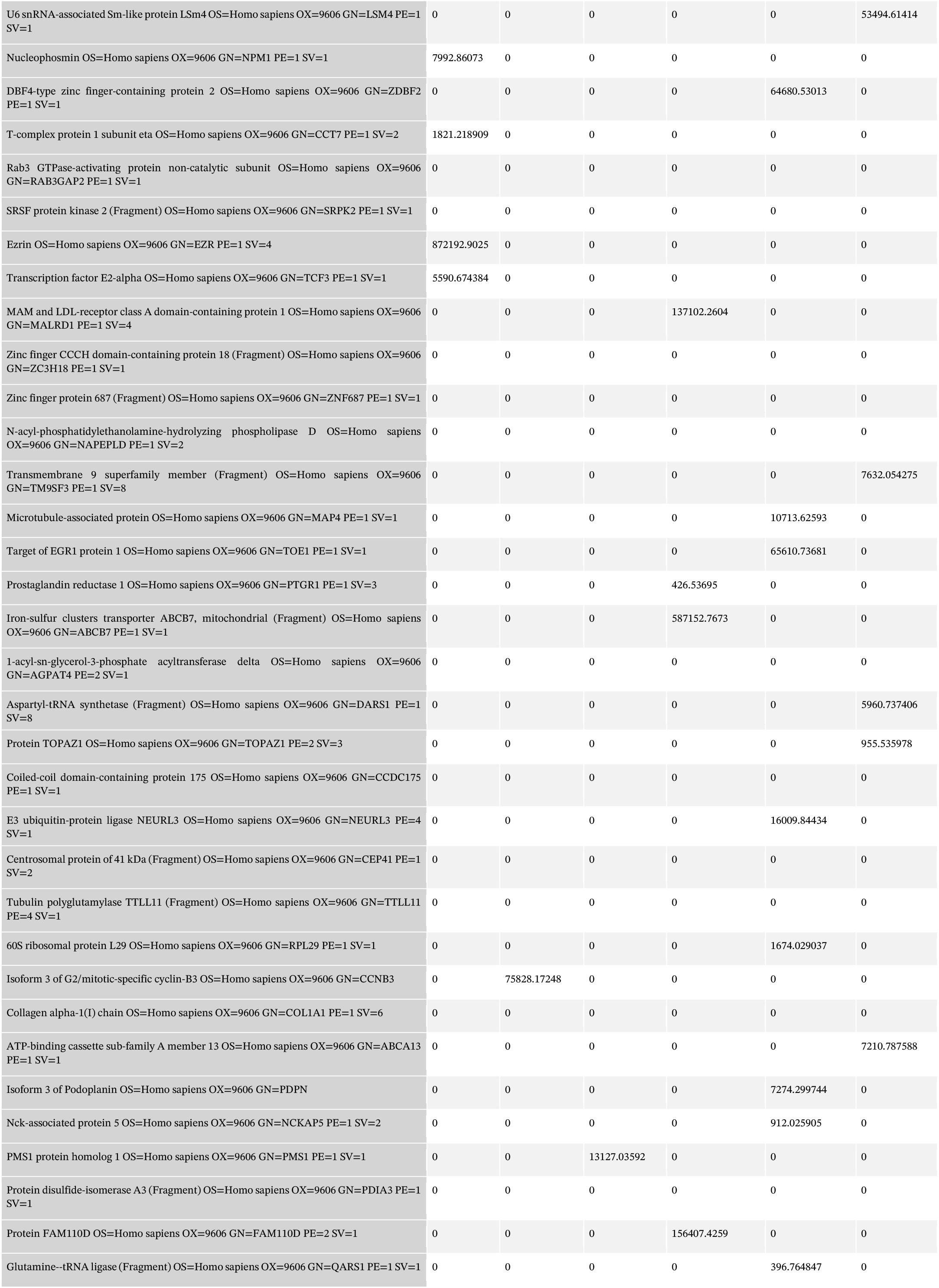
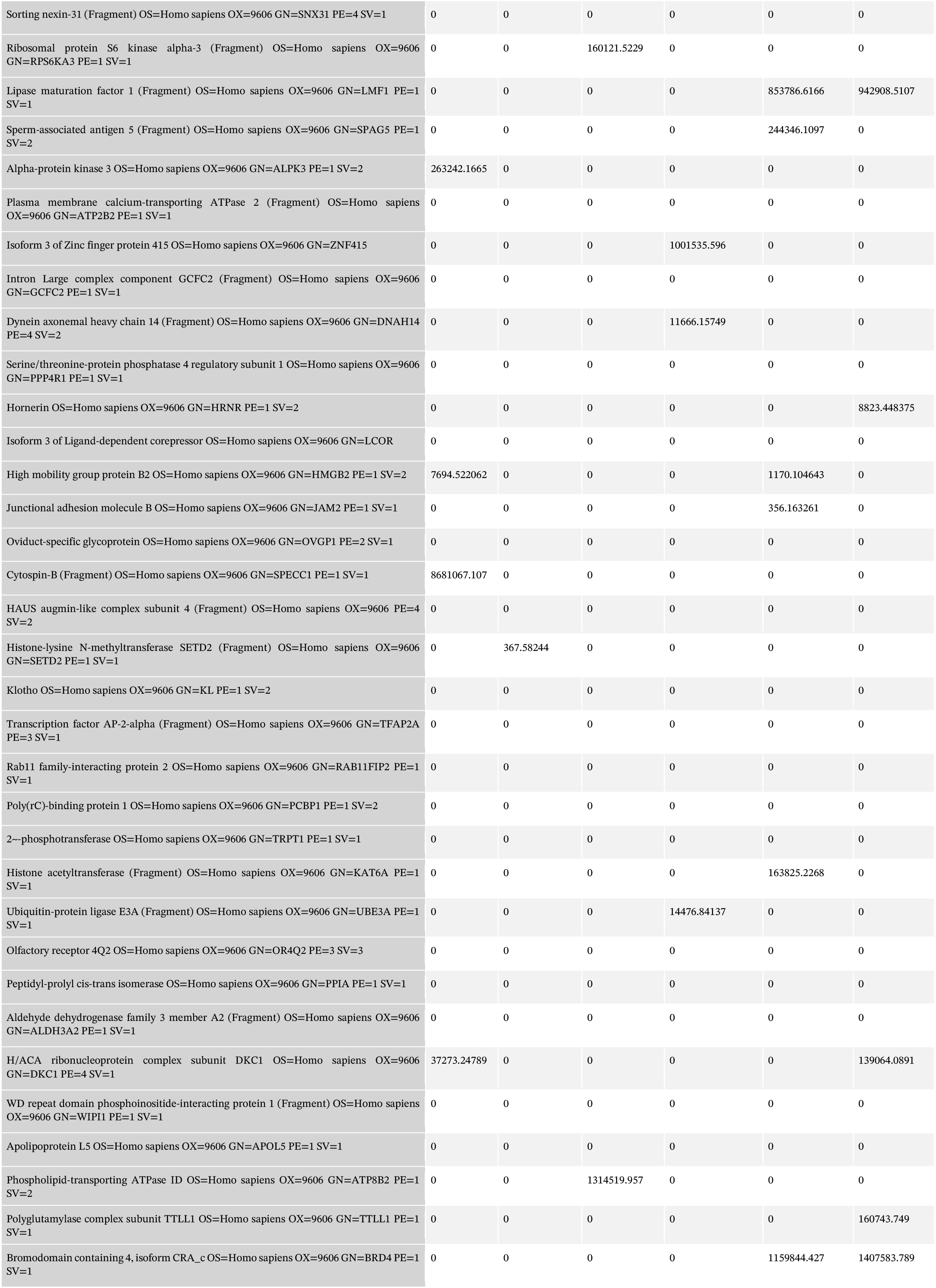
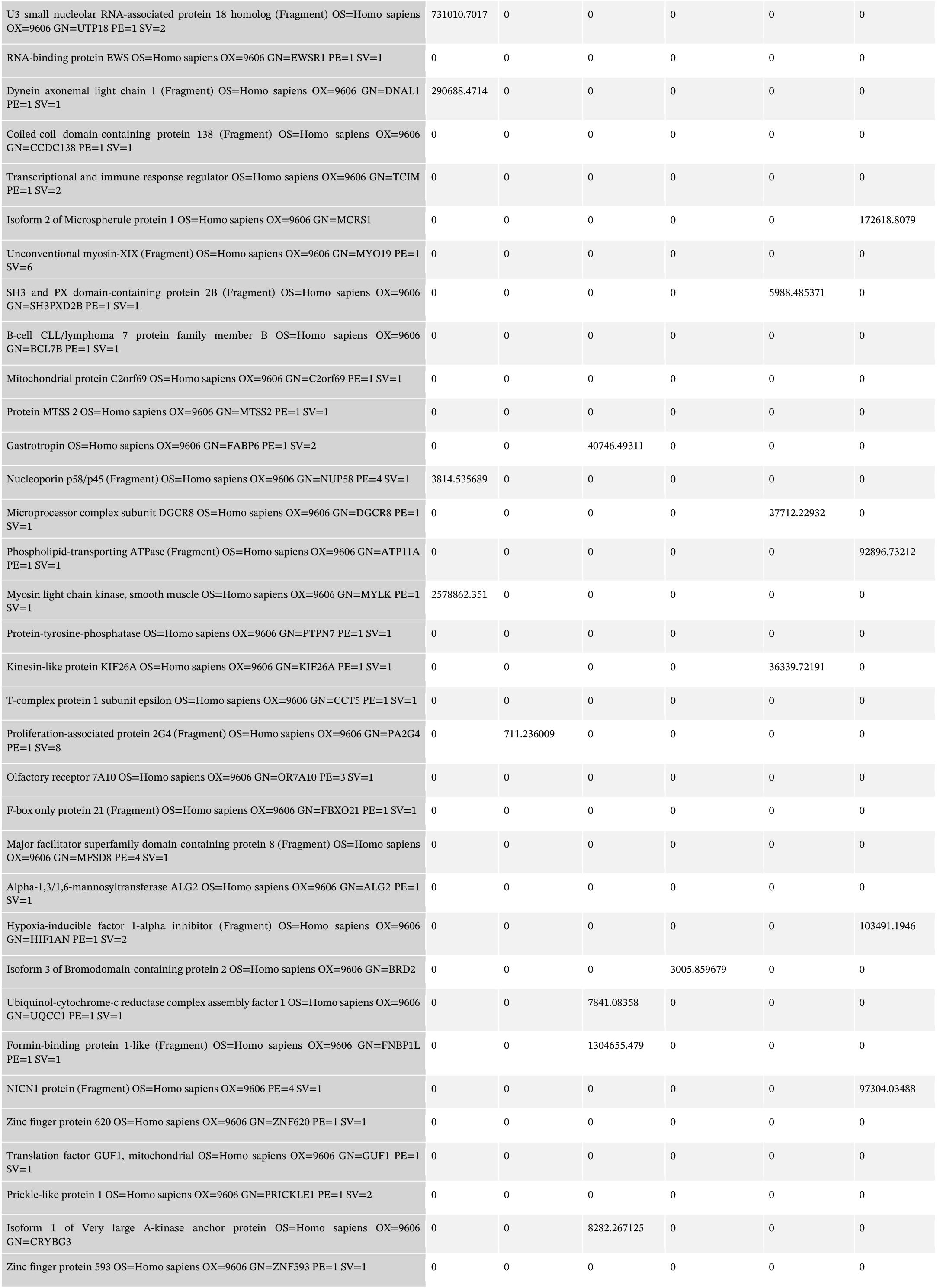
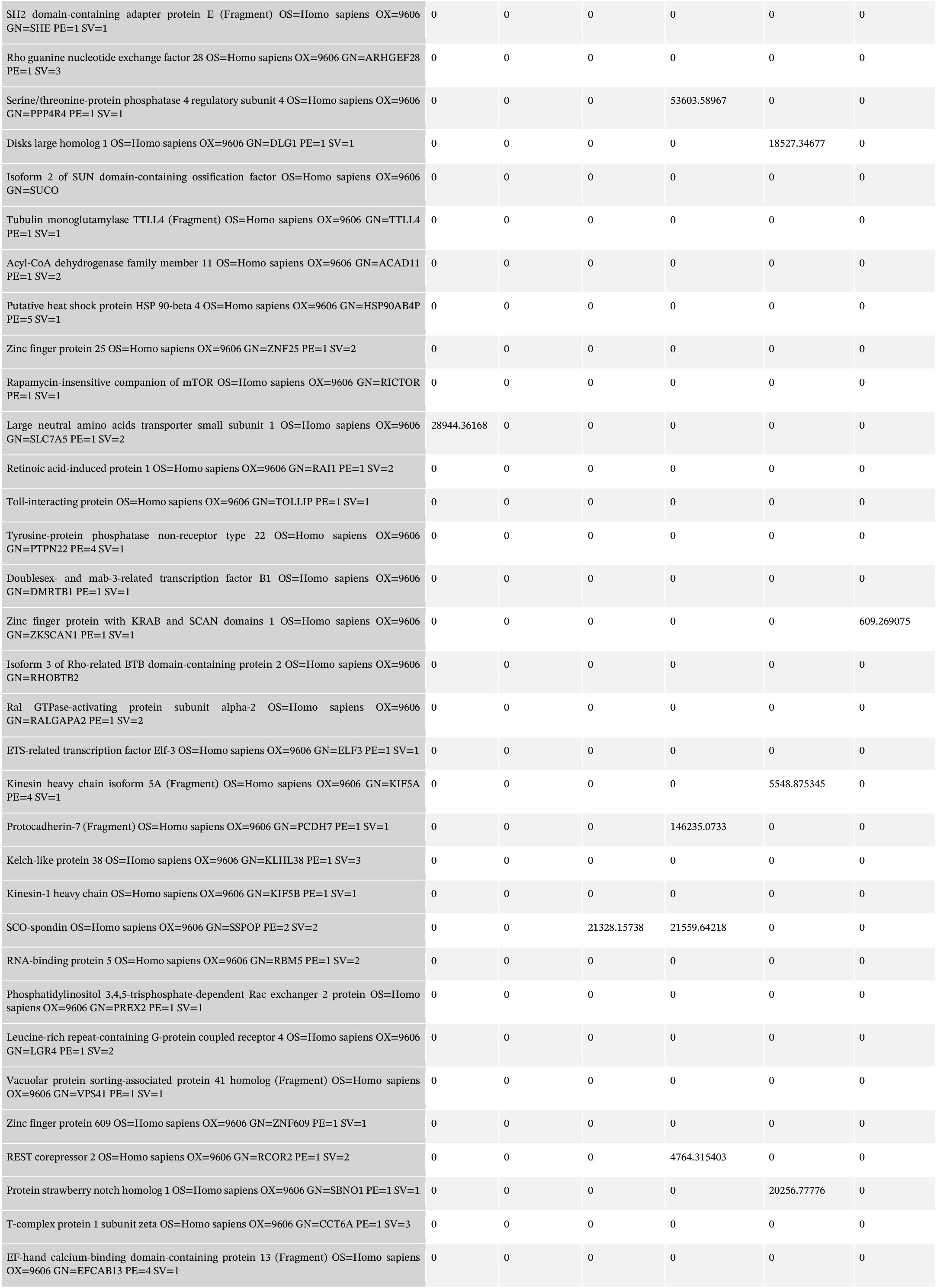
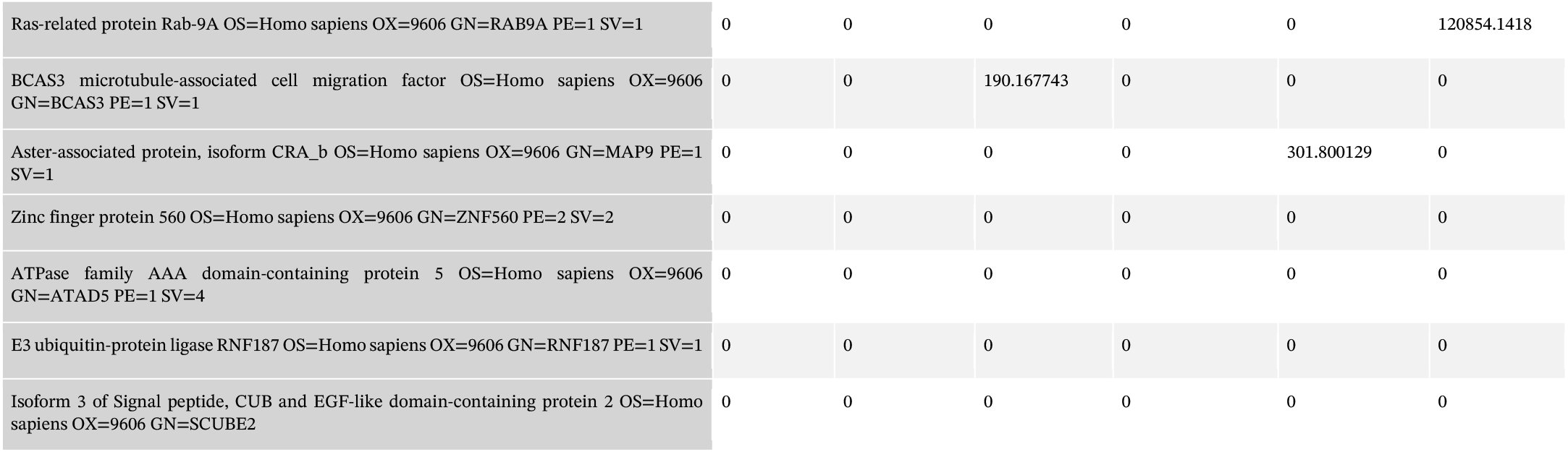
Liquid Chromatography-Tandem Mass Spectrometry analysis. (snaR-A pulldown, scramble pulldown, and beads-only control)

**Supplemental Table 2.**
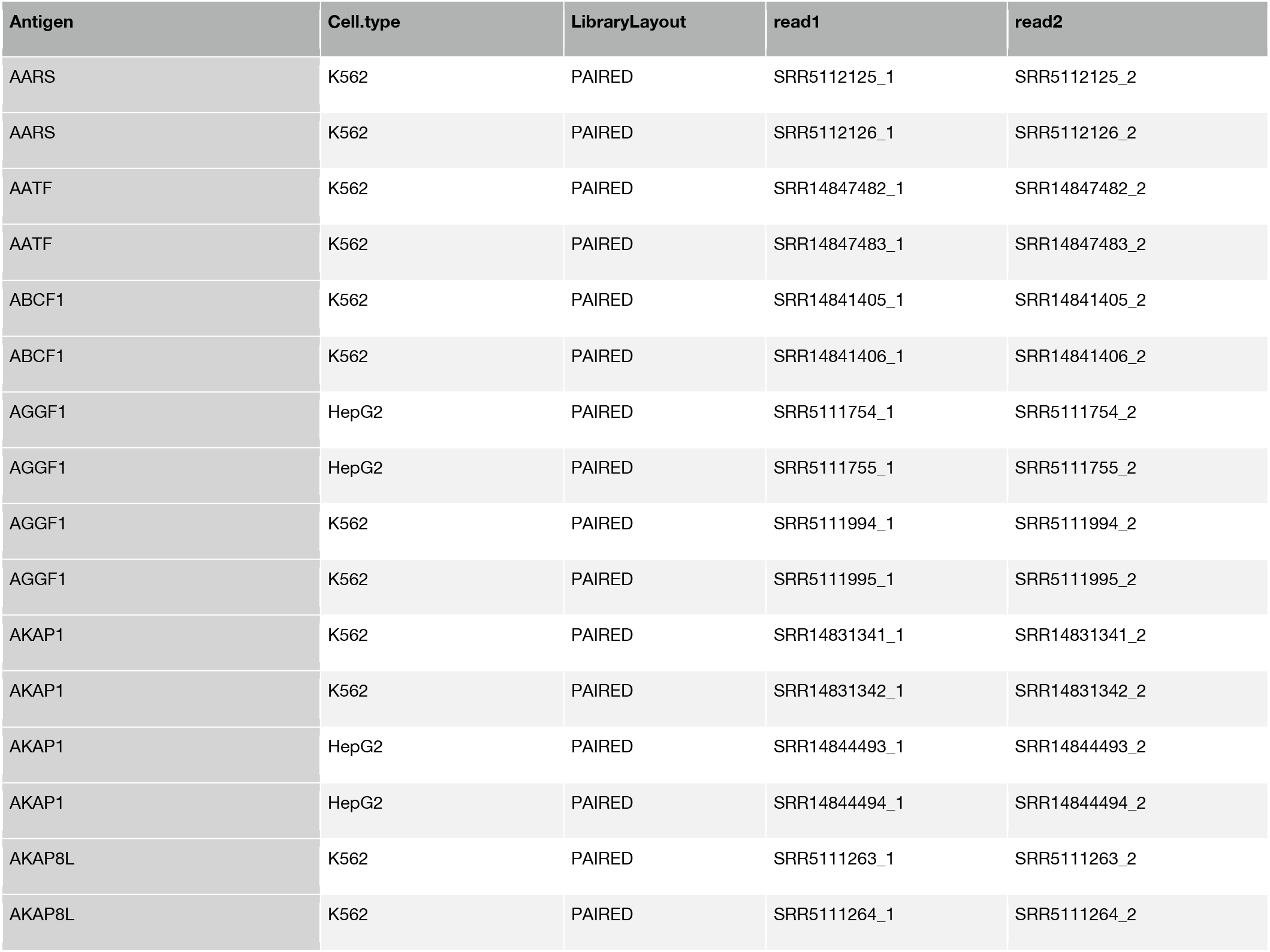
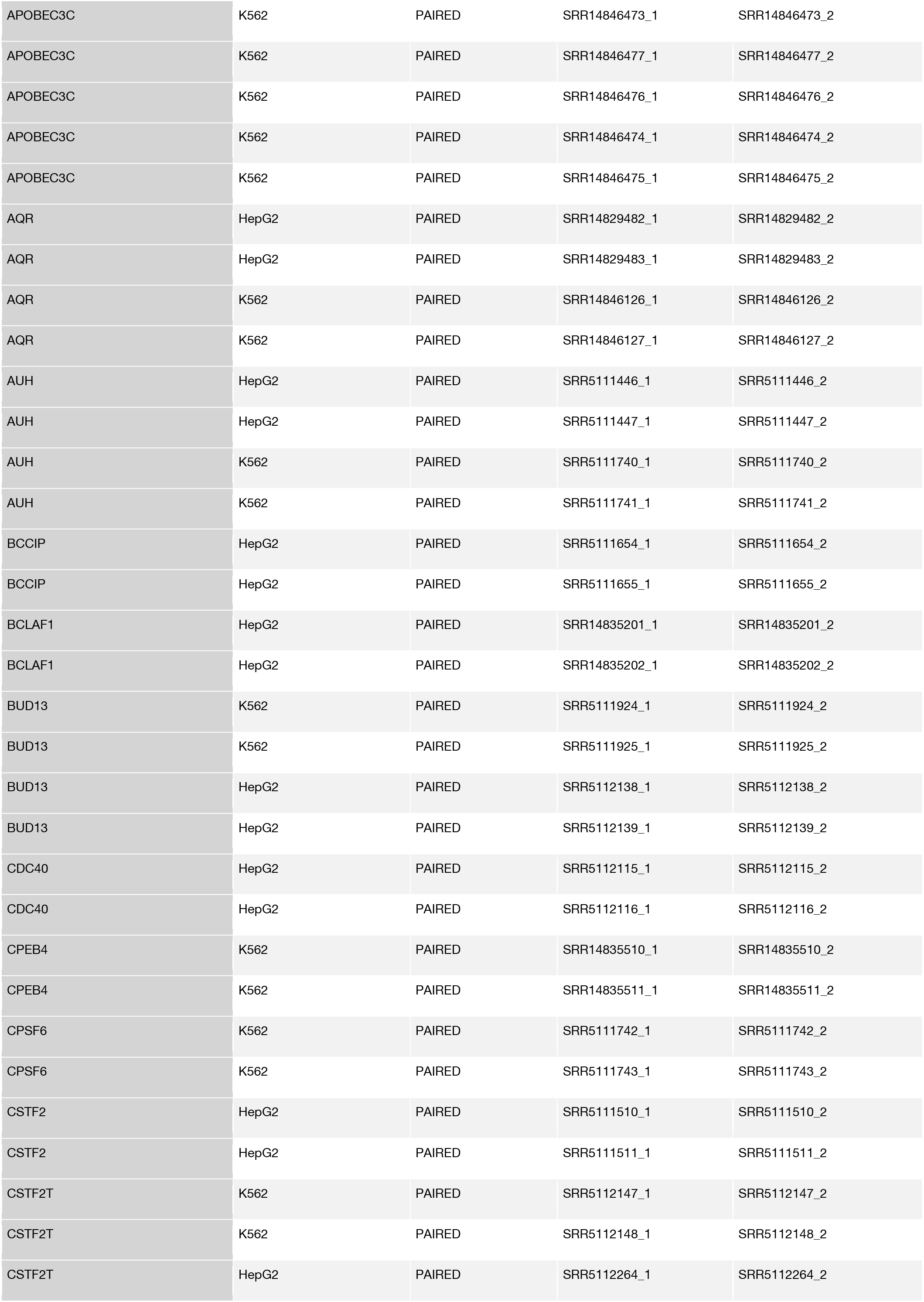
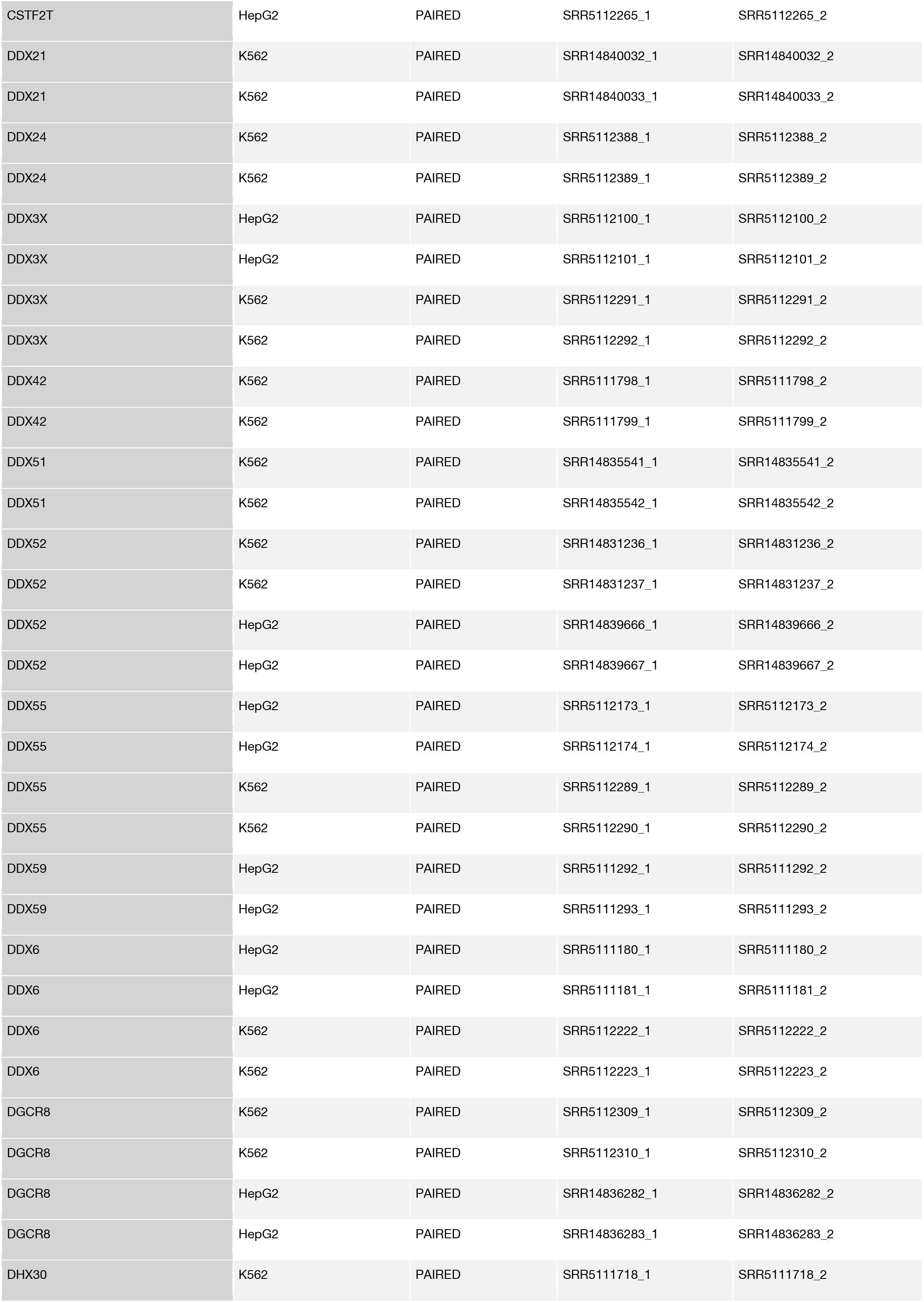
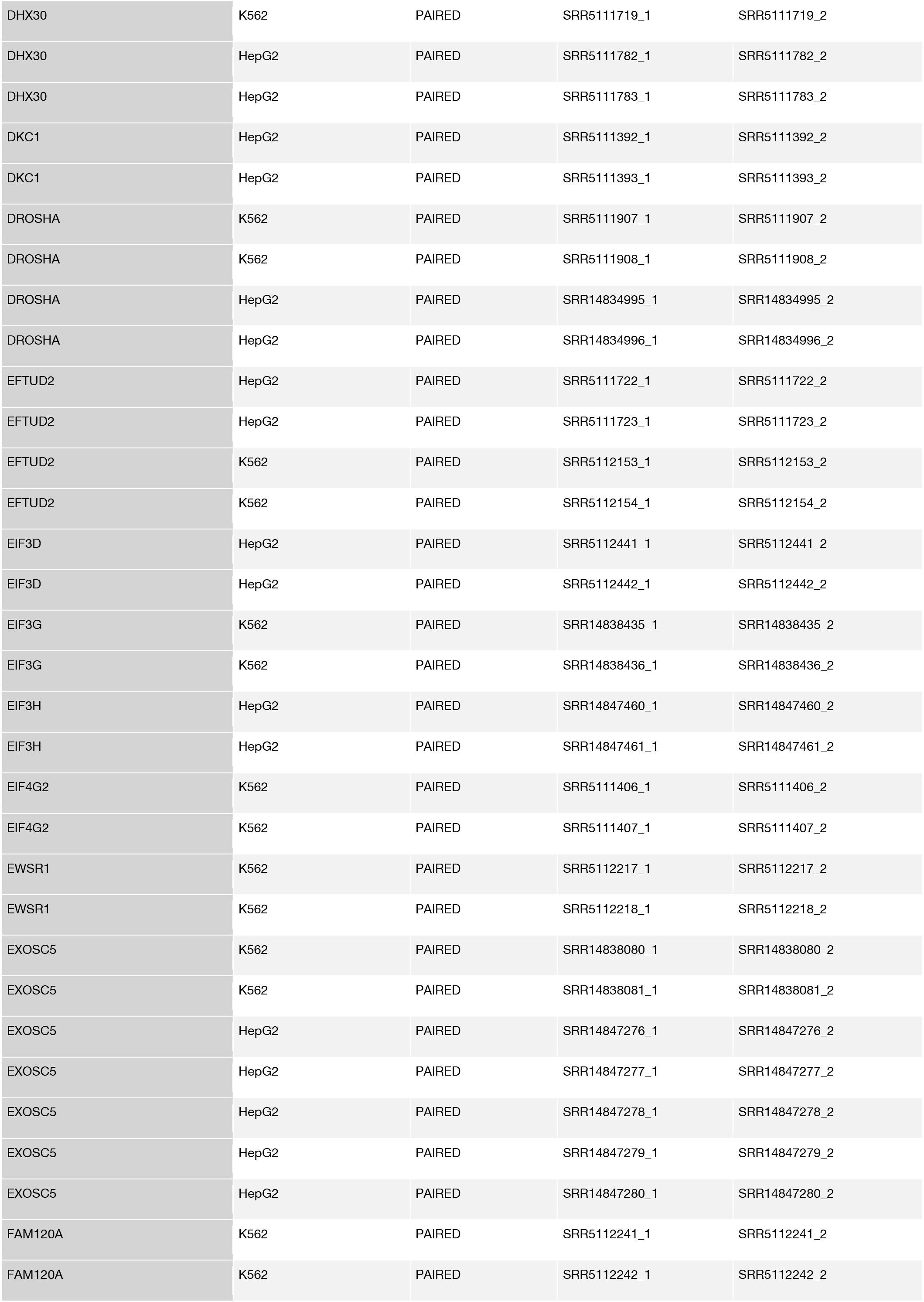
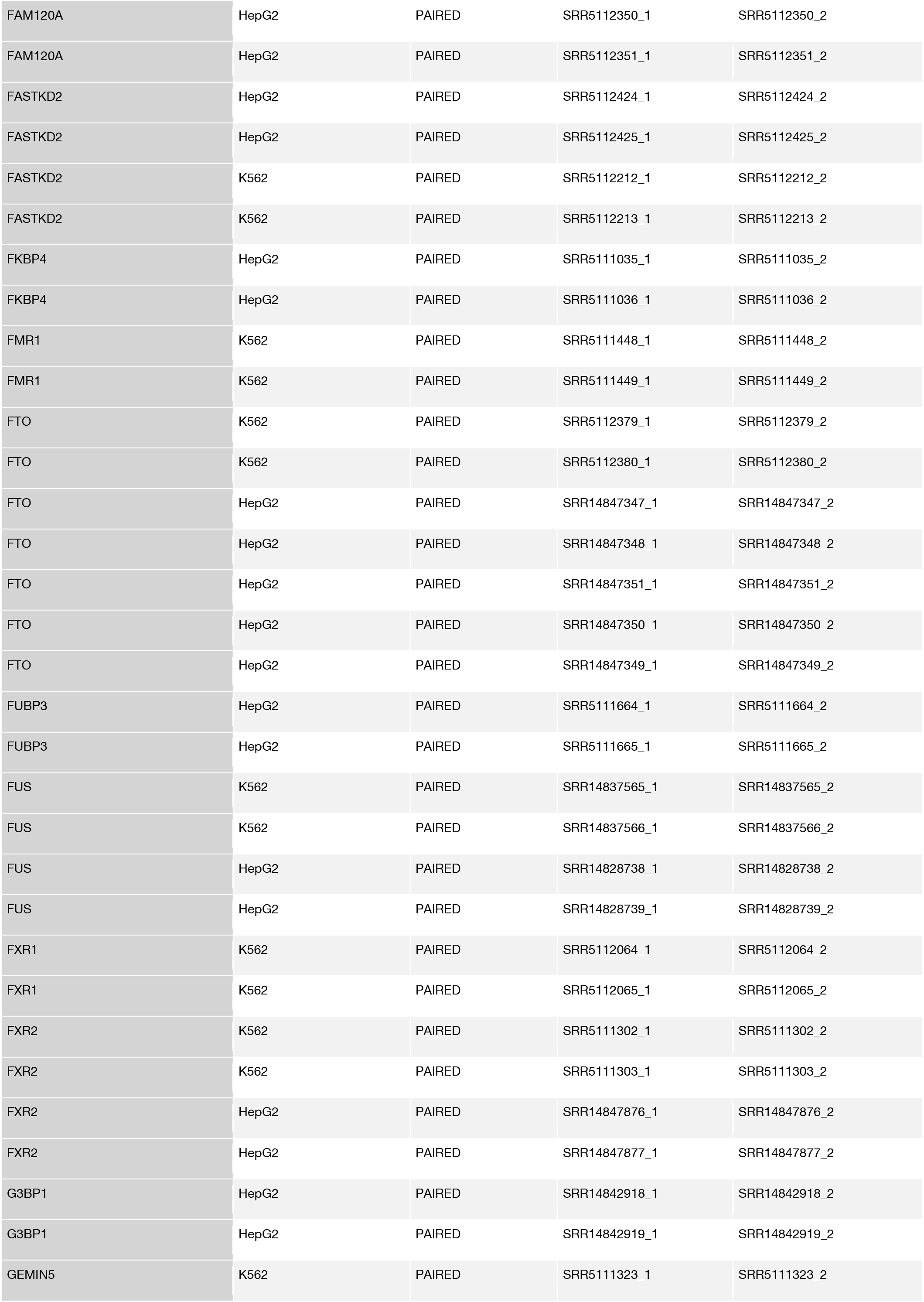
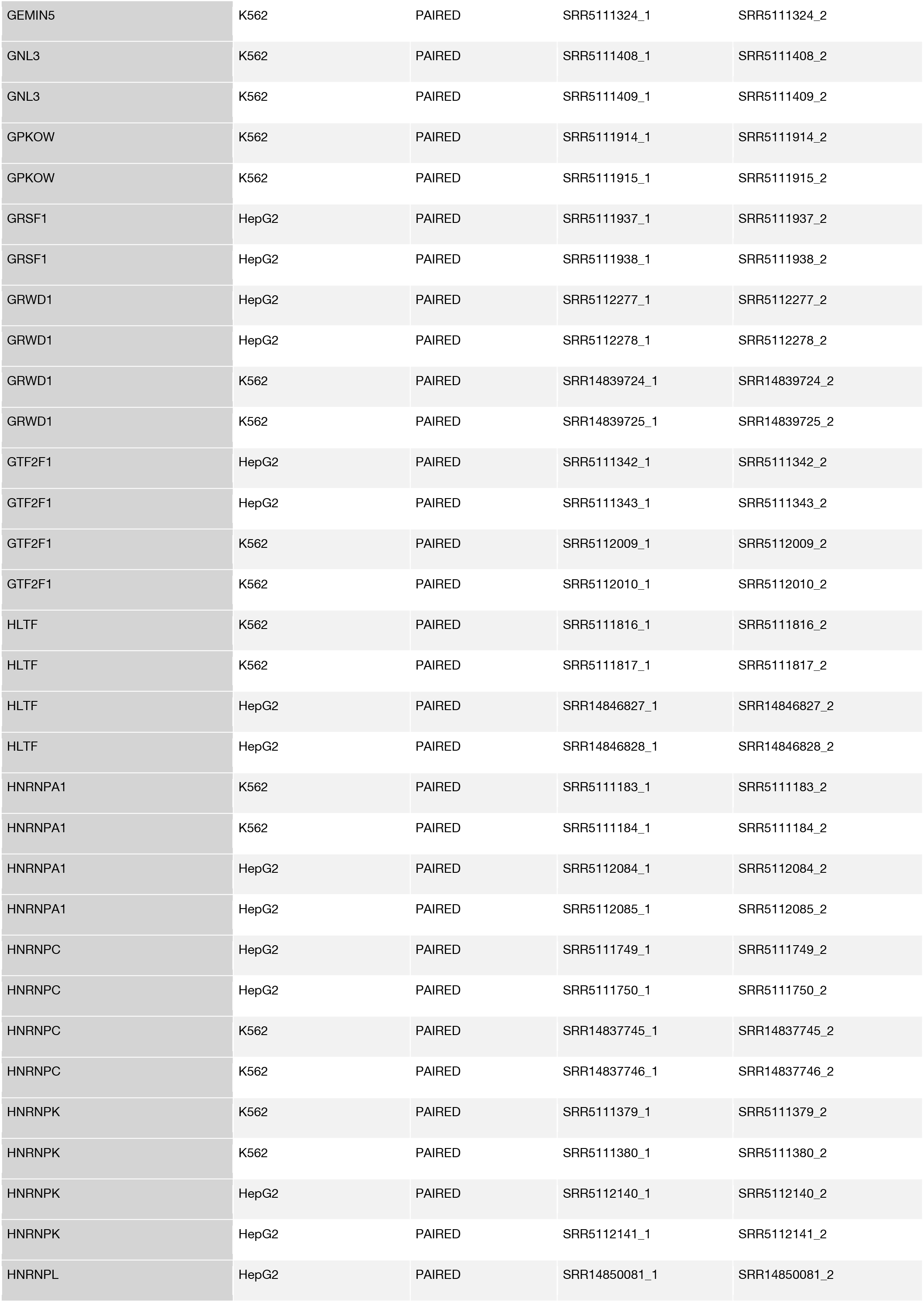
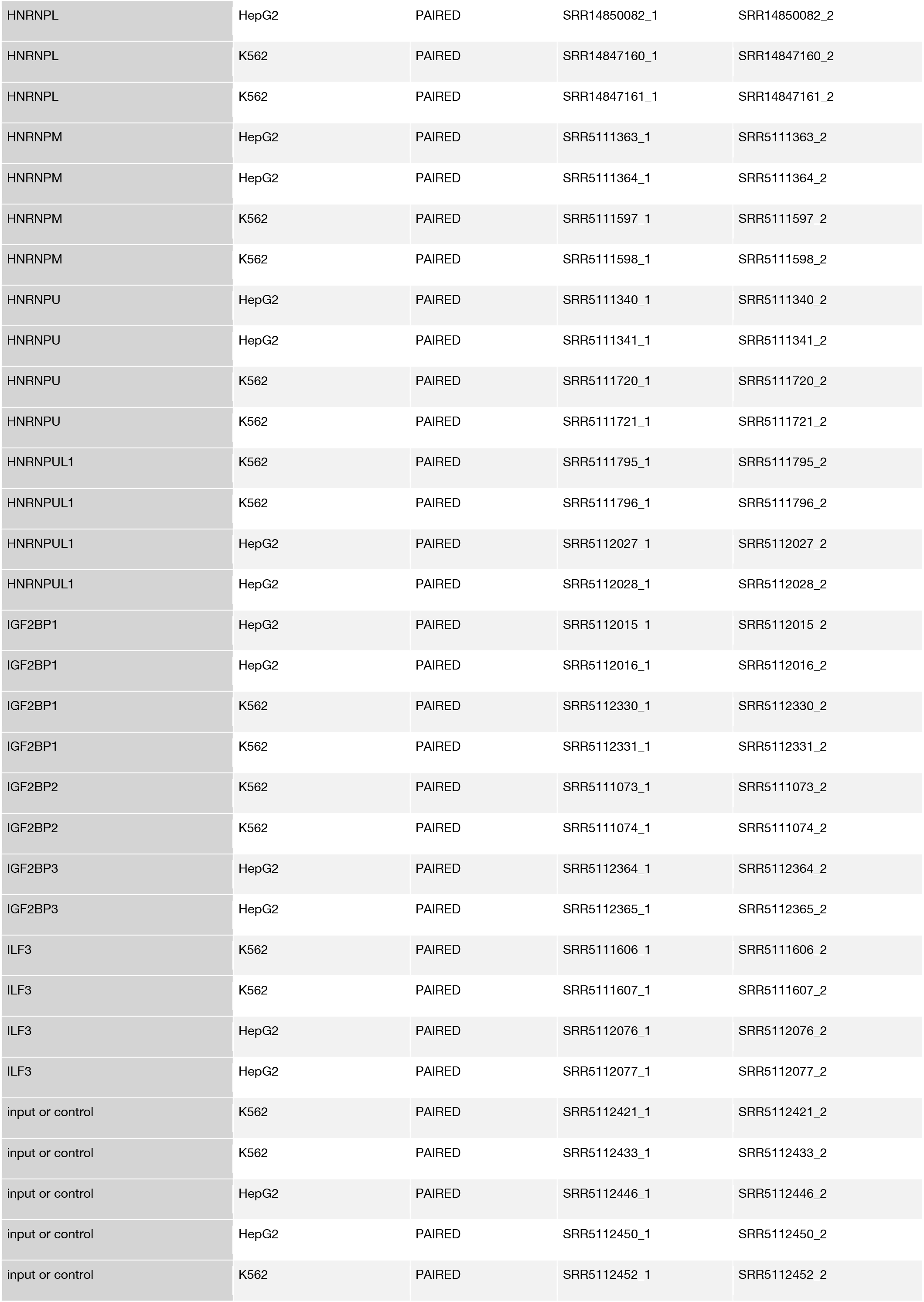
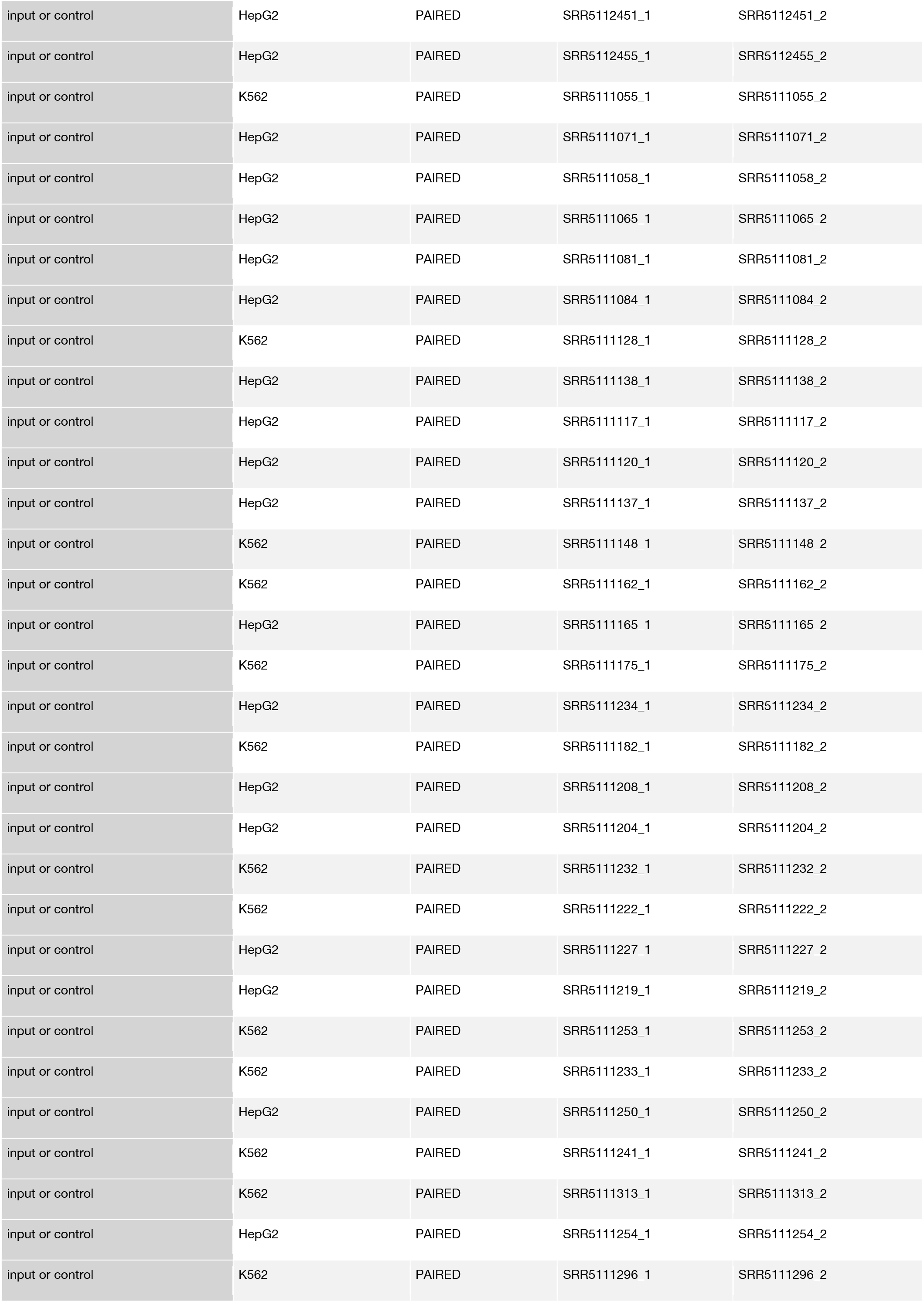
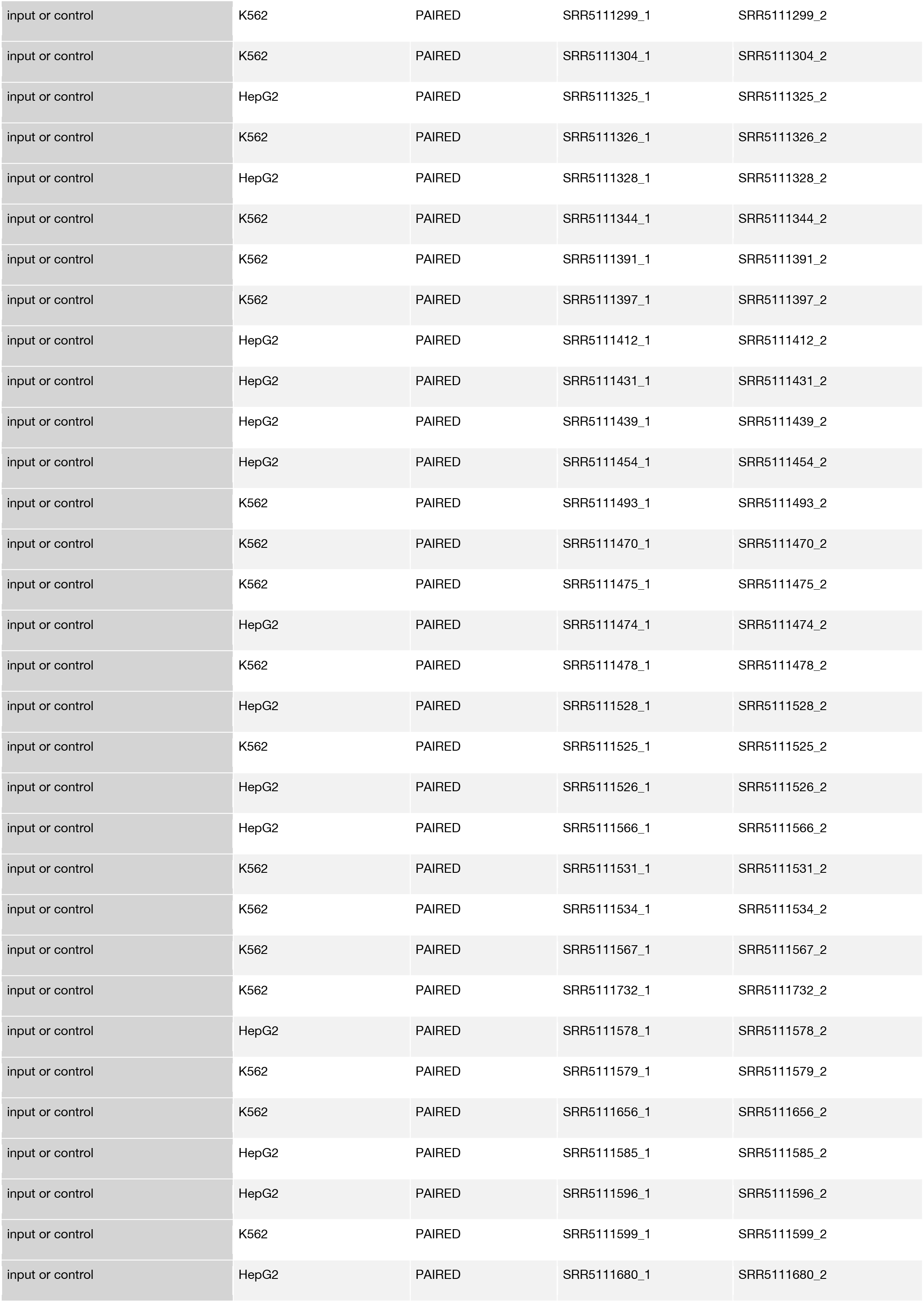
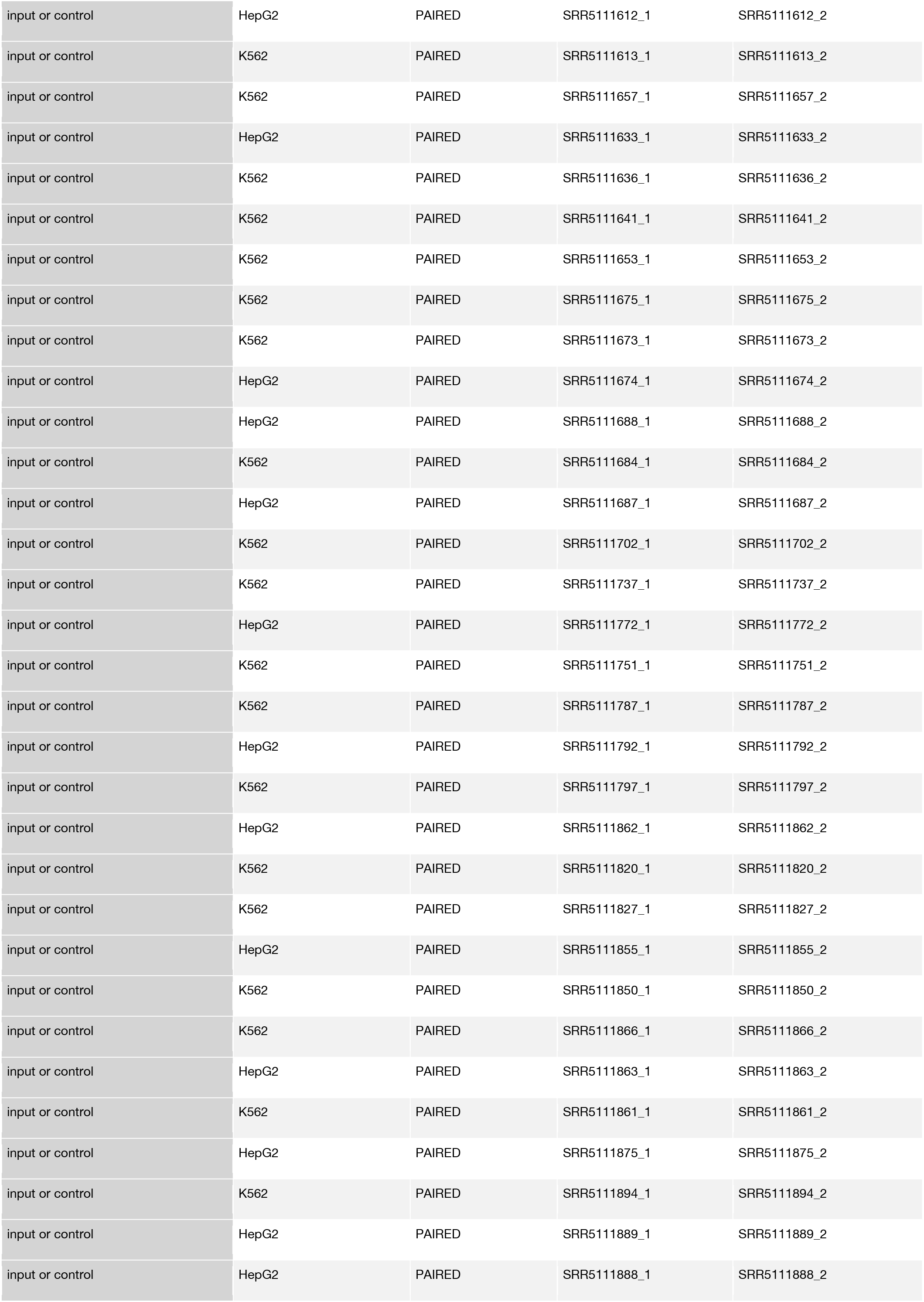
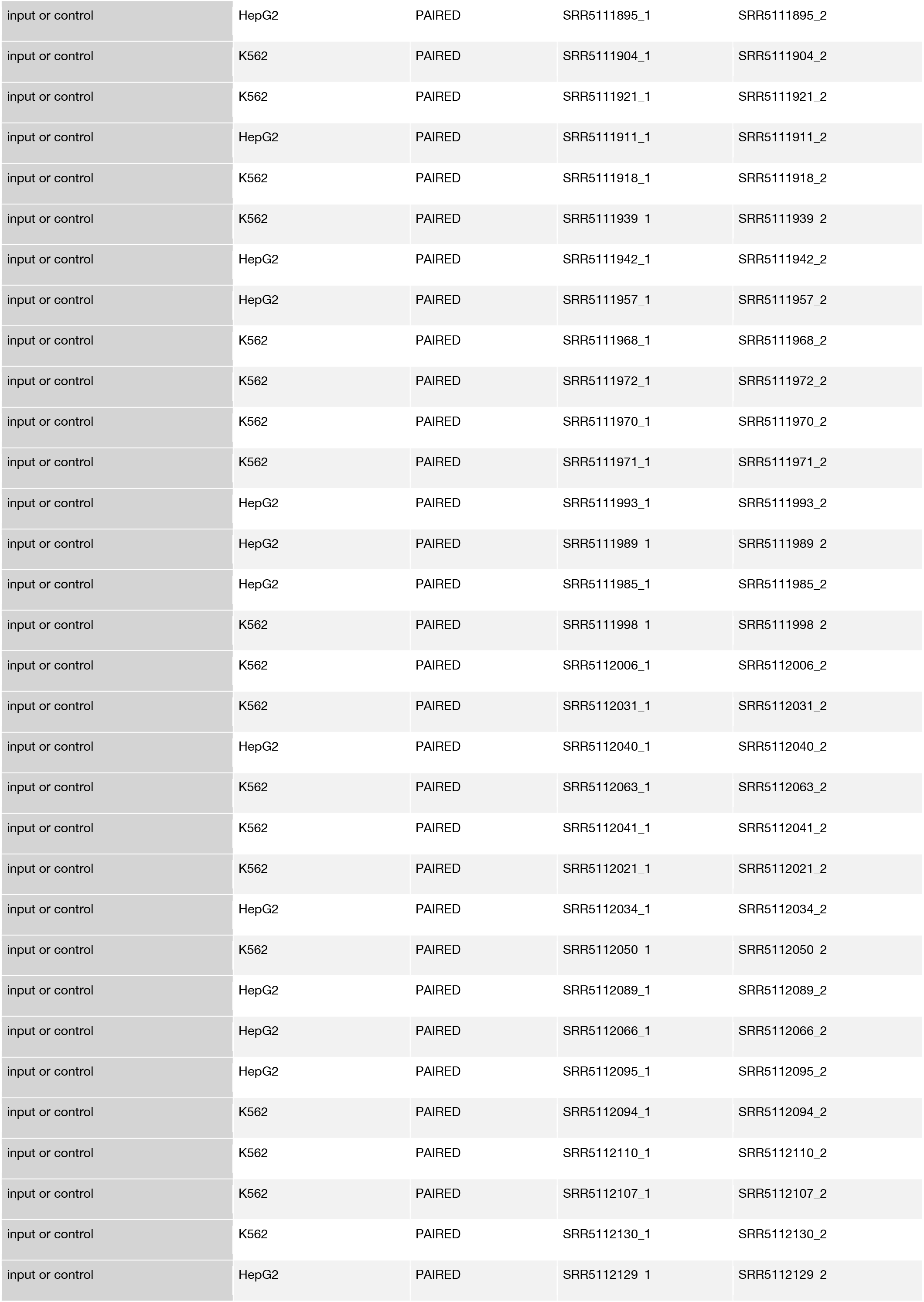
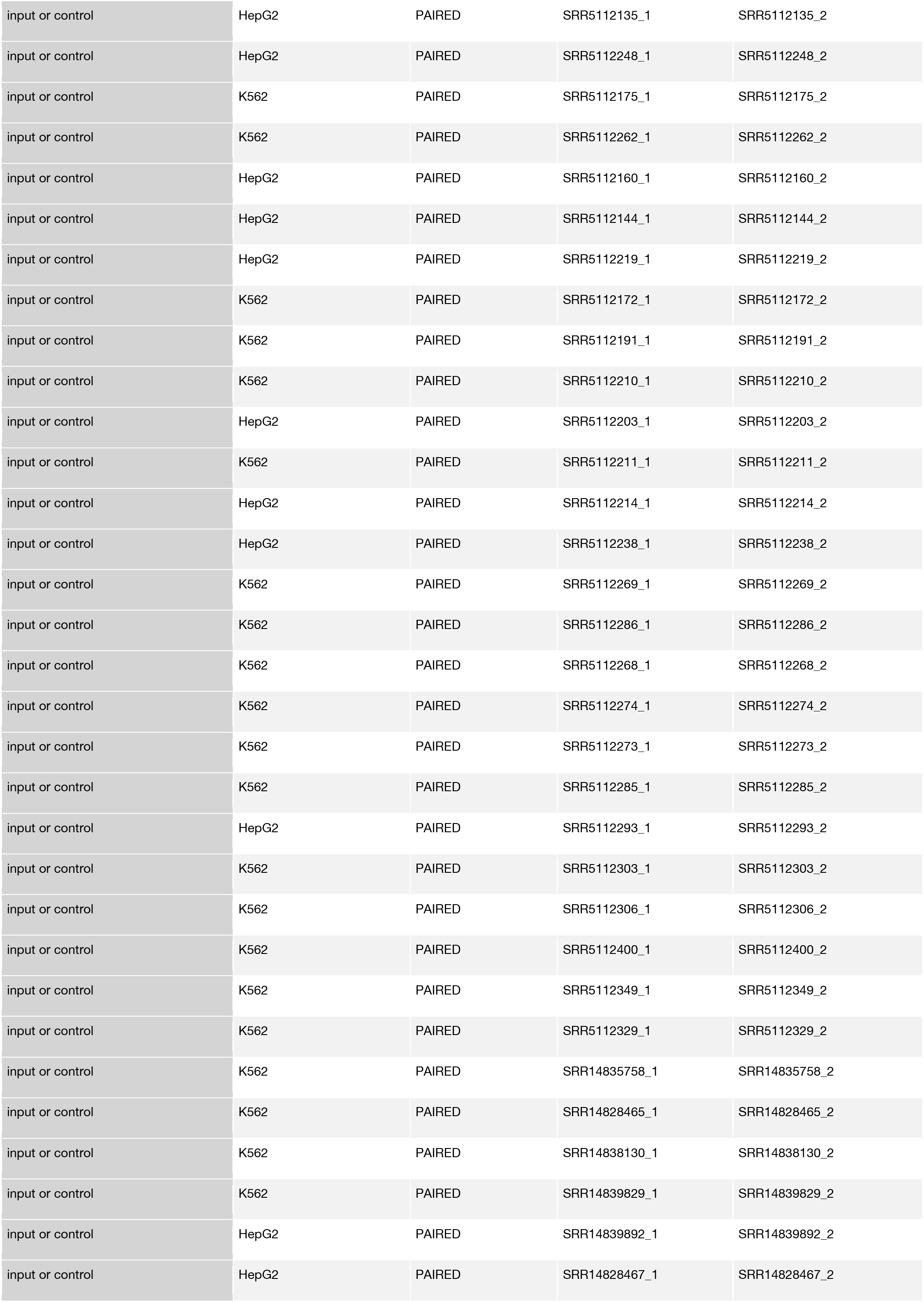
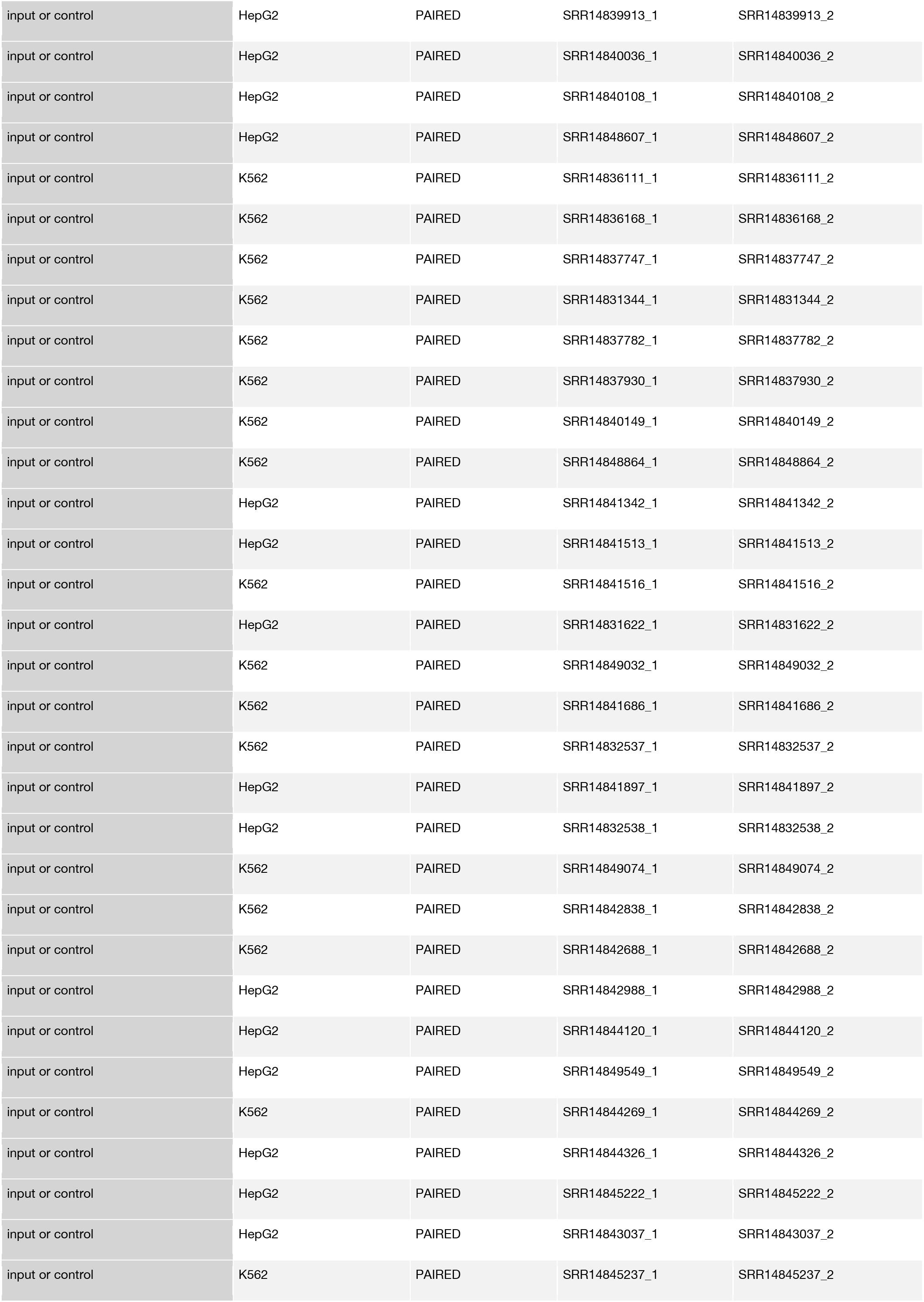
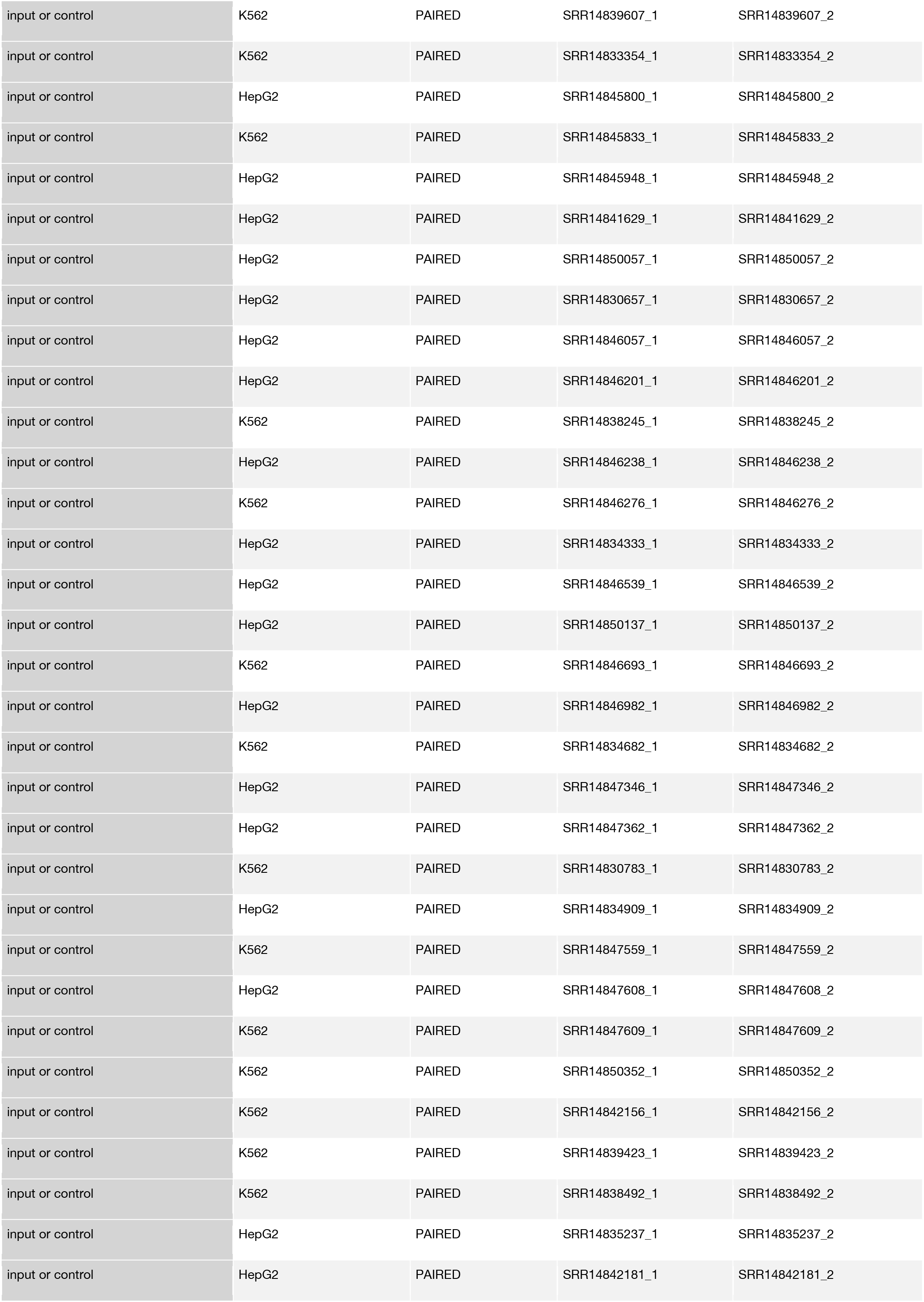
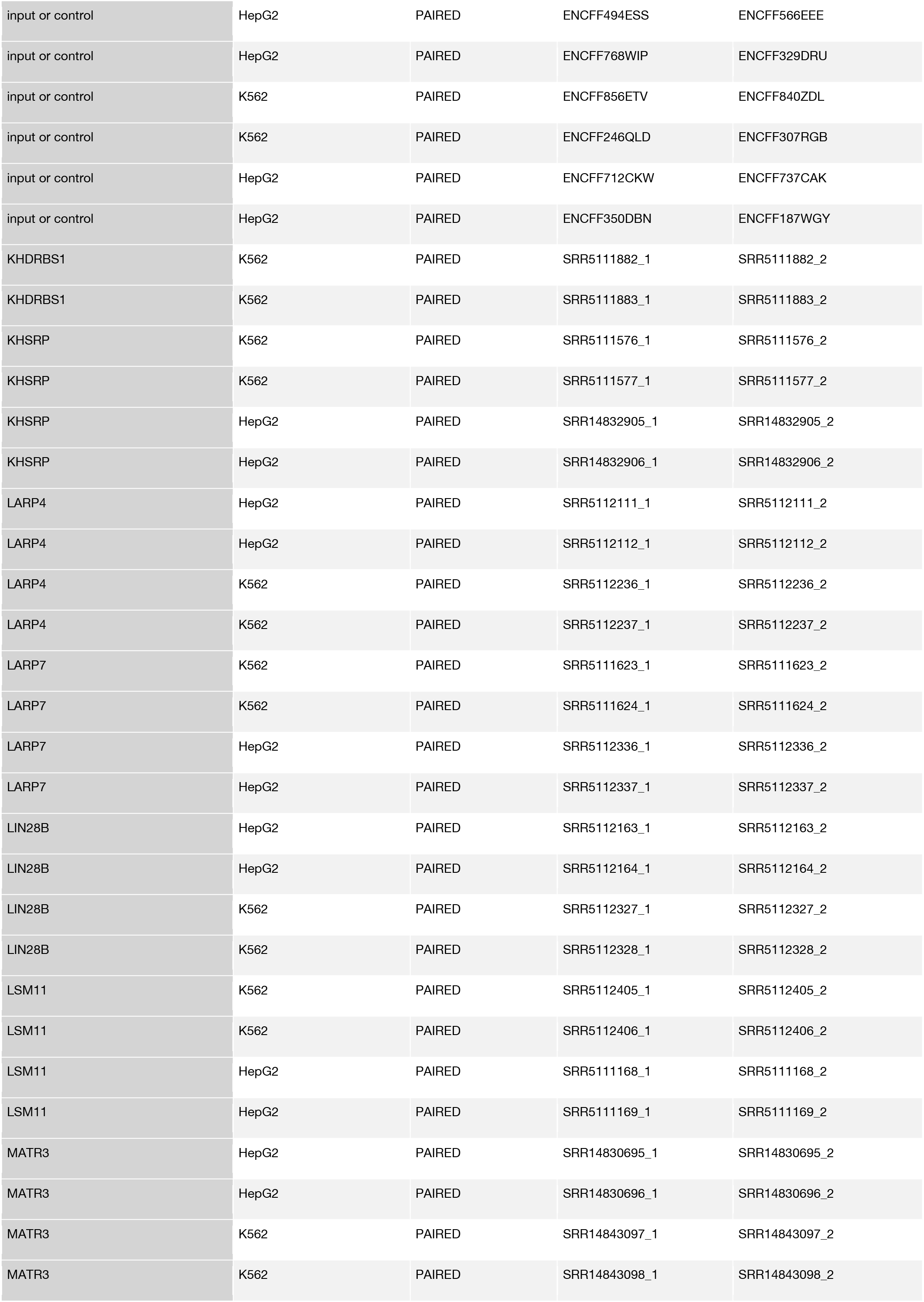
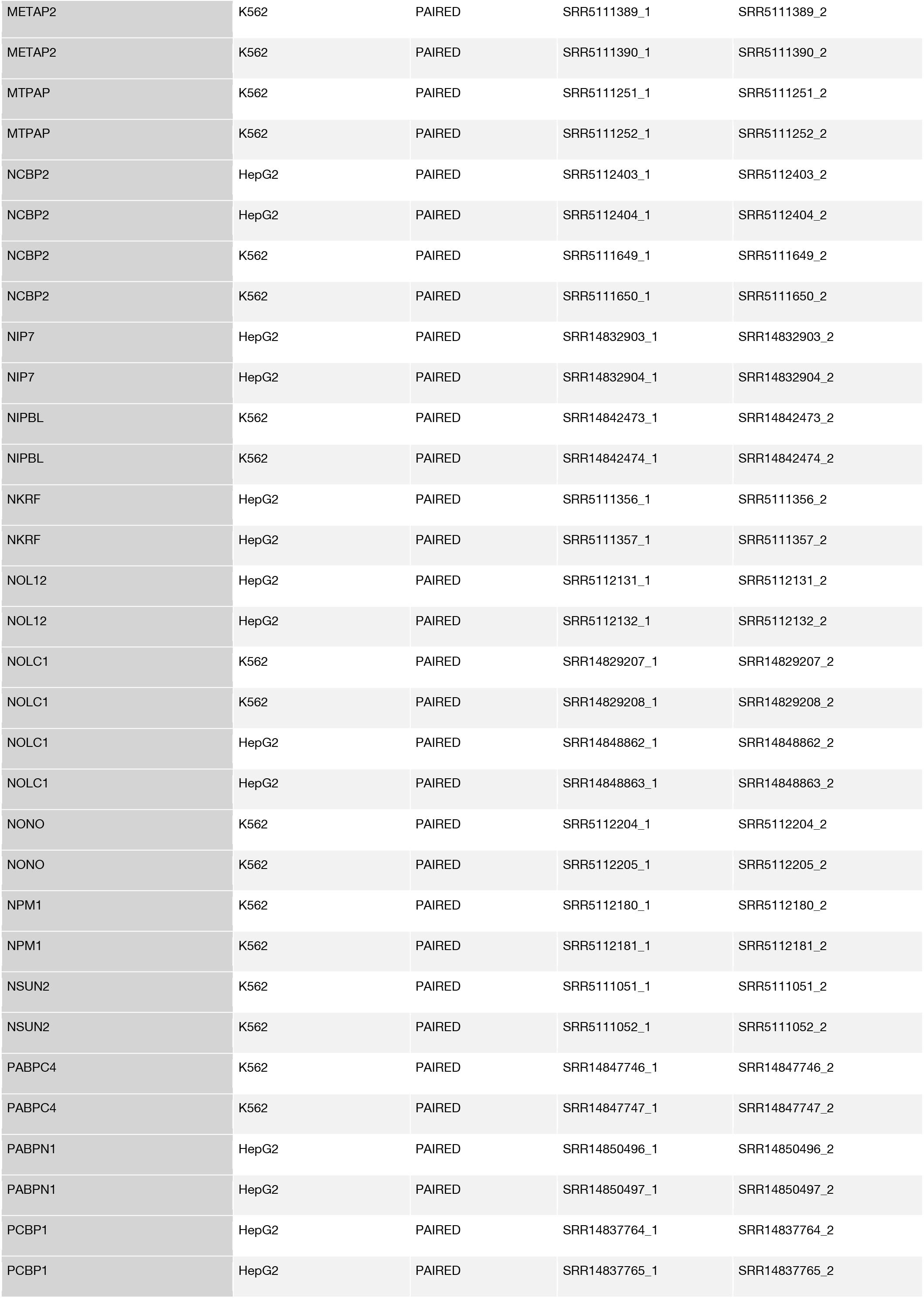
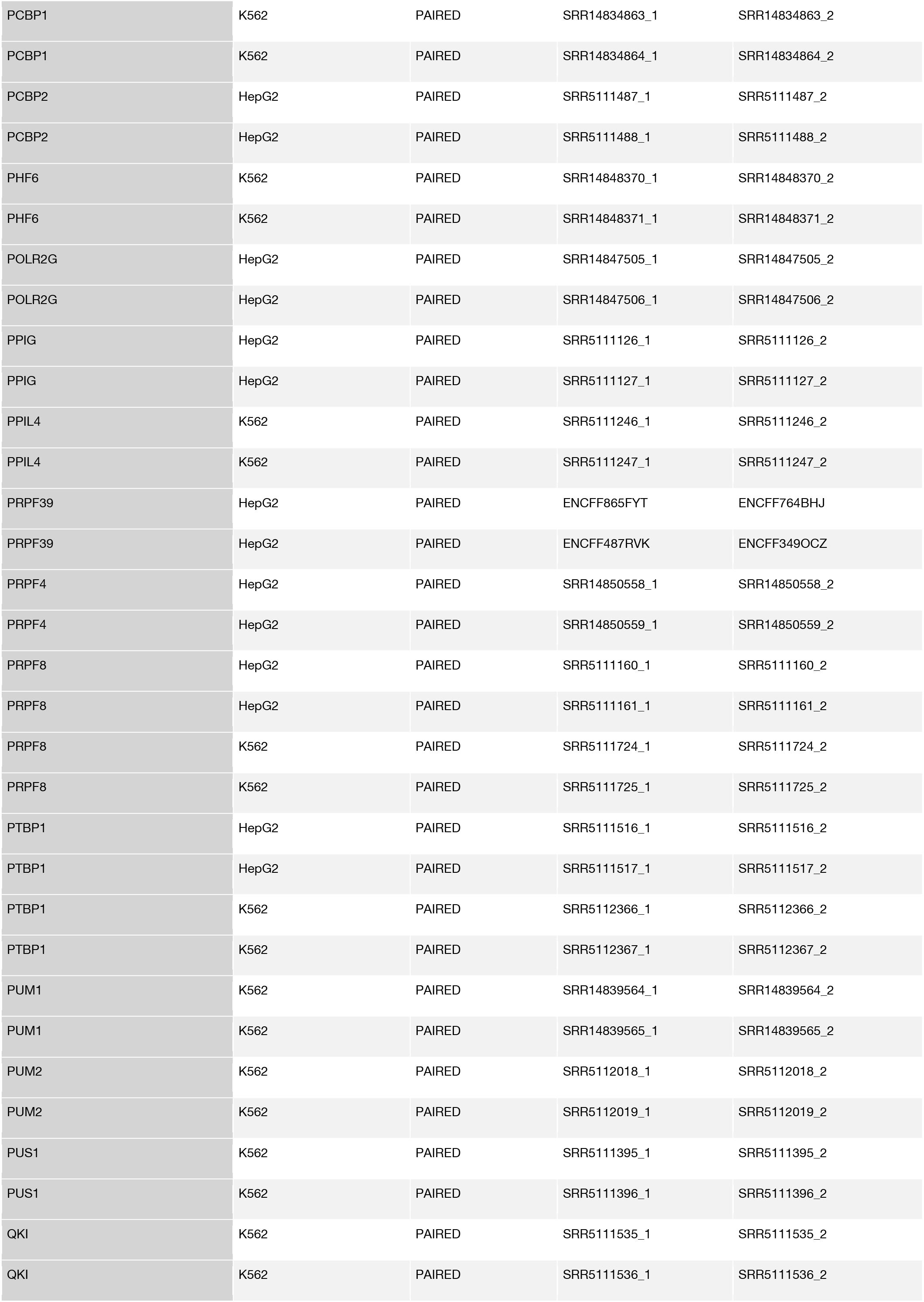
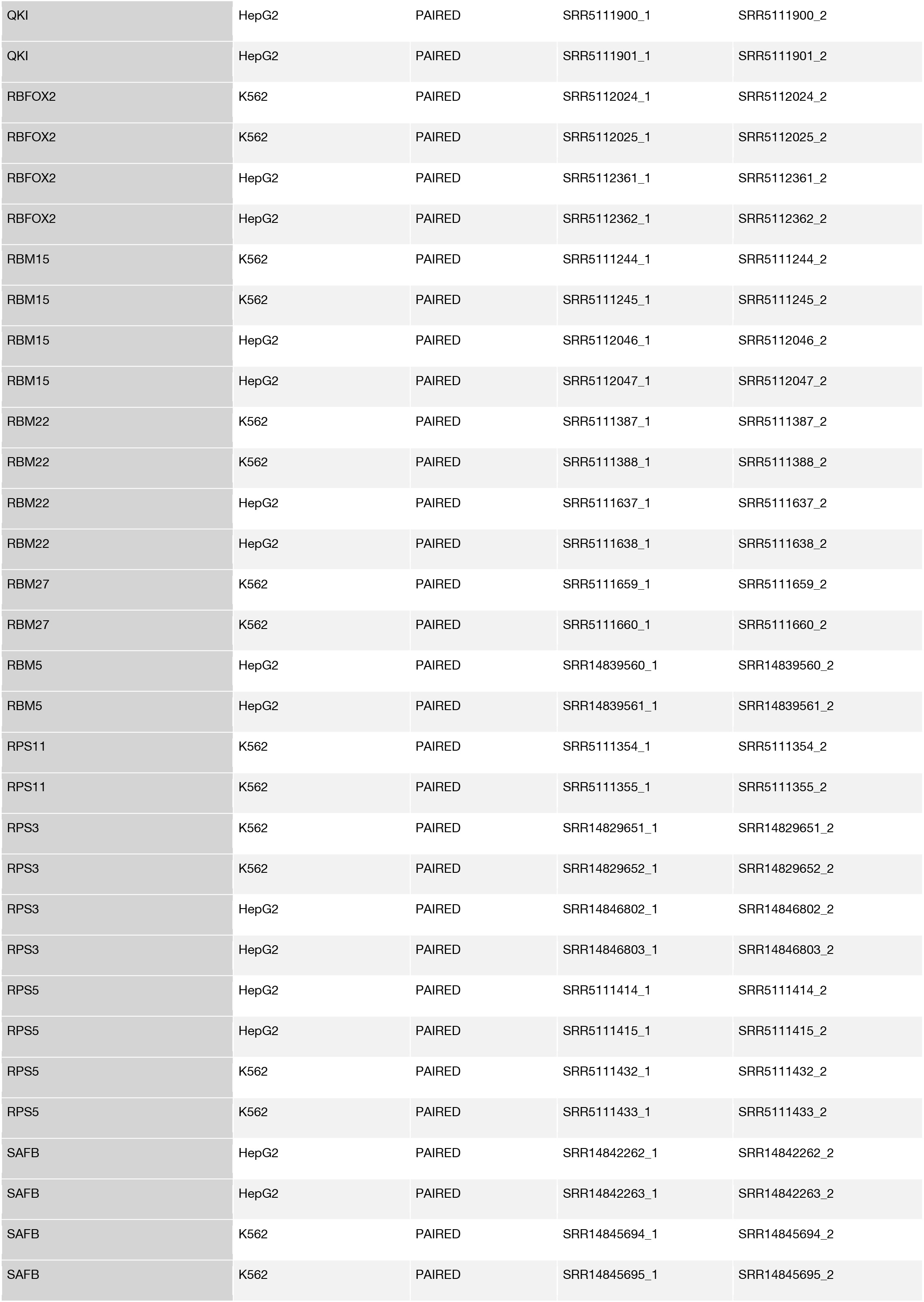
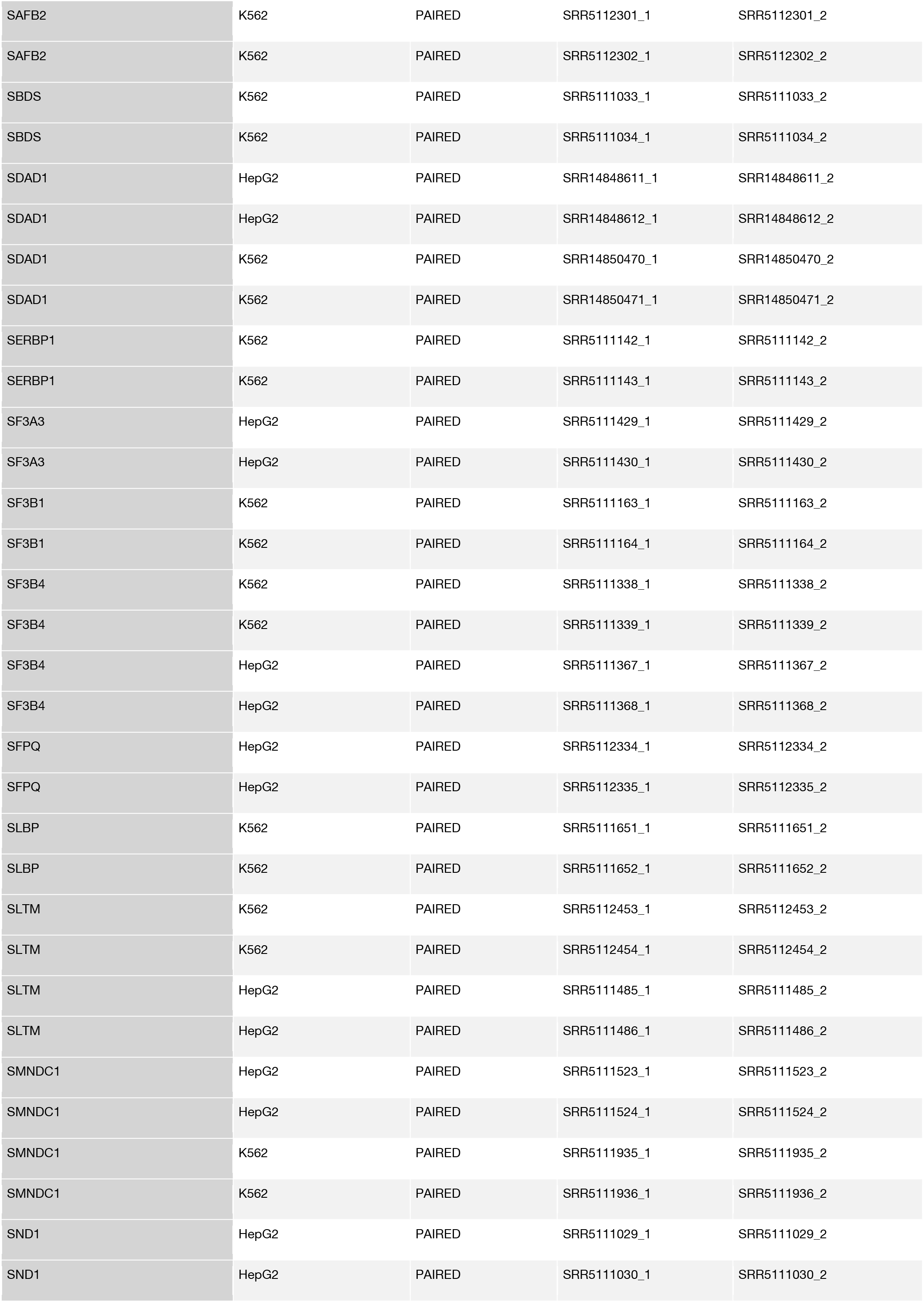
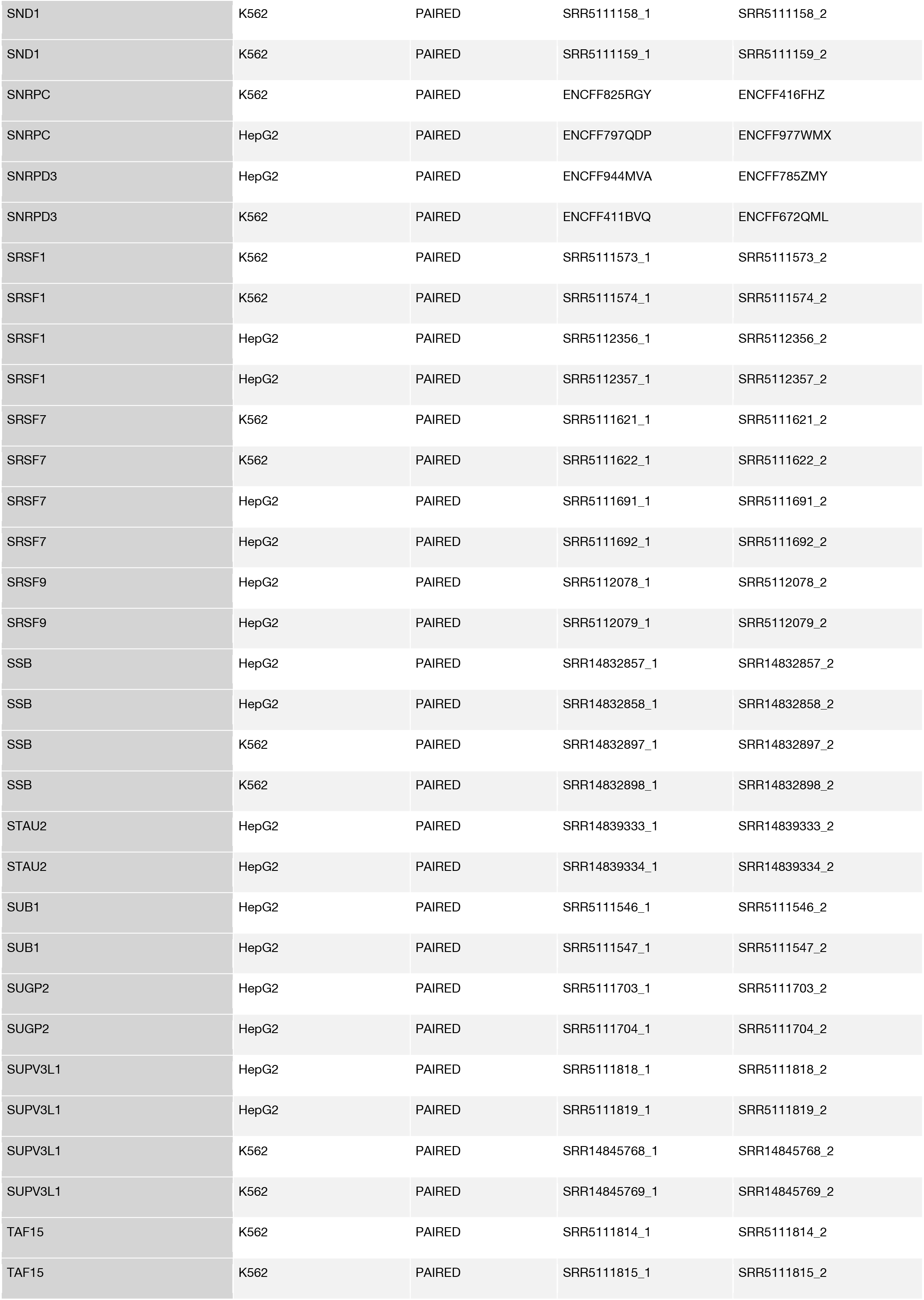
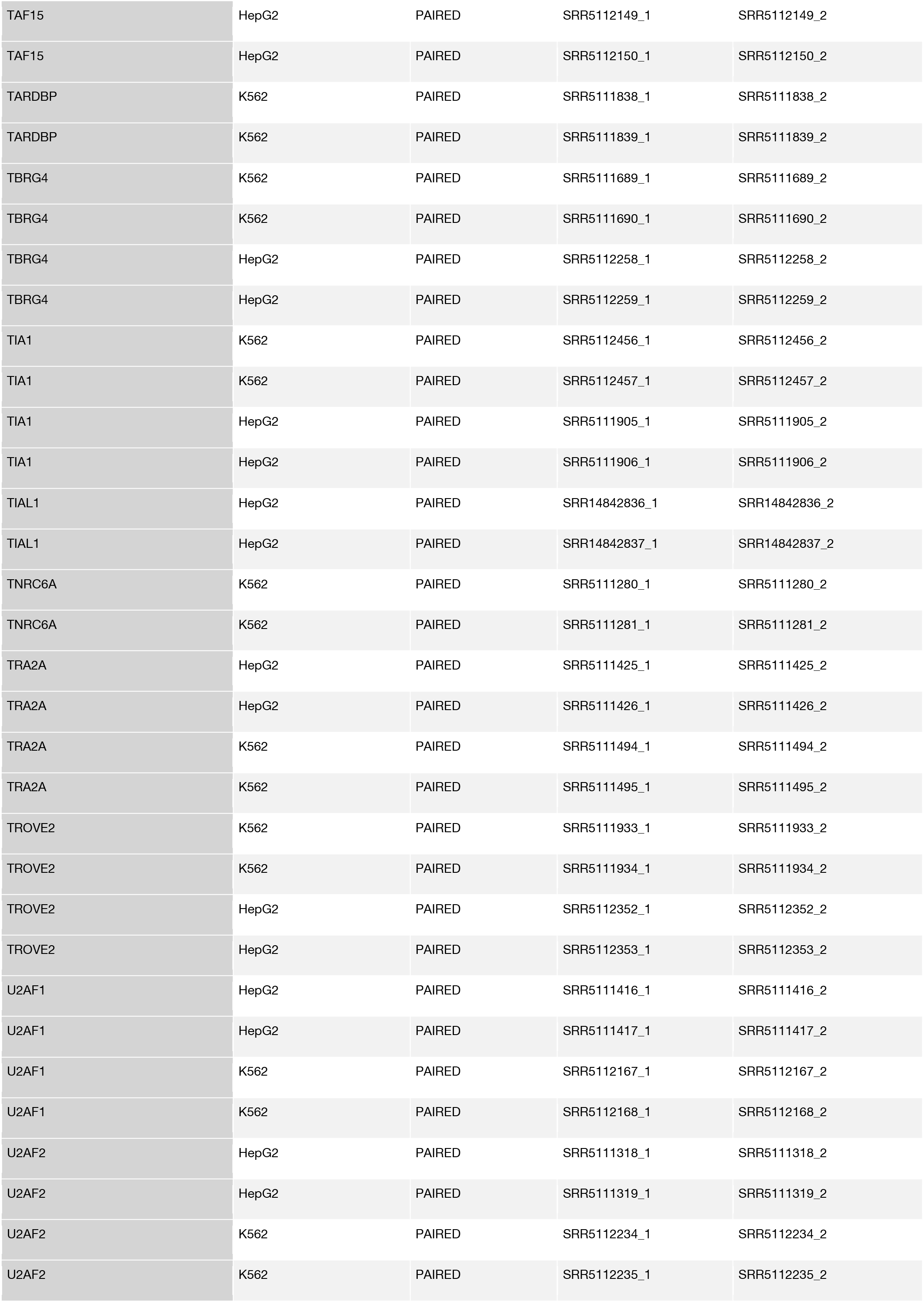
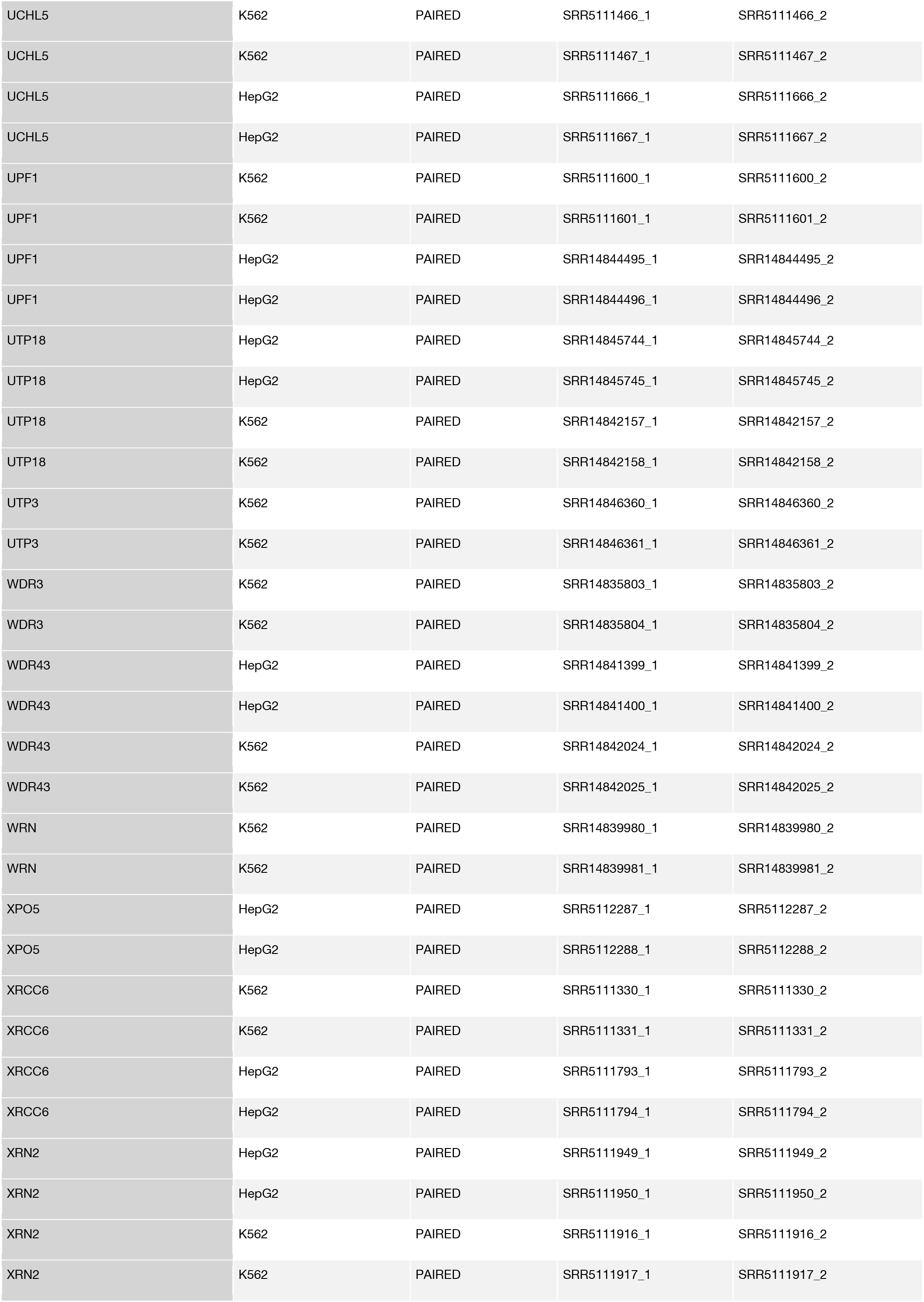
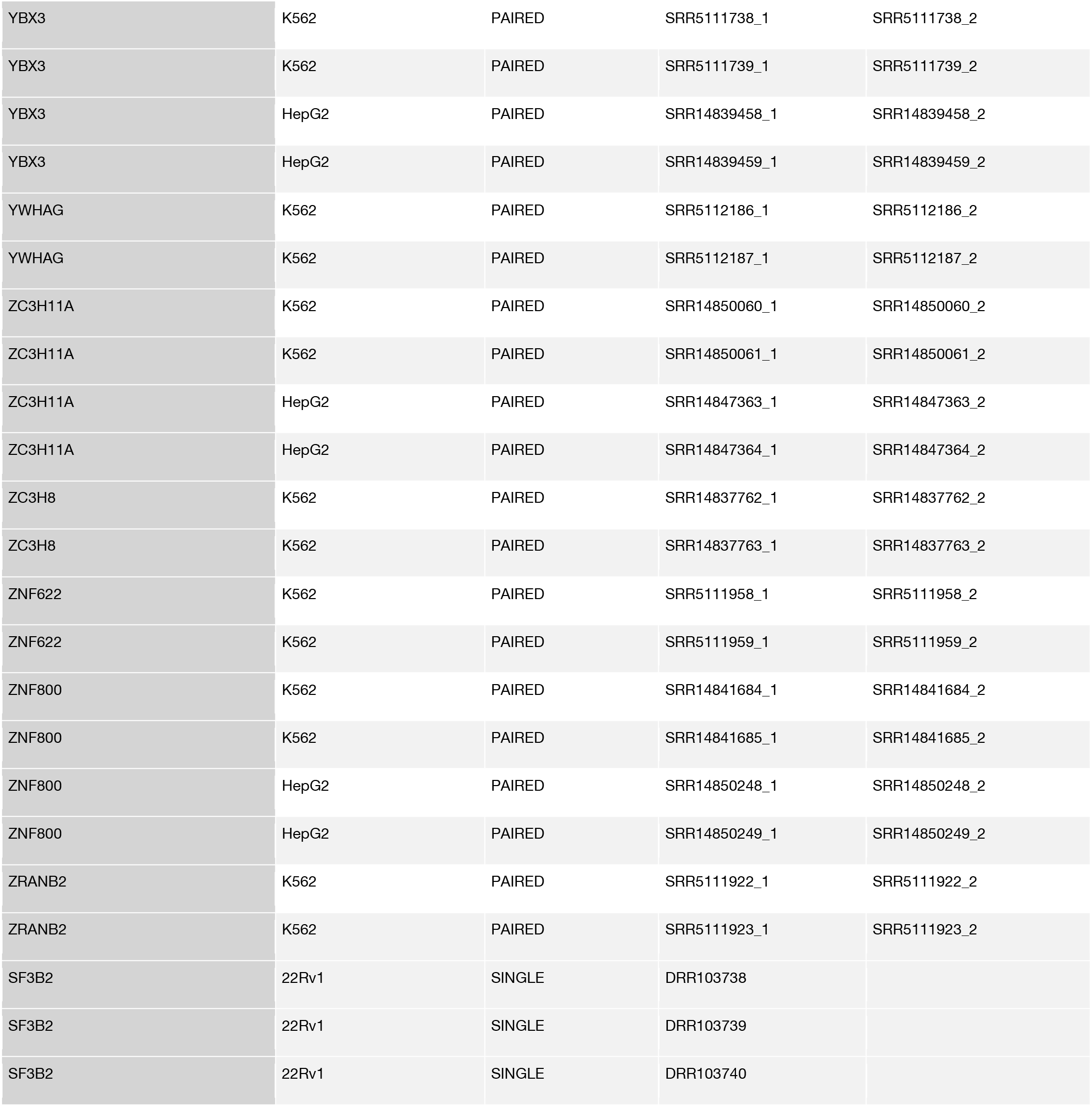
CLIP, eCLIP, and PAR-CLIP sequencing data used in this study.

